# Molecular mechanism of plasmid-borne resistance to sulfonamides

**DOI:** 10.1101/2022.06.30.498311

**Authors:** Meenakshi Venkatesan, Michael Fruci, Lou Ann Verellen, Tatiana Skarina, Nathalie Mesa, Peter J. Stogios, Alexei Savchenko

## Abstract

The sulfonamides (sulfas) are the oldest class of synthetic antibacterial that target the essential, conserved dihydropteroate synthase (DHPS) enzyme, encoded by *folP*, through chemical mimicry of its substrate *p*-aminobenzoic acid (*p*ABA). Resistance has complicated their clinical utility and is widespread in pathogenic species. Resistance is mediated by acquisition of *sul* genes on mobile genetic elements, which code for the so-called Sul enzymes that are divergent DHPS enzymes with intrinsic sulfa-insensitivity. Even decades after the discovery of this resistance mechanism, its molecular details have not been understood. In this study, we elucidate the molecular basis for intrinsic resistance of Sul enzymes using x-ray crystallography, enzymology, mutagenesis, intrinsic tryptophan fluorescence, antibiotic susceptibility of a contemporary Δ*folP* strain, and adaptive laboratory evolution of *folP.* We show that the active sites of Sul enzymes possess a modified *p*ABA-interaction region based on insertion of a Phe-Gly sequence. This insertion is necessary for discrimination between *p*ABA and sulfonamides, more than 1000-fold loss in binding affinity of sulfas to Sul enzymes, and robust *pan*-sulfonamide resistance. We detect no fitness cost due to this active site modification, as it does not compromise the rate of dihydropteroate biosynthesis and complements the thymidine-auxotrophy of an *E. coli folP* deletion strain. Lab-evolved sulfa-resistance *folP* recapitulated this mechanism through the same active site insertion. Finally, we show that this insertion and a nearby loop confer increased active site flexibility of Sul enzymes relative to DHPS. These results provide a molecular foundation for revisiting DHPS-targeted antibacterials to evade resistance.

## Introduction

Antimicrobial resistance (AMR) is a serious and growing problem in the management and prevention of infectious diseases. A recent analysis estimates that more than 1 million deaths were directly caused by and nearly 5 million deaths were associated with antimicrobial-resistant bacterial infections in 2019 (Murray et al., 2022). AMR is caused by multiple mechanisms including mutations in the genes coding for antimicrobial drug targets, enzyme-mediated drug direct modification or modification of the drug target, the action of drug efflux pumps, or acquisition of drug-insensitive alternatives to the drug target itself (Blair et al., 2015). Antimicrobial resistance-conferring genes (ARGs) often concentrate on mobile genetic elements (MGEs) encoding multiple ARGs, thereby conferring multidrug resistance and exchanging across species even across vast evolutionary distances in the bacterial kingdom (Partridge et al., 2018). Exacerbating the problem, the rate of discovery of truly new antimicrobial drug classes has crashed since the golden era of antimicrobial drug discovery in the mid-twentieth century, causes of which have been well-discussed (Beyer and Paulin, 2020; Butler et al., 2022; Lewis, 2020). New antimicrobial treatment options, including revitalizing old classes of drugs that have fallen out of favor, are urgently needed.

The sulfonamides (sulfas) were the first commercially available synthetic antibiotics. They were discovered in the 1930s and possess broad-spectrum bacteriostatic activity against both Gram-positive and Gram-negative bacteria; these compounds are used in both human and veterinary medicine world-wide (Lees et al., 2021; Ovung and Bhattacharyya, 2021; Sköld, 2014). Sulfas are used for the treatment of pneumonia (co-trimoxazole), bacterial meningitis, urinary tract infections (UTI) (sulphamethazine), bronchitis, ear infections (co-trimoxazole or sulphatrim), Crohn’s disease (sulphadiazine), traveller’s diarrhoea (co-trimoxazole) and topical bacterial infections (silver sulphadiazine) (Bryskier, 2005). As is the case for many classes of antimicrobials, resistance to sulfas complicates their clinical utility, and their usage has been diminished due to the prevalence of resistance (Huovinen, 2001; Vázquez-López et al., 2020): for example, the WHO reported that *Escherichia coli* and *Klebsiella pneumoniae* that cause UTIs possess high degrees of resistance (54% and 43%, respectively) to co-trimoxazole (“Global Antimicrobial Resistance and Use Surveillance System (GLASS) Report: 2021,” n.d.). With the rise in resistance to other classes of antibiotics that represent front-line treatment regiments, such as carbapenem and fluoroquinolones (Codjoe and Donkor, 2017; Drlica et al., 2019), classical antibiotics including sulfas are being re-evaluated for their priority in clinical practice (Cassir et al., 2014; Kaye et al., 2017; Theuretzbacher et al., 2015; Zayyad et al., 2017). In this respect, a detailed understanding of the molecular basis of sulfa resistance is essential in efforts to extend or revitalize the utility of this class.

Sulfonamides target dihydropteroate synthase (DHPS) encoded by *folP* gene and conserved throughout bacterial species. DHPS is essential for bacterial viability due to its involvement in folate synthesis, a pathway that is present only in bacteria and primitive eukaryotes. DHPS catalyzes the condensation of *para*-aminobenzoic acid (*p*ABA) and 6-hydroxymethyl-7,8-dihydropterin pyrophosphate (DHPP), producing 7,8-dihydropteroic acid (7,8-DHP) (**Figure 1a**); this compound is further transformed in the pathway, ultimately leading to tetrahydrofolate, an essential precursor for DNA and RNA synthesis. Sulfonamides structurally resemble *p*ABA (**Figure 1, Supporting Information Figure S1 for structures of sulfonamides**) and thus their mode of action is to directly compete with *p*ABA for binding to DHPS; condensation of DHPP and sulfas may also occur, forming a dead-end pterin-sulpha adduct (Swedberg et al., 1979; Yun et al., 2012; Zhao et al., 2016).

**Figure 1.**
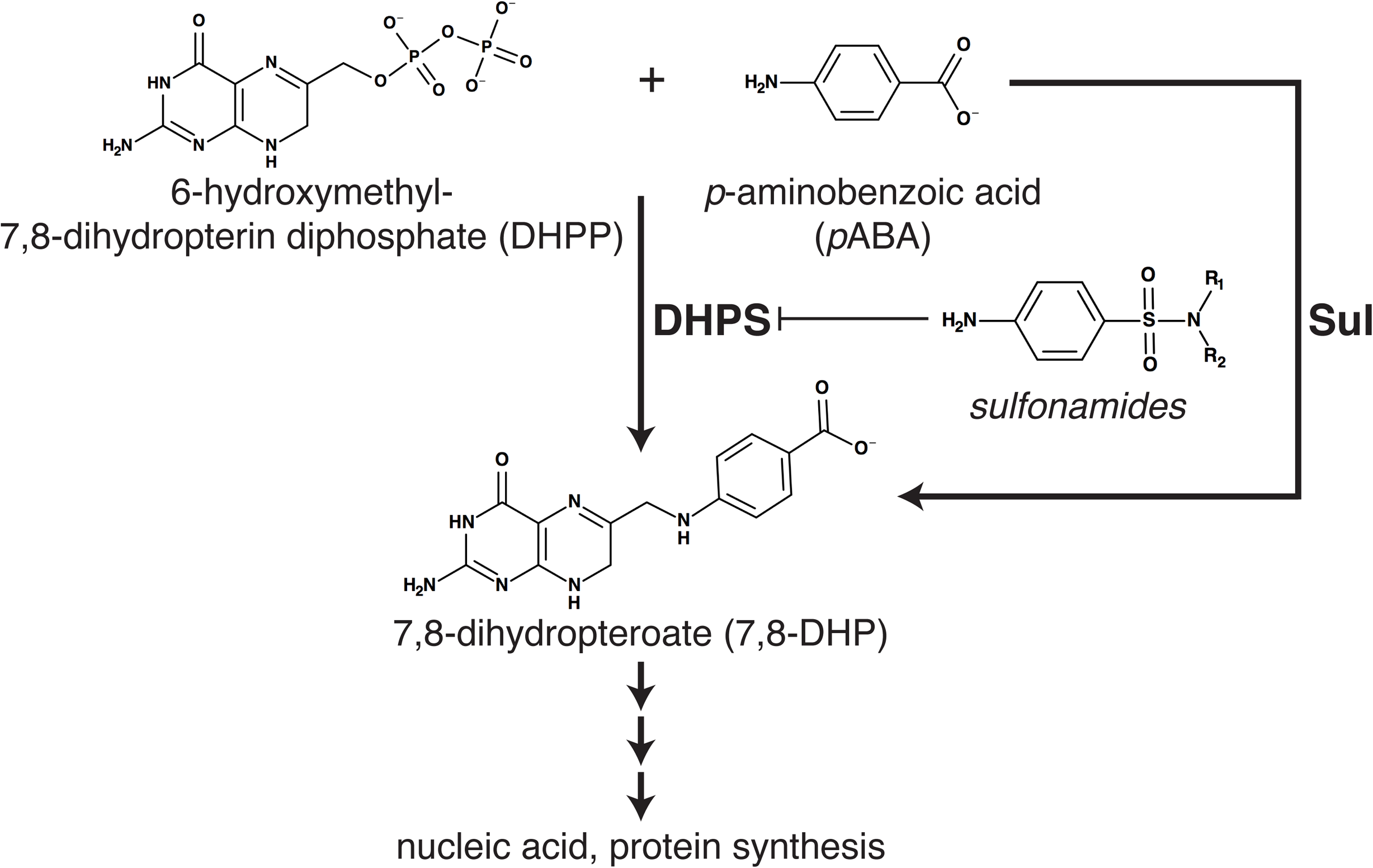
Schematic of folate pathway and chemical structures of sulfonamides and DHPS/Sul ligands.

DHPS enzymes adopt the classical (α/β)_8_ triose phosphate isomerase (TIM) barrel fold which positions the *p*ABA and DHPP binding sites within the central cavity, with two loops actively involved in the catalytic cycle. As described for DHPS from *Yersinia pestis* (*Yp*DHPS) and *Bacillus anthracis* (*Ba*DHPS), these loops (loops 1 and 2) are involved in pyrophosphate binding, coordination of a Mg^2+^ ion which facilitates release of the pyrophosphate leaving group and formation of a substructure formed by the two loops, which forms the *p*ABA binding site (Yun et al., 2012). The conformation of the loop 1-loop 2 substructure and the *p*ABA binding site are highly dependent on the presence of pyrophosphate within in the active site, which hindered analysis of this region of the active site until the crystal structure of the near-transition state complex of *Yp*DHPS was determined (Yun et al., 2012). The structural data on *Yp*DHPS confirmed the enzyme operates via a S_N_1 reaction mechanism, whereby DHPP binds to the active site first, then the pyrophosphate is eliminated, leaving a carbocation form of the pterin (DHP^+^) that reacts with the weakly nucleophilic *p*ABA amine group to form 7,8-dihydropteroate (Yun et al., 2012).

The resistance to sulfonamides occurs by two distinct mechanisms: through acquisition of mutations in *folP* and/or horizontal gene transfer of foreign, highly sequence-divergent genes coding for DHPS variants that are sulfa-insensitive. Through structural analyses, the molecular basis of resistance through mutations in *folP* is well understood: resistance-conferring amino acid substitutions in DHPS have been mapped to residues in loops 1 and 2 that line the *p*ABA/sulfonamide binding site in several bacterial species (Baca et al., 2000; Griffith et al., 2018; Yun et al., 2012). These mutations have the effect of increasing the K_M_ parameters for sulfonamides with less dramatic effects of the K_M_ for *p*ABA, thus conferring a substrate discrimination capacity to the DHPS enzymes that the WT enzymes lack (Griffith et al., 2018).

Sulfonamide resistance genes encoded on plasmids are frequently reported in clinical isolates in Gram-negative pathogenic species including *Escherichia coli*, *Acinetobacter baumannii*, and *Klebsiella pneumoniae* (Carvalho et al., 2021; Li et al., 2021; Liu et al., 2020; Sköld, 2000). Transferable sulfonamide resistance was first noted in the 1950s and 1960s between *Shigella* and *E. coli* using *in vitro* and *in vivo* models (Akiba et al., 1960; Kasuya, 1964) however, this transferable resistance was not identified and characterized until 1975 and later. To date, four mobilizable *sul* genes have been detected. *sul1* was initially discovered in 1975 in *E. coli* and *Citrobacter* sp. (Skold, 1976; Wise and Abou Donia, 1975), *sul2* discovered in 1980 in *E. coli* causing UTIs (Radstrom and Swedberg, 1988; Swedberg and Skold, 1983, 1980), *sul3* was identified in *E. coli* from pigs in 2003 (Grape et al., 2003; Perreten and Boerlin, 2003) and *sul4* has recently been discovered from an unknown bacterium in wastewater (Razavi et al., 2017). According to the CARD Database which tracks AMR elements (Alcock et al., 2020) indicates that 40% of *Pseudomonas aeruginosa*, 16% *Enterobacter cloaceae*, 18% *Klebsiella pneumoniae* and 44% of *Actinetobacter baumannii* publicly available genomes contain the *sul1* gene, reflecting its high degree of spread. Similarly, 4-7% of plasmids in these species harbor the *sul2* gene. The *sul3* gene is less prevalent, present only less than 1% of plasmids in these species. The *sul* genes are often part of multiple resistance gene clusters (Carvalho et al., 2021; Li et al., 2021; Liu et al., 2020; Mori et al., 2021; Pavelquesi et al., 2021; Sun et al., 2021). *Sul* genes are disseminated to much higher degree in some species, with the CARD database indicating the prevalence of *sul1* as high as 50% in plasmids in *Comamona testosteroni*, an environmental species that rarely causes infections (Farooq et al., 2017) and *sul2* as high as 45% in *Acinetobacter towneri*, a species known to serve as reservoir of ARGs (Zou et al., 2017)*. suls* are highly dispersed in the environment and in fact, *sul1* is used as a surrogate marker for the dissemination of ARGs and tracking anthropogenic influence on the environment i.e. waste-water treatment and decontamination of ARGs (Pazda et al., 2019; Pruden et al., 2012; Wang et al., 2020; Zhuang et al., 2021).

The origin(s) of the *sul* genes has been the subject of recent research which suggested that they evolved from lateral transfer of chromosomal *folP* genes of *Rhodobiaceae* and *Leptospiraceae* species (Sánchez-Osuna et al., 2019). However, the direct ancestor of *sul* genes has not been identified or the mechanism of their capture onto MGEs. Sánchez-Osuna *et* al (Sánchez-Osuna et al., 2019) also identified a two amino-acid “Sul motif” that is distinctive of the Sul enzymes relative to DHPS (a Phe-Leu sequence) and confirmed that *folP* enzymes from *P. lavamentivorans* DS-1 and *L. interrogans* that harbor this motif confer sulfa resistance when expressed in *E. coli*. However, the molecular role of this motif, and nearby amino acids, for resistance was not determined. Consistent with the notion that *sul* genes originated from evolution of *folP* within environmental microbiota, two other studies using functional metagenomics sampling of soil or agricultural environments identified divergent *folP* genes that confer sulfa resistance (Lau et al., 2017; Willms et al., 2019). Interestingly, these DHPS enzymes also harbor the “Sul motif” but its molecular role and relevance to resistance was not studied.

While sulfas inhibit the chromosomally-encoded DHPS enzyme, Sul enzymes possess the ability to distinguish the natural substrate *p*ABA from sulfonamides and thus acquisition of Sul enzymes restores this step in the folate synthesis pathway. The first observation of the ability of Sul enzymes to distinguish between *p*ABA and sulfonamides occurred from a comparison of DHPS activity from sulfa-resistant *E. coli* vs sulfa-sensitive *E. coli* (Swedberg and Skold, 1980). Approximately 10000-fold more sulfonamide was required to inhibit the activity of a crude purification of Sul1 as compared to DHPS (Swedberg and Skold, 1980). Sul enzymes exhibit on average ∼30% sequence identity to the *E. coli* DHPS enzyme (*Ec*DHPS). There have been no reported structure-function studies on Suls elucidating the molecular basis for resistance and thus the molecular basis by which the Sul enzymes confer resistance is not well understood.

Here, we use x-ray crystallography, *in vitro* activity characterization, intrinsic tryptophan fluorescence, heterologous expression in a contemporary *E. coli* Δ*folP* strain and adaptive laboratory evolution of *folP* to reveal the molecular basis of sulfa resistance by the plasmid-encoded Sul1, Sul2, and Sul3 enzymes (herein, collectively referred to as Sul enzymes). While these Sul enzymes largely resemble DHPS enzymes in overall structure, they possess a remodeled *p*ABA-binding region and different conformational dynamics of their active sites as compared to DHPS enzymes. Sul enzymes possess an insertion of a phenylalanine residue positioned to block sulfonamide binding, and we show this residue is necessary for sulfonamide resistance.

## Results

### Despite sequence divergence the Sul enzymes demonstrate catalytic properties similar to those of DHPS

To gain an understanding of the *sul* incurred resistance mechanism we first characterize the catalytic properties of the Sul enzymes in comparison to DHPS. We purified the Sul1, Sul2 and Sul3 enzymes recombinantly expressed in *E. coli* along with DHPS from the same species (*Ec*DHPS) as an archetype for *folP* encoded enzymes. Since the substrate DHPP is not commercially available, we leveraged and modified a previously published synthesis protocol (Hevener et al., 2010) to obtain this substrate by full product conversion (see Experimental for more details) from 6-hydroxymethyl-7,8-dihydropterin (6-HMP) using recombinantly purified 6-hydromethyl-7,8-dihydropterin pyrophosphokinase (*Ec*HPPK) (Blaszczyk et al., 2004). We then characterized the activity of these four enzymes *in vitro* by measuring the release of pyrophosphate (PP_i_) by the Malachite Green assay (see the details in Materials and Methods). In the presence of excess *p*ABA (200 µM), the saturation of the second co-substrate (DHPP) could not be reached and thus we were not able to obtain the kinetic parameters for this substrate under the used experimental conditions. This phenomenon, previously reported for several bacterial DHPS enzymes is attributed to the S_N_1 ordered bi-substrate reaction mechanism followed by this family of enzymes (Hammoudeh et al., 2014). Accordingly, the DHPP is always the leading substrate for initiating the reaction and promotes accommodation of *p*ABA in the active site (Yun et al., 2012). Thus, we proceeded to characterization of Sul and *Ec*DHPS enzymes kinetic properties with respect to the *p*ABA co-substrate in the presence of excess of DHPP (200 µM). All three Sul enzymes demonstrated similar enzymatic properties reflected in close values of K_M_ (8.4-9.7 µM) and k_cat_ (0.34-0.46 s^-1^) for *p*ABA, which were also comparable to those for *Ec*DHPS (7.8 µM and 0.38 s^-1^, respectively) (**Table 1 and Supporting Information Figure S2**). Accordingly, the three Sul and *Ec*DHPS enzymes showed no significant difference in catalytic efficiency (k_cat_/K_M_) for the *p*ABA co-substrate (**Table 1**). These results suggested that any representative of the Sul enzymes can effectively replace the activity of *folP* encoded endogenous enzyme in folate biosynthesis in *E. coli*.

**Table 1 -.** Kinetic parameters for *p*ABA and SMX of Sul and *Ec*DHPS enzymes.

| Enzyme | Native substrate (*pABA*) | | | Sulfa (SMX) | | | | Fold change<br>$k_{\text{cat}}/K_M$ |
| --- | --- | --- | --- | --- | --- | --- | --- | --- |
| | $K_M$ (mM) | $k_{\text{cat}}$ ($\text{s}^{-1}$) | $k_{\text{cat}}/K_M$ ($\text{s}^{-1} \text{mM}^{-1}$) | $K_i$ (mM) | $K_M$ (mM) | $k_{\text{cat}}$ ($\text{s}^{-1}$) | $k_{\text{cat}}/K_M$ ($\text{s}^{-1} \text{mM}^{-1}$) | *pABA* vs.<br>SMX |
| **Sul1<sup>WT</sup>** | 8.41±1.4 | 0.38 | 44.69 | 735±89 | 1004±77 | 0.26 | 0.26 | 170.6 |
| **Sul2<sup>WT</sup>** | 9.63±1.2 | 0.46 | 48.10 | 609±140 | 800±58 | 0.28 | 0.35 | 139.4 |
| **Sul3<sup>WT</sup>** | 9.66±0.9 | 0.42 | 43.33 | 373.6±77 | 636±67 | 0.25 | 0.39 | 112.2 |
| **Sul1<sup>F178G</sup>** | 9.80±1.2 | 0.34 | 34.75 | 3.03±0.3 | 9.4±0.9 | 0.51 | 53.6 | 0.93 |
| **Sul3<sup>F177G</sup>** | 9.49±1.1 | 0.47 | 49.09 | 4.16±0.3 | 9.3±0.9 | 0.49 | 52.75 | 0.65 |
| ***EcDHPS*** | 7.82±0.6 | 0.38 | 48.59 | 5.15±1.1 | 7.67±0.7 | 0.46 | 59.4 | 0.82 |

*pABA* and SMX kinetic parameters of Sul and *EcDHPS* enzymes. The substrate turnover $k_{\text{cat}}$ and catalytic efficiency $k_{\text{cat}}/K_M$ was calculated from the $V_{\text{max}}$ and $K_M$ obtained from the Michaelis-Menten non-linear regression using GraphPad Prism v5.0. Error represents SD for the mean value (mean of biological replicates of $n=3$ ) for each *pABA*/SMX concentration.

### Sulfa drugs fail to inhibit the Sul enzymes

In line with our kinetic analysis, a previous study reported similar affinity of Sul and *Ec*DHPS enzymes toward the *p*ABA co-substrate while demonstrating that sulfathiazole was 10000-fold less effective in abrogating the activity of Sul enzyme compared to *Ec*DHPS (Swedberg and Skold, 1980). We therefore sought to determine whether sulfa drugs can be metabolized in the dihydropteroate synthase reaction catalyzed by Suls. In a control experiment we confirmed the *Ec*DHPS inhibition by sulfamethoxazole (SMX) in the reaction leading to production of a dead-end pterin-sulfa adduct. By mass spectrometry, we validated formation of the pterin-SMX adduct catalyzed by *Ec*DHPS (**Supporting Information Figure S3**) when SMX is in the low µM range. According to our assay the activity of *Ec*DHPS was inhibited by SMX with a K_i_ of 5.1 µM. This value is similar to this enzyme’s K_M_ for *p*ABA (**Table 1 and Supporting Information Figure S3**), reflecting the inability to discriminate between the natural substrate and this sulfa compound. According to our assay the catalytic efficiency of *Ec*DHPS for SMX and *p*ABA is 49 and 59 s^-1^ mM^-1^, respectively, also reflecting similarity in chemical structure between *p*ABA and SMX (**Table 1 and Supporting Information Figure S3**).

In the presence of SMX and with DHPP in excess, the Sul enzymes activity also leads to the formation of pterin-SMX adduct by mass spectrometry. However, significantly higher concentrations of SMX are necessary for the formation of the adduct, indicating lower efficiency of Sul toward SMX as compared to *Ec*DHPS. More specifically, our kinetic analysis suggests that Sul1 demonstrates 131-fold less affinity to SMX, while Sul2 and Sul3 show 104 and 83-fold decrease in affinity, respectively (**Table 1**). Similarly, low affinity of Suls with respect to SMX is reflected in the catalytic efficiency values difference for SMX as aa substrate versus *p*ABA (between 111 to 137-fold lower). Likewise, the dihydropteroate synthase activity of Suls is inhibited by SMX with much lower K_i_ values (between 143 to 73-fold lower) than in case of *Ec*DHPS. Altogether, these results unambiguously confirm the inability of sulfa drugs such as SMX to inhibit Sul enzymes despite their enzymatic properties being similar to those of DHPS enzymes. This divergence appears to be based on ability of Sul enzymes to discriminate between *p*ABA and SMX compounds, favoring the former as a substrate.

### Crystal structures of Sul enzymes reveal active features responsible for ligand binding

To understand the molecular basis of resistance conferred by Sul enzymes since no structures were available at the time of this research, we pursued the structural characterization of the Sul1, Sul2 and Sul3 enzymes in apo form as well as in complex with ligands. We obtained diffraction-quality crystals of Sul1 in the presence of DHPP (2.32 Å), Mg^2+^ and *p*ABA; Sul2 in the *apo-*enzyme state (1.85 Å) and in the presence of 7,8-DHP and Mg^2+^(1.74 Å); Sul3 in the *apo*-enzyme state (2.80 Å), in the presence of 6-HMP (2.42 Å), and in the presence of DHPP, Mg^2+^ and *p*ABA (2.00 Å). We note that successful crystallization of the Sul2·7,8-DHP·Mg^2+^·PP_i_ and Sul3·6-MP·Mg^2+^·PP_i_ complexes, as described below, was obtained only using an incubation procedure where the Sul enzyme was first incubated with DHPP, followed by *p*ABA (see Experimental for more details). We used Molecular Replacement to solve these crystal structures using the structure of *Ec*DHPS (Achari et al., 1997); full x-ray crystallographic statistics are shown in **Table S1**.

According to our structural data, all three representative Sul enzymes adopted a common (ɑ/β)_8_ TIM barrel fold (**Figure 2a, Supporting Information Figure S4**) also shared by DHPS enzymes (**Figure 3**). The active site cleft harboring ligands is located on one face of the barrel. Comparison of our 6 crystal structures revealed conformational differences in three key active site loops (designated as loops 1 through 3 here), depending on the ligand binding state of the enzymes (**Figure 2a and Figure 2b**). These loops play key roles in interactions with ligands, as described below for Sul2 and Sul3. In particular, loop 1 adopts a conformation away from the ligand binding cleft and catalytic center in the absence of *p*ABA or 7,8-DHP, and collapses onto the catalytic center in the presence of these molecules (**Figure 2**). Loop 3 undergoes a similarly large conformational change upon binding of *p*ABA or 7,8-DHP (**Figure 2**). Loop 2 undergoes conformational changes in a more confined space, but it also reacts to the presence of the ligands (**Figure 2**). In the Sul2·7,8-DHP·Mg^2+^·PP_i_ complex and Sul3·6-MP·Mg^2+^·PP_i_ complexes, details of which are described below, loops 1 through 3 adopt the same conformation which we interpret as the catalytically-competent conformation of the Sul enzymes.

**Figure 2.**
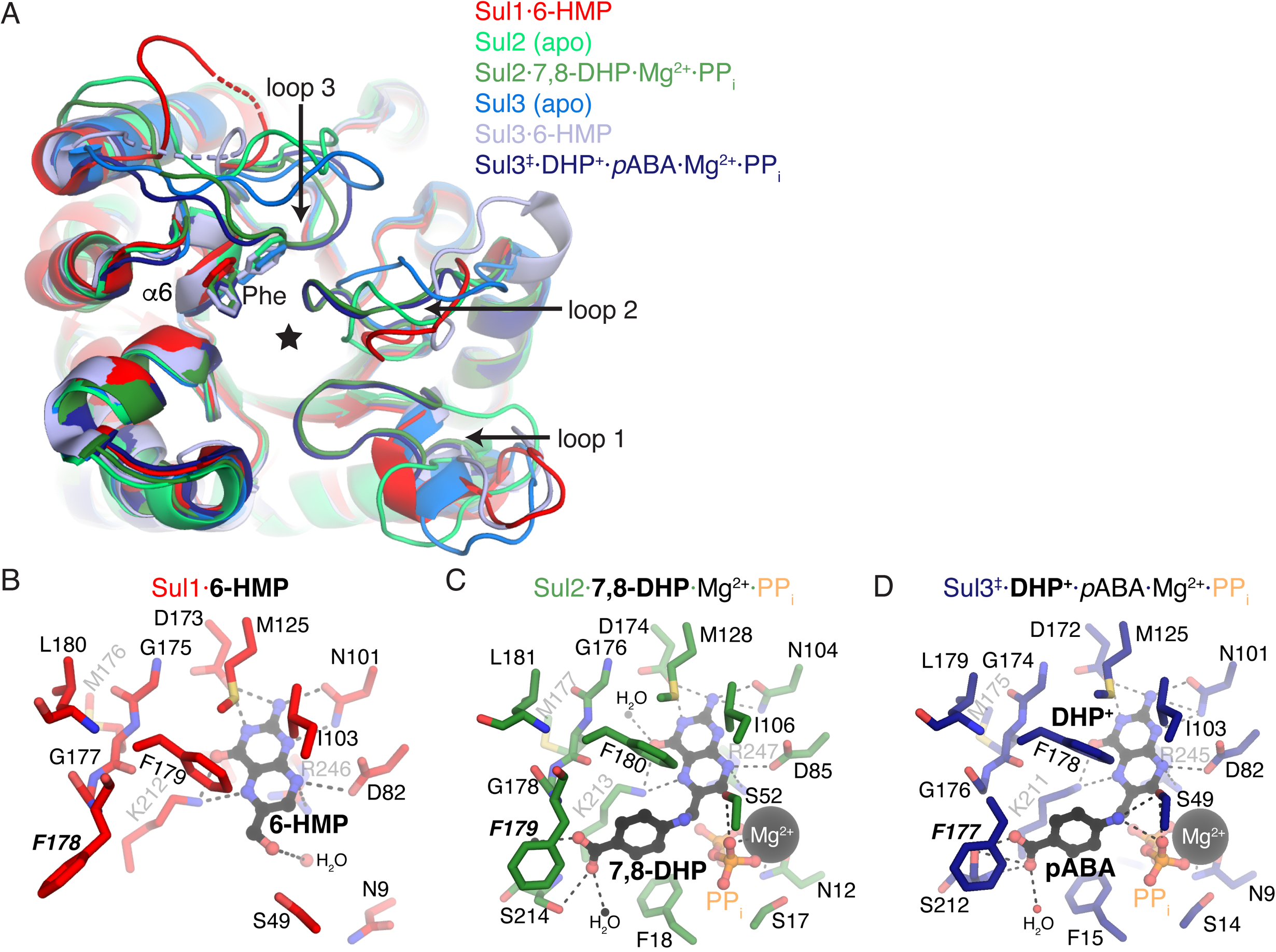
Structures of Sul enzymes. A) Superposition of Sul enzyme structures determined in this study. Loops 1 through 3 are labeled. Active site Phe that confers sulfonamide resistance is shown in stick representation and labeled. Star indicates location of catalytic site. B through D) Catalytic sites of Sul enzymes. Bound ligands are shown in black ball-and-stick, magnesium ions in spheres. Active site Phe that confers sulfonamide resistance is labeled in bold italics.

**Figure 3.**
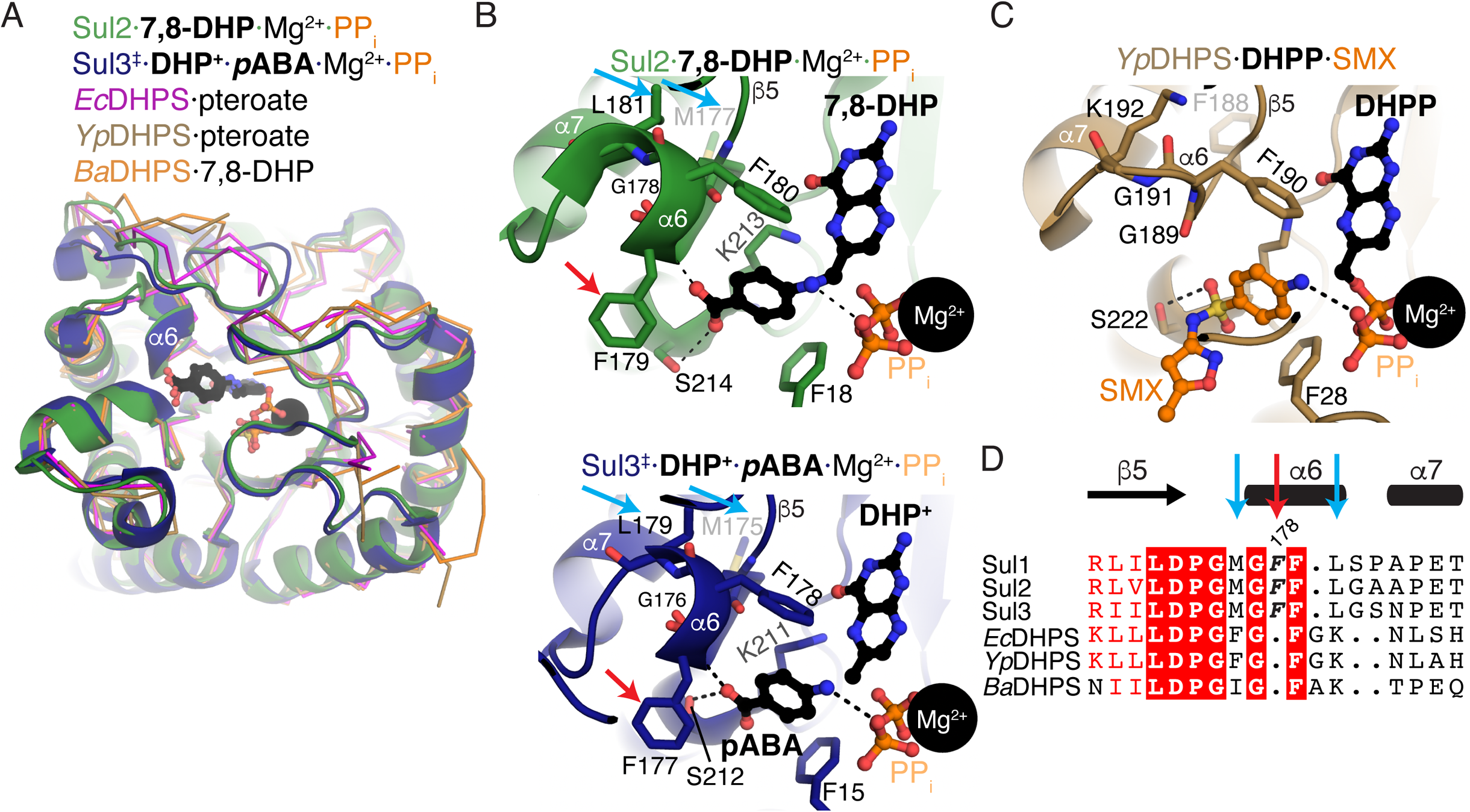
Comparison of structures of Sul and FolP enzymes. A) Superposition of Sul2·7,8-DHP·Mg^2+^·PP_i_ complex, Sul3·6-MP·Mg^2+^·PP_i_ complex, CbFolP·pteroate complex (PDB 3tr9), BaFolP·7,8-DHP complex (PDB 3tya), YpFolP·pteroate complex (PDB 3tyu). B) Focus on *p*ABA/sulfonamide interacting areas of catalytic sites. Amino acids shown in sticks are those that interact with *p*ABA and/or sulfamethoxazole (in the case of panel with YpFolP·DHPP·SMX complex, PDB 3tzf). For both panels, ligands are shown in ball- and-stick and Mg^2+^ ions are shown as black spheres. C) Structure-based multiple sequence alignment of Sul and FolP enzymes. Shading represents increasing sequence conservation. Secondary structure elements shown above alignment. Red arrow and bold+italics representation indicates Phe insertion in Sul enzymes and blue arrow indicates substitutions to hydrophobic residues nearby the Phe insertion (Met and Leu) in Sul enzymes. Residue F178 refers to Sul1 numbering.

We also noticed that an active site Phe residue in each of the Sul enzymes (Sul1 Phe178, Sul2 Phe179, Sul3 Phe177) adopted different sidechain rotamers depending on the ligand binding state (**Figure 2**). In the *apo*-enzyme states of Sul2 and Sul3, the sidechain is positioned towards loop 3 as this loop itself is positioned far from the catalytic center. In any of the ligand-bound structures, this residue adopts a conformation away from loop 3 and towards the *p*ABA binding site, which will be described in detail in subsequent sections.

Inspection of the active site of the Sul1 crystal structure revealed electron density that we modeled as 6-HMP (**Supporting Information Figure S4A**). There was some evidence of additional density beyond the 6-HMP suggestive of a phosphodiester oxygen, however, this additional density could not be confidently modeled due to the presence of loop 2 residue Ser49. In fact, loops 1 and 2 blocked the Sul1 active site across the dimerization interface, suggesting crystallization induced what we interpret as a non-functional conformation of these loops (**Supporting Information Figure S4A**). Nonetheless, the 6-HMP molecule formed multiple interactions with Sul1, including with Asn101, Asp173 and Lys212, each of which formed multiple hydrogen bonds with the pterin ring. As well, the plane of the pterin ring formed van der Waals interactions with Phe179 and Ile103.

The crystal of Sul2 grown in the presence of 7,8-DHP and Mg^2+^ showed evidence of these ligands and PP_i_; even though this crystal was grown under aerobic conditions, and our demonstration of sulfa-adduct formation results in oxidized product, inspection of the electron density supports the presence of 7,8-DHP and not the oxidized pteroic acid (**Supporting Information Figure S4**). This structure represents a fully organized active site, with each of loops 1, 2 and 3 organized in the active site, and reflects both products of the Sul enzymatic reaction before release of free PP_i_ (**Figure 2d**). The interactions observed between Sul1 and 6-HMP were also observed in the Sul2·7,8-DHP·Mg^2+^·PP_i_ complex, but the presence of Mg^2+^ and PP_i_ revealed additional interactions contributed by loops 1 residues Asn12, Ser17, Phe18 and the loop 2 residue Ser52, consistent with the role of these loops in PP_i_ release. In comparison, *p*ABA formed fewer interactions with the enzyme, with interactions between the *p*ABA carboxylate with Ser214, the *p*-amino group with Ser52, and hydrophobic interactions with aliphatic region of Lys213 and the sidechain of Phe179.

The crystal of Sul3 that grew in the presence of DHPP, Mg^2+^ and *p*ABA reflected a completely organized active site. The crystal contained clear electron density corresponding to *p*ABA, Mg^2+^ and PP_i_ (**Supporting Information Figure S4f**). The crystal showed weaker but noticeable electron density for the residual pterin ring that we interpret as the carbocation at the C9 position that was observed for *Yp*DHPS (DHP^+^) (Yun et al., 2012), and the occupancy of this compound was refined to 0.65. There was a clear break in electron density between the C9 position of the pterin ring and the *p*-amino group of *p*ABA (**Supporting Information Figure S4f**). These observations are consistent with *in crystallo*-capture of the Michaelis transition state of a S_N_1 enzymatic reaction mechanism of Sul3, wherein the PP_i_ group has been cleaved from the DHPP substrate but the *p*ABA molecule had not yet been ligated to DHP^+^. The position of DHP+ in the Sul3·6-MP·Mg^2+^·PP_i_ complex overlapped with its position in the *Yp*DHPS transition state complex structure (Yun et al., 2012). All the protein-ligand interactions observed in the Sul2·7,8-DHP·Mg^2+^·PP_i_ complex were observed in the Sul3·6-MP·Mg^2+^·PP_i_ complex, including the hydrophobic interaction between *p*ABA and Phe177.

### The pABA-interaction region is remodeled in Sul enzymes including insertion of a key Phe residue

Next, to gain insights into how Sul and DHPS enzymes compare in ligand interactions, we compared the Sul2/Sul3 complex structures with representative ligand-bound structures of *Ec*DHPS and DHPS from *Yersinia pestis* and *Bacillus anthracis* (Dennis et al., 2018; Yun et al., 2012). Superposition of these DHPS complex structures with the Sul2·7,8-DHP·Mg^2+^·PP_i_ complex or Sul3·6-MP·Mg^2+^·PP_i_ complex showed a high degree of overall structural conservation (RMSD 1.0-1.2 Å over 158-190 Cɑ atoms) except for conformational differences 3; loops 1 and 2 adopted a similar conformation in all the structures, dependent on full visualization of the DHPS active site (**Figure 3a**). The pterin group and the *p*ABA group of the ligands bound to Sul2/Sul3 or DHPS all occupied the same spatial locations. All the Sul2/Sul3 residues that interacted with the pterin ring are highly conserved in these representative DHPS enzymes (**Supporting Information Figure S5a, S6**). Similarly, while these DHPS structures did not contain Mg^2+^·PP_i_, the Sul2/Sul3 residues that co-ordinated these ligands are also conserved in these DHPS enzymes (**Supporting Information Figure S5a**).

In contrast, we noticed dramatic reorganization of the ɑ6 helix that formed the *p*ABA interacting region of the Sul enzymes (**Figure 3b**). The Sul enzymes contained a single Phe residue insertion (Phe178/Phe179/Phe177 in Sul1/Sul2/Sul3 respectively) relative to DHPS (**Figure 3b, 3c, 3d**). This Phe residue provided an additional hydrogen bond to *p*ABA through its backbone amide nitrogen (**Figure 3b**). Notably, the sidechain of this residue was localized near to the carboxylic acid group of *p*ABA and occupied a spatial location that overlaps with the sulfonamide drug group (i.e. the azole group of SMX), as indicated by the structure of the *Yp*DHPS·DHPP·SMX complex (**Figure 3c)**. The Sul enzymes also contained changes C-terminal to this Phe residue, including a deletion of a Gly residue from DHPS (Gly191 in *Yp*DHPS), and a substitution of a Leu residue (Leu180/Leu181/Leu179 in Sul1/Sul2/Sul3) in place of a Lys residue in DHPS (Lys92 in *Yp*DHPS) (**Figure 3b, 3c, 3d)**. This Leu residue was packed against Met176/Met177/Met175 (Sul1/Sul2/Sul3) which represents a substitution to smaller hydrophobic residue relative to the equivalent position in DHPS (i.e. Phe188). These changes in the Sul enzymes move the backbone torsion angles of residues between 179 to 181 into right-handed ɑ-helical conformation; in contrast, only *Yp*DHPS Gly189 in this region this region adopts a left-handed ɑ-helical conformation. C-terminal to ɑ6, the Sul enzymes contained two small amino acids (Ser-Pro in Sul1, Gly-Ala in Sul2, Gly-Ser in Sul3) that enabled a tight turn to connect to ɑ7 to continue the DHPS fold.

Altogether, since sulfonamides are *p*ABA structural analogs, these observations suggest that the sequence alterations in this region of the Sul enzymes play a key role in steric hindrance with sulfonamides and resistance thereto.

We also observed that the conformation of loop 3 in the Sul2·7,8-DHP·Mg^2+^·PP_i_ complex or Sul3·6-MP·Mg^2+^·PP_i_ complexes were nearly identical and their position fixed towards the active site (Fig. 3a). This loop formed numerous interactions with the key Phe (Supporting Information Figure S5b). In particular, Sul2 Arg138 and Sul3 Lys136 formed hydrophobic packing interactions (via the aliphatic region of their sidechains) with Sul2 Phe179 or Sul3 Phe177, respectively. As well, the backbone carbonyl oxygen of these Phe residues formed interactions with the backbone carbonyl oxygens of Sul2 Asp137 or Sul3 Thr135, as well as with the sidechains of Sul2 Gln132 or Sul3 Gln129. Sul2 and Sul3 contain an alanine residue (Ala136 or Ala134) that approach the key Phe residue. Notably, loop 3 in general is not well conserved among the Sul enzymes or DHPS enzymes (**Supporting Information Figure S6** and (Baca et al., 2000)), however, these arginine, glutamine and alanine residues mentioned above are highly conserved across the Sul enzymes. Collectively, these observations suggest loop 3 may form a complementary structure to stabilize the conformation of the key Phe residue in ɑ6.

Leveraging our structure of the Sul2·7,8-DHP·Mg^2+^·PP_i_ complex, we modelled all tested sulfonamides in the Sul2 active site through superposition with the *p*ABA region of DHP (**Supporting Information Figure S7**). This analysis showed that all compounds would pose a steric clash with the ɑ6 Phe residue, including the most primitive sulfa sulfanilamide, where the sulfonamide nitrogen approaches within 2.1 Å of this residue.

As implied by this structural analysis, the ɑ6 Phe residue is a key determinant of sulfa-resistance. To test this hypothesis, we mutated this Phe to Gly in Sul1 and Sul2 (Sul1^F178G^ and Sul3^F177G^) and tested the kinetic properties of these recombinantly-purified variants with *p*ABA and SMX. The Sul1^F178G^ and Sul3^F177G^ variants both showed similar K_M_, k_cat_ and catalytic efficiency parameters for *p*ABA as the WT Suls (**Table 1**), indicating that the dihydropteroate synthase reaction is unaltered by the Phe to Gly mutation. Interestingly, SMX inhibition of both the Sul1^F178G^ and Sul3^F177G^ variants was significantly reduced as compared to WT Sul1 and Sul3, with a drop in K_i_ parameters of 243- and 90-fold, respectively. As well, both the Sul1^F178G^ and Sul3^F177G^ variants showed increase preference for SMX as a substrate for the dihydropteroate synthase reaction, as demonstrated by increased K_M_ (107- and 68-fold, respectively) and increased k_cat_/K_M_ (206- and 135-fold, respectively) as compared to the WT Sul1 and Sul3. In essence, mutation of the ɑ6 Phe in Sul1 and Sul3 reverted these Sul enzymes to sulfa-susceptible DHPS enzymes, as all of their kinetic parameters more closely resembled those of *Ec*DHPS than the WT Sul1 and Sul3 enzymes. These results unambiguously show that the ɑ6 Phe plays the key role in sulfa-resistance in the Sul1 and Sul3 (and presumably, Sul2) enzymes.

### Sul enzymes differ from EcDHPS in active site dynamics in response to ligand-binding

Given that the Sul enzymes have a modified *p*ABA-binding site made up of the α6 helix and nearby loop 3, we hypothesized that Sul enzymes and *Ec*DHPS differ in conformational dynamics in response to ligand binding. Taking advantage of the fact that Sul enzymes naturally do not possess tryptophan residues, we performed a systematic mutagenesis approach to insert a single Trp residue in the α6 helix or loop 3 in the Sul enzymes and then evaluated ligand-induced changes in intrinsic tryptophan fluorescence (ITF) (**Figure 4**). Tryptophan fluorescence intensity has been a widely applied methodology to determine binding dissociation constants and protein conformational changes. Tryptophan residues are sensitive reporters of local environment and conformation changes in proteins. We interpreted changes in ITF (ΔF 340 nm) as indicating a change in the local environment of the Trp probe, consistent with a conformational change of the α6 helix and loop 3. In particular, the key α6 Phe residue (Phe178/Phe177 in Sul1/Sul3 respectively) was mutated to Trp (**Figure 4A, 4D**). Similarly, the loop 3 residues that closely interacted with the key α6 Phe (Sul1 Arg136 and Sul3 Lys136) were mutated to Trp. As our structure of the Sul3·6-MP·Mg^2+^·PP_i_ complex showed an organized loop 3 allowing a full view of the active site, we conducted a deeper mutagenesis study of this loop in Sul3 and singly mutated each of Ala133, Thr135, Val137 to Trp.

**Figure 4.**
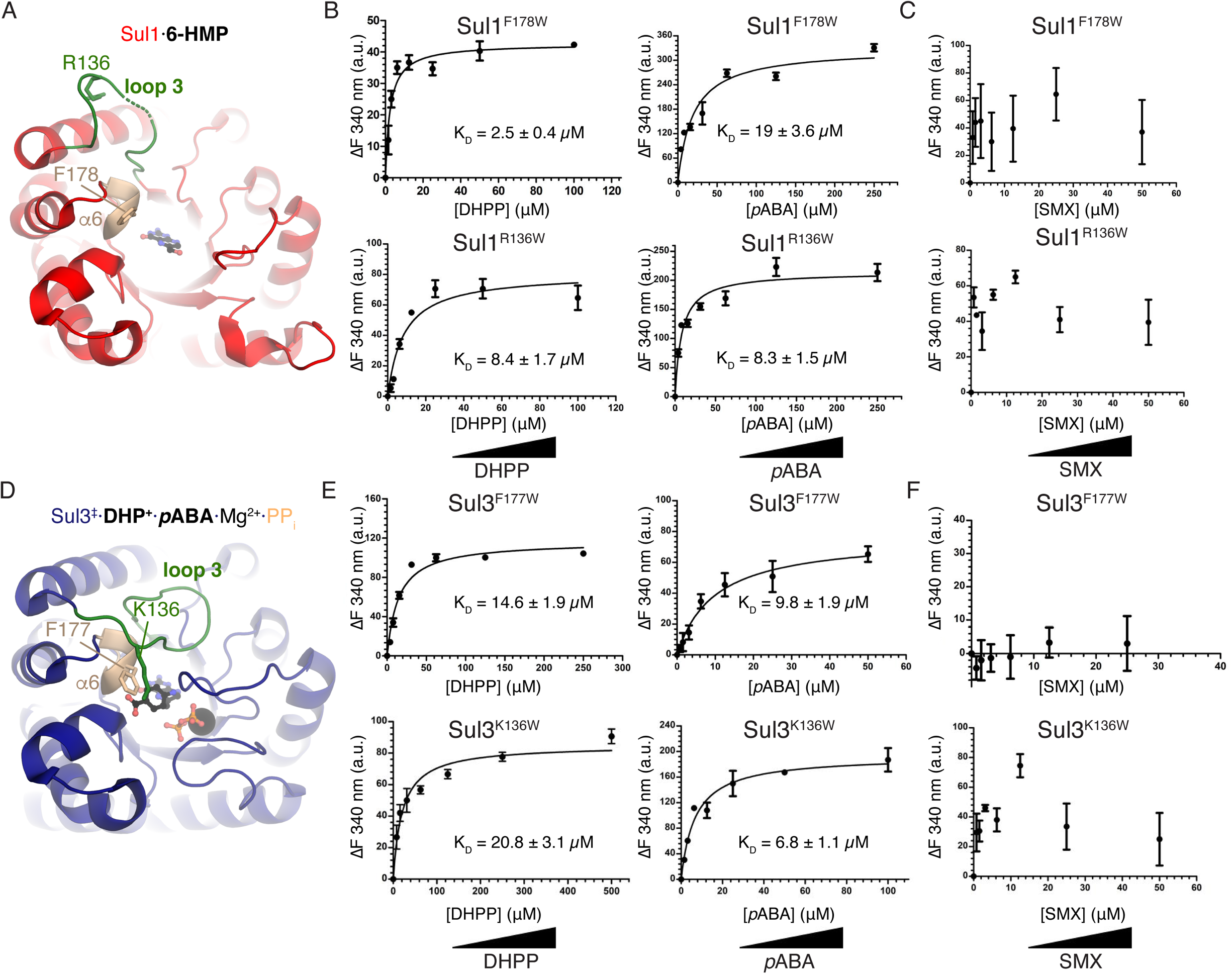
Sul enzymes’ ɑ6 helix and loop 3 respond to natural ligands but not to sulfas. A) Location of sites for introduction of Trp probes in the structure of Sul1. B) Titration of DHPP and *p*ABA, and C) SMX into Sul1^F178W^ (top) and Sul1^R136W^ (bottom) showing substrate-induced fluorescence emission intensity change at 340 nm (error bars represent ±SD) and calculated K_D_ values. Triplicate readings were average and K_D_ reported as (±SD). D) Location of sites for introduction of Trp probes in the structure of Sul3. E) Titration of DHPP and *p*ABA, and F) SMX into Sul1^F177W^ (top) and Sul1^K136W^ (bottom).

We first established that the WT Sul1 and Sul3 enzymes produce negligible background ITF signal (**Supporting Information Figure S8A, B**) and do not impact growth or sulfa resistance of a *de novo E. coli* Δ*folP* strain *trans*-complemented with pGDP::*sul1* W mutants (see next section for further discussion on this strain) (**Supporting Information Figure S9, Table S3**). All the Sul1 and Sul3 variants with Trp residues introduced into the α6 helix and loop 3 showed dose-dependent DHPP or *p*ABA ligand-induced fluorescence ITF changes (**Figure 5B, 5E** show the Sul1^F178W^, Sul1^R136W^, Sul3^F177W^ and Sul3^K136W^ variants; all other Sul3 variants are shown in **Supporting Information Figure S8C**). Note, according to the S_N_1 reaction of Suls, to investigate the ITF change in response to *p*ABA, we performed the ligand titration in presence of saturating excess of the first ligand, DHPP. We fit the ITF titration data to the Chergoff-Hill equation to derive the binding dissociation constants (K_d_) for DHPP and *p*ABA; the K_d_ derived for all the Sul1 and Sul3 variants studied by ITF were in a similar micromolar range (2-23 µM) (**Figure 5 and Supporting Information Figure S8**). The observation that all variants show similar K_d_ values for the ligands suggested that interactions with either were not largely altered by the mutations. These results indicated that as implied by our structural analyses, indeed, the α6 helix and loop 3 of Sul enzymes change conformation in response to the natural ligands.

**Figure 5.**
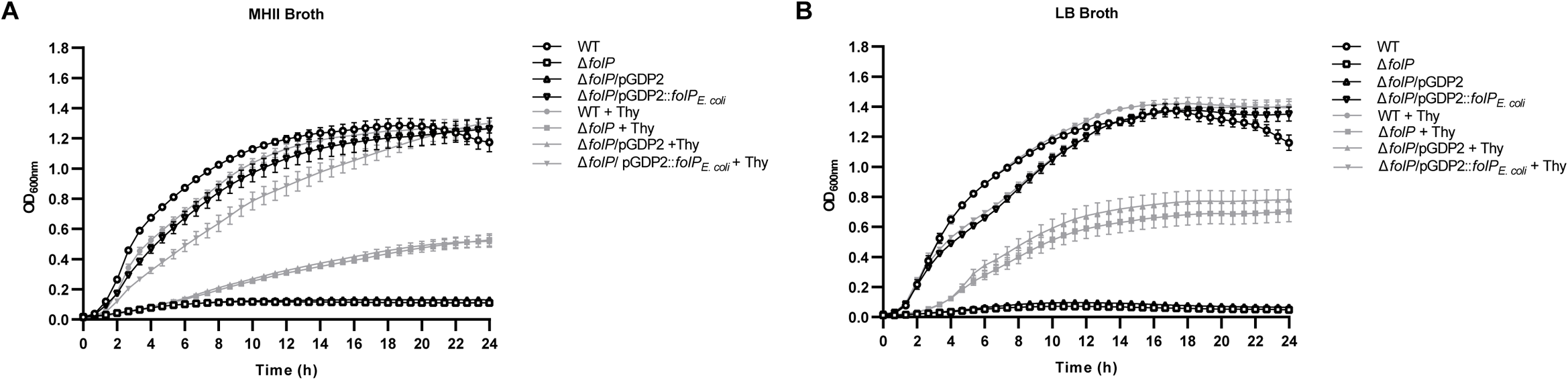
Addition of thymidine promotes the growth of the *E. coli folP* deletion strain. A) Growth curves for WT *E. coli* (WT) and *E. coli* Δ*folP* derivatives in MHII medium (A) or LB medium (B), without (black lines and symbols) or with (gray lines and symbols) 200 μg/mL thymidine (Thy). Data is taken from at least three biological replicates performed in technical triplicate. ODs at 600 nm were baseline-corrected by subtracting background absorbance at 600 nm of the uninoculated media from the absorbance at 600 nm of the inoculated media.

Next, using the ITF assay, we evaluated whether the α6 helix and loop 3 Trp mutants of the Sul enzymes showed conformational changes in response to sulfa drugs. We did not observe any ITF change in any of the Sul1 and Sul3 Trp variants with titration with SMX (**Figure 5C, 5F, Supporting Information S8**), indicating that the α6 helix and loop 3 residues do not change conformation and consistent with our data showing that the affinity of the Sul-sulfa interaction is hundreds-fold lower.

We also sought to understand if the regions of *Ec*DHPS equivalent to α6 helix and loop 3 undergo similar conformational changes in response to the natural ligands or SMX. The closest corresponding residues in the α6 helix (Phe190) and loop 3 of *Ec*DHPS were mutated to Trp (**Supporting Information Figure S10A**). Since *Ec*DHPS naturally possesses a tryptophan residue (Trp92) located on the opposite face of the (α/β)_8_ barrel (**Supporting Information Figure S10A**), to ensure the only ITF signal was from the Phe190Trp or Met148Trp probes, these variants also contain the mutation Trp92Phe. Indeed, the *Ec*DHPS^W92F^ mutation abrogates ligand-induced ITF signals (**Supporting Information Figure S10B**). Both the α6 helix and loop 3 variants (*Ec*DHPS^F190WW92F^ and *Ec*DHPS^M148WW92F^ variants, respectively) displayed DHPP-concentration dependent ITF changes, with similar calculated K_d_ values (8, 13 µM) (**Supporting Information Figure S10C**). However, titration experiments with *p*ABA in the presence of saturating excess DHPP produced no *p*ABA-induced fluorescence intensity changes. Finally, titration of the two *Ec*DHPS Trp variants with SMX also produced no ITF change (**Supporting Information Figure S10C**). We validated that none of these *Ec*DHPS mutations affects growth or sulfa resistance of our *E. coli* Δ*folP* strain *trans*-complemented with the corresponding mutants on pGDP::*folPEc* (**Supporting Information Figure S9, Table S3**). Together, these data indicated that no additional local conformational change occurs in these positions of *Ec*DHPS beyond that induced by DHPP binding.

### Deletion of the chromosomal folP gene in E. coli compromises growth in thymidine-limited media

To be able to assess the full contribution of the cloned *sul* genes and their mutated derivatives on conferring sulfonamide resistance and fitness of *E. coli*, an unmarked, in-frame *folP* deletion was constructed in the parent strain, *E. coli* BW25113 (see Experimental section for details). To date, two *E. coli folP* transposon mutant knockout strains have been made available to the research community, including the BW25113Δ*folP* strain and the widely-used C600Δ*folP* strain (Fermér and Swedberg, 1997); however, in both instances, the *folP* gene deletions are marked with a kanamycin resistance cassette. Furthermore, the BW25113Δ*folP* knockout strain (*E. coli* ID JW3144; Horizon Discovery), has previously been shown (Yamamoto et al., 2009) and reconfirmed here (data not shown) to harbor a partial duplication in which both the disrupted *folP* gene and the WT *folP* gene are present. Thus, to the best of our knowledge, this is the first report of an unmarked, in-frame *folP* deletion in *E. coli*.

In agreement with the previously reported thymidine-auxotrophic phenotype of the C600Δ*folP*::Kan^R^ mutant, deletion of the *folP* gene in BW25113 resulted in cells auxotrophic for thymidine – this mutant failed to grow in thymidine-limited Mueller-Hinton II medium, but grew, albeit slowly, in thymidine-supplemented MHII medium (**Figure 4 and Supporting Information Figure S11**) (Fermér and Swedberg, 1997; Lau et al., 2017). Similarly, the *folP* deletion strain also failed to grow in commercially prepared LB medium, which reportedly contains low amounts of thymidine (Giroux et al., 2017) but grew slowly in thymidine-supplemented LB medium (**Figure 4**). Still, the growth of the *folP* deletion strain in MHII or LB medium was only partially restored by the addition of thymidine (**Figure 4 and Supporting Information S11)**. The phenotypic morphology of the *folP* deletion strain on thymidine-supplemented solid media is shown in **Supporting Information Figure S11**. The WT strain formed large, circular, convex colonies, whereas the *folP* deletion strain formed small, circular colonies. Introduction of the wild type, *folP* gene on a low-copy-number plasmid, pGDP2 (Cox et al., 2017) into the Δ*folP* strain fully restored growth in thymidine-limited medium comparable to WT levels, whereas the empty vector control strain, like the *folP* deletion strain, failed to grow on thymidine-limited media and formed small colonies on thymidine-supplemented media (**Figure 4 and Supporting Information Figure S11**). Taken together, these data are consistent with (Fermér and Swedberg, 1997) in which an *E. coli folP* knockout requires the folate-end product, thymidine, for growth.

### Sul enzymes confer pan-sulfa resistance

Introduction of the individually cloned *sul1*, *sul2*, or *sul3* genes on pGDP2 into the *E. coli* Δ*folP* strain fully restored growth under thymidine-limited conditions comparable to the WT and *folP*/pGDP2::*folP_Ec_* strains (**Supporting Information Figure S12**), suggesting that the Sul enzymes can produce sufficient levels of 7,8-DHP to support growth (expression of the Sul enzymes was confirmed by Western blot, **Supporting Information Figure S13**). To assess the contribution of the individual Sul enzymes to sulfa resistance, the susceptibility of the *folP* deletion strain expressing *sul1*, *sul2*, or *sul3* to a range of sulfa antibiotics and co-trimoxazole was determined. As seen in **Tables 2 and S2**, expression of the Sul enzymes in the *folP* deletion strain resulted in high levels of *pan*-sulfonamide resistance and increased resistance to co-trimoxazole when compared with the WT *E. coli* and Δ*folP*/pGDP2::*folP_Ec_* strains. It is interesting to note that the Sul1-, Sul2-, or Sul3-expressing cells conferred the same level of resistance for each individual sulfa drug tested and for co-trimoxazole, although MIC values varied between sulfa drugs; the lowest degree of resistance was to SMX and sulfisoxazole (SOZ), and the highest degree of resistance was to sulfanilamide (SAA) and sulfadimethoxine (SDT).

**Table 2.**
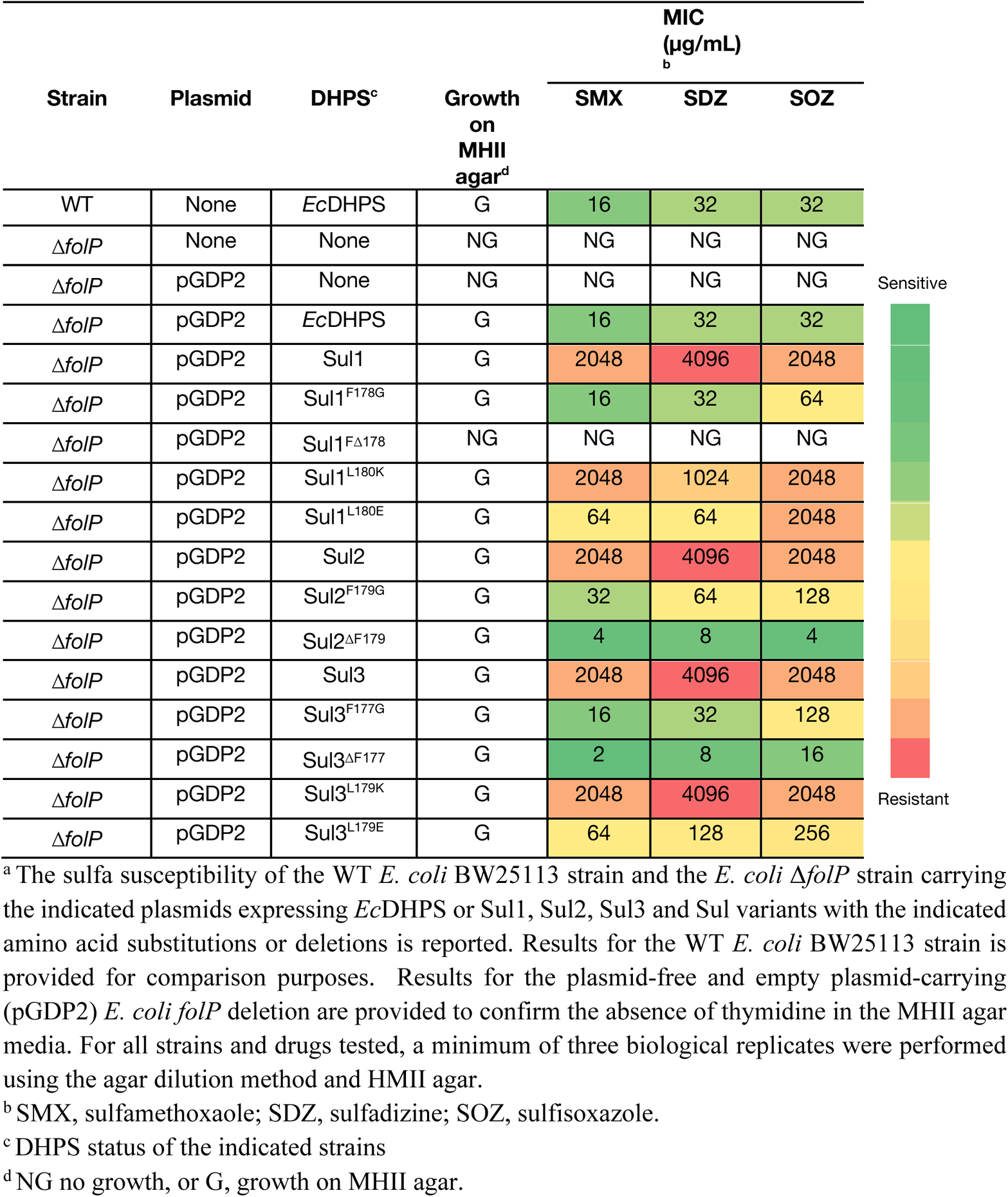
Sulfa susceptibility of the *E. coli* Δ*folP* deletion expressing WT or mutated Sul enzymes in the *p*ABA interaction region.^a^

### Sul enzymes intrinsic sulfa resistance is dependent on the α6 helix phenylalanine residue

To determine if the key Phe residue within the α6 helix of Sul enzymes plays a role in their intrinsic sulfa resistance, this residue was either mutated to a Gly residue or deleted in each of the *sul* expression constructs, and the impact of these mutations on sulfa resistance and complementation of the thymidine-auxotrophy of the *E. coli* Δ*folP* strain was assessed. Substitution of the Phe residue to Gly in Sul1, Sul2, and Sul3, compromised resistance to all sulfa drugs tested and to co-trimoxazole (**Table 2 and Table S2**) but did not impact bacterial growth compared to the Δ*folP* strains individually expressing the WT *sul* genes (**Supporting Information Figure S12**), or enzyme expression levels (**Supporting Information Figure S13**). The observed increase in susceptibility of the Sul Phe to Gly variants is consistent with our *in vitro* observations that drug binding to these variants in enhanced relatively to WT and that their folate production is not hindered by this mutation. Thus, this key Phe residue in the Sul1, Sul2, and Sul3 enzymes is essential for conferring *pan*-sulfonamide resistance.

Deletion of the key Phe residue in Sul2 and Sul3 further increased sulfa sensitivity compared to the equivalent Gly substitution mutants (**Tables 2 and S2)** but did not impact growth compared to the Δ*folP* strains individually expressing the WT Sul enzymes (**Supporting Information Figure S12**) or enzyme expression levels (**Supporting Information Figure S13**) in the Δ*folP* strain. In contrast, deletion of Phe178 in Sul1 resulted in *E. coli* cells unable to grow on MHII agar and, thus, sulfa MICs could not be determined, although this mutant was able to grow, albeit slowly, in MHII broth (**Supporting Information Figure S13**). Of note, deletion of Phe178 in Sul1 did not adversely impact protein expression levels of this enzyme when compared to the *folP*-deletion strain expressing the WT Sul1 enzyme (**Supporting Information Figure S13**). Therefore, deletion of the key Phe residue in Sul1 appears to diminish enzyme activity, folate production and ultimately, impaired bacterial growth, indicating its importance for maintaining the dihydropteroate synthase reaction.

We also evaluated whether the Leu residue on the α6 helix of Sul enzymes (Sul1 L180, Sul2 L181 and Sul3 L179) contributes to sulfa resistance. Mutation of this Leu residue to a negatively charged residue (Glu) in Sul1 or Sul3 dramatically compromised resistance to all sulfa drugs tested and to co-trimoxazole, whereas mutation to a positively charged side chain (Lys) modestly compromised resistance to several sulfa drugs and did not impact co-trimoxazole resistance level (**Tables 2 and S2)**. Mutation of the Leu residue to either a Glu or a Lys in Sul1 and Sul3 did not adversely impact bacterial growth (**Supporting Information Figure S12**) or Sul enzyme expression levels (**Supporting Information Figure S13**), suggesting that the observed decrease in susceptibility is due to enhanced enzyme-drug interactions. These data suggest that the hydrophobic sidechain of Leu is important for resistance and based on its location in the Sul enzyme structures, contributes to stabilizing the conformation of the α6 helix and key Phe residue.

### Adaptive laboratory evolution of FolP recovers a Sul-like Phe-Gly insertion that confers sulfa resistance

Having established that modifications of the *p*ABA-interaction region in the Sul enzymes is responsible for rendering these enzymes invulnerable to inhibition by sulfas, we were interested in whether such modifications in the chromosomally-expressed *Ec*DHPS enzyme would similarly confer resistance. To that end, we carried out four independent laboratory evolution experiments to assess if exposure of *E. coli* BW25113 to the prototypical sulfa drug, SAA, could select for amino acid mutations in *Ec*DHPS that mimic the Sul enzymes. To determine this, *E. coli* BW25113 was grown in media containing ½ the MIC of SAA (1024 μg/mL) and was passaged daily over a 7-day period into fresh media and drug. On Days 1 and 7, cells were serially diluted and plated on MHII agar without or with 256 μg/mL SMX. In all four trials, SAA exposure for one day did not result in the recovery of SMX-resistant (SMX^R^) colonies, indicating that longer-term exposure was necessary. Similarly on Day 1, no SMX^R^ colonies were recovered for the drug-free control and the vehicle control groups. Long-term (7-day) exposure of *E. coli* to SAA led to the recovery of SMX^R^ colonies and, again, no SMX^R^ colonies were recovered for the drug-free control and the vehicle control groups. 20 randomly selected SMX^R^ isolates (5 colonies from each trial) were subsequently picked and assessed for resistance to SAA and three additional sulfas (SMX, SDZ, SOZ). As seen in **Figure 6A**, 18 isolates showed an increase in resistance to all 4 sulfa drugs while two isolates showed an increase in resistance to SMX, SDZ, SOZ, but not SAA.

**Figure 6.**
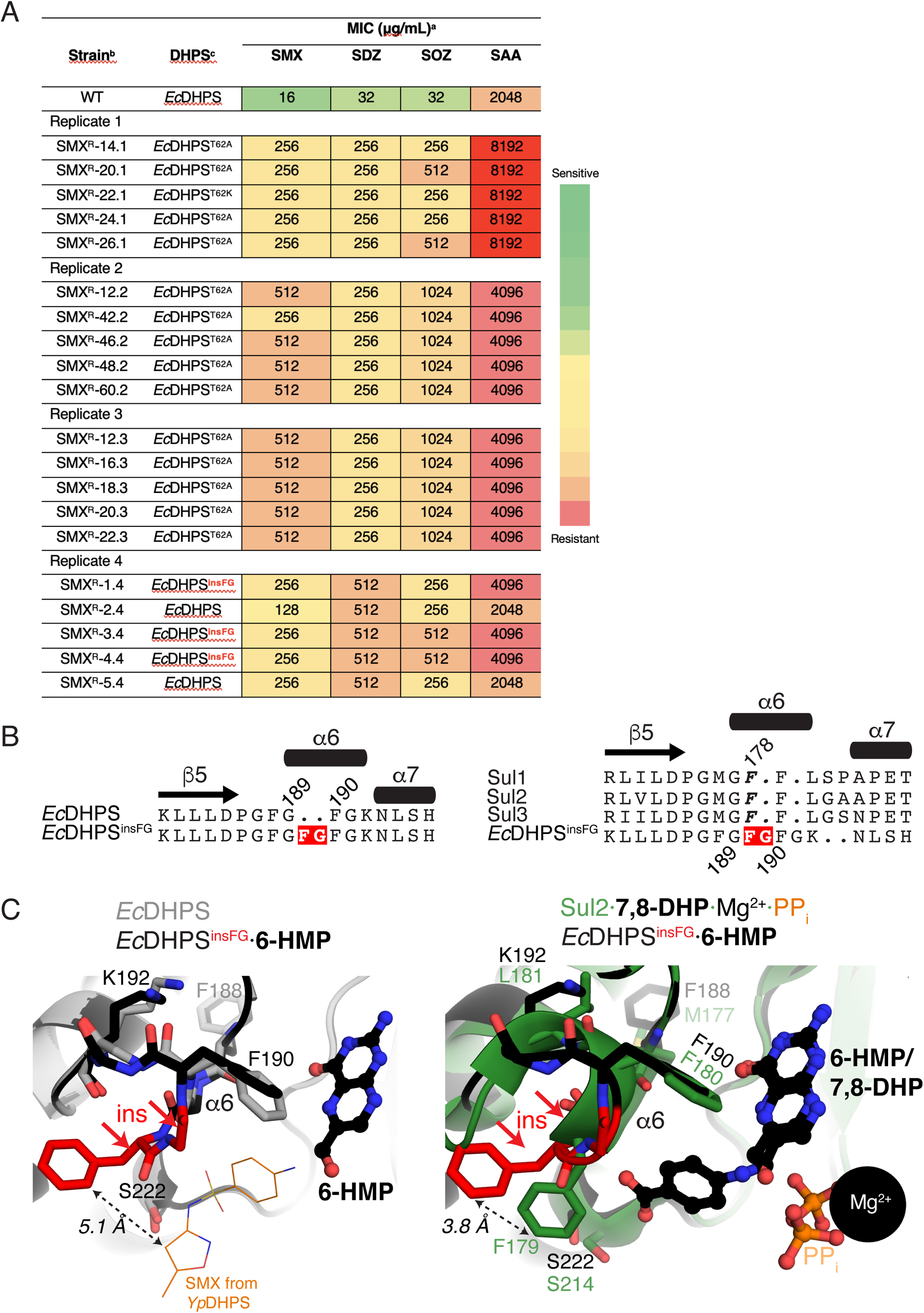
Adaptive laboratory evolution recovers a *Ec*DHPS mutant with Sul-like modifications in ɑ6. A) Sulfa resistance of the sulfamethoxazole-resistant mutants of *E. coli* BW25113 selected following a 7-day sulfanilamide exposure^a^. B) Sequence alignment of *Ec*DHPS^insFG^ insertion with *Ec*DHPS (left) and Sul enzymes (right), with insertion in *Ec*DHPS^insFG^ coloured red. Phe178 from Sul1 is indicated. C) Left = Comparison of the structure of the *Ec*DHPS^insFG^·6-HMP complex with *Ec*DHPS (PDB 1ajz). Also shown is the position of SMX from the *Yp*FolP·DHPP·SMX complex structure (PDB 3tzf) in thin lines. The distance between the inserted Phe and SMX is shown with a double-sided arrow. Right = Comparison of the structure of the *Ec*DHPS^insFG^·6-HMP complex with the Sul2·7,8-DHP·Mg^2+^·PP_i_ complex. The distance between the inserted Phe residue in *Ec*DHPS^insFG^ and Phe179 in Sul2 is shown with a double-sided arrow. Insertion FG sequence is coloured in red and with two red arrows in both panels. a Wild-type E. coli BW25113 was exposed to sulfanilamide (half the MIC; 1024 μg/mL) over 7 days and mutants resistant to 256 μg/mL of sulfamethoxazole (SMX) were selected. Four independent in vitro laboratory evolution experiments were conducted (Replicates 1-4). b Results are for 5 randomly selected SMX-resistant (SMXR) mutants from each independent trial are shown. Results for the *E. coli* BW25113 strain (WT) is provided for comparison purposes. For all strains and drugs tested, a minimum of 3 biological replicates were performed using the agar dilution method and MHII agar. c DHPS status of the indicated strains; The folP genes from each mutant was colony PCR amplified and sequenced to identify mutations in the DHPS coding region. EcDHPS, is the E. coli WT DHPS enzyme.

To assess the involvement of *Ec*DHPS in this sulfa resistance, the 20 recovered SMX^R^ mutants were examined for mutations in the *folP* gene. Among all the mutations found in the coding region of *Ec*DHPS, the most frequently observed mutations involved threonine 62 (15/20 isolates) (**Figure 6A**). Two SMX^R^ isolates were recovered in which no mutations were detected in the *folP* open reading frame, indicating the presence of a mutation(s) elsewhere in the chromosome that confers sulfa resistance in these isolates. Three SMX^R^ isolates contained an insertion of six base pairs coding for Phe-Gly, yielding *Ec*DHPS Gly187_Phe188insPheGly (*Ec*DHPS^insFG^). Intriguingly, sequence alignment showed that the location of the Phe-Gly insertion of *Ec*DHPS^insFG^ occurs in ɑ6, the same region that the Sul1, Sul2, and Sul3 enzymes possess a Phe insertion relative to DHPS (**Figure 6B**) and thus, we studied this mutant in greater detail.

To validate that the Phe-Gly insertion in *Ec*DHPS contributes to sulfa resistance in *E. coli*, we expressed the mutated *folP* gene (with a C-terminal FLAG-tag to monitor expression) on pGDP2 and introduced into the Δ*folP* strain to yield Δ*folP*/pGDP2::*folP*_insFG_. The ability to complement the thymidine-auxotrophy of the Δ*folP* strain and DHPS expression was also assessed. Introduction of the cloned *folP*_insFG_ gene into the Δ*folP* strain conferred increased resistance to all sulfa drugs tested and co-trimoxazole compared to that of the Δ*folP* strain expressing WT *Ec*DHPS on pGDP2 (**Table S4**), complemented the thymidine-auxotrophy of the *folP* deletion strain (**Supporting Information** Figure S14) and yielded WT levels of DHPS expression (**Supporting Information Figure S13**). These data indicate that this mutation contributes to *pan-*sulfa resistance. However, expression of *Ec*DHPS^insFG^ in the Δ*folP* strain did not confer the same level of sulfa resistance as the parent strains, suggesting that the presence of this resistant DHPS enzyme was not in itself sufficient to explain the high levels of sulfa resistance in these isolates. This suggests that an additional mutation(s) is present elsewhere in the chromosome that contribute to sulfa resistance in these isolates.

Next, we wanted to assess if introduction of a single Phe residue, rather than a Phe-Gly two residue insertion, at a similar position as the key Phe residues of Sul1, Sul2, and Sul3, is sufficient to impart sulfa resistance to the *Ec*DHPS enzyme. To test this, a Phe residue was inserted after amino acid 189 (Gly189_Phe190insPhe) within the WT *folP* sequence on plasmid pGDP2 to generate *folP*_Fins189_ and the impact of this mutation on sulfa susceptibility, ability to complement the thymidine-auxotrophy and DHPS expression assessed in the *folP* deletion strain. Surprisingly, insertion of Phe alone led to an increase in susceptibility to cotrimoxazole and most sulfa antibiotics except for sulfapyridine, for which the MIC remained unchanged compared to *E.coli* expressing WT DHPS (**Table S4**). Introduction of the cloned *folP*_Fins189_ restored growth to WT levels under thymidine-limited conditions and yielded WT levels of DHPS expression (**Supporting Figure S13**), suggesting that this mutation does not adversely impact DHPS enzymatic activity and folate production. Thus, insertion of Phe alone within the *Ec*DHPS enzyme is not sufficient to confer sulfa resistance, indicating an important role for the second inserted Gly.

To reveal the molecular basis of sulfonamide-resistance of *Ec*DHPS^insFG^, we crystallized the enzyme and determined its structure at 2.73 Å. The crystal grew in the presence of DHPP and *p*ABA, however, we observed electron density only for 6-HMP in the active site (**Fig. 6C, Supporting Information S4**). The structure revealed that the Phe-Gly insertion was localized to the ɑ6 helix, extending it from two to four residues, and the Phe residue was positioned near to the position of the key Phe residues in the Sul enzymes (Phe178/Phe179/Phe177 in Sul1/Sul2/Sul3, respectively) (**Fig. 6C**). Comparison of the *Ec*DHPS^insFG^·6-HMP complex with the Sul2·7,8-DHP·Mg^2+^·PP_i_ complex showed that this inserted Phe was positioned 3.8 Å from the position of Phe179 in Sul2. Comparison of the *Ec*DHPS^insFG^·6-HMP complex with the *Yp*DHPS·DHPP·SMX complex showed that the inserted Phe was 5.1 Å from the closest atom of the methoxazole group of SMX; the backbone carbonyl of the inserted Phe was positioned closer yet to this atom of SMX (4.2 Å). These distances could be closed by a rotation in the sidechain rotamer of the inserted Phe or from movement of the backbone of the inserted Phe and Gly residues. The structure of the *Ec*DHPS^insFG^·6-HMP complex also suggested that the role of the inserted Gly residue is to add flexibility to maintain the relative positioning of the inserted Phe and Phe192, a key residue for pterin ring binding (**Fig. 6C**). These observations indicate that insertions in the ɑ6 helix of DHPS enzymes that evolve through exposure to sulfas mimic the natural Sul enzyme sequence in this region and would, similarly to the Sul enzymes, occlude sulfa binding through steric repulsion.

## Discussion

In this era of the antibiotic resistance crisis and witness few new scaffolds being advanced to clinical use, an understanding molecular details of the mechanisms of resistance are essential for the modification of existing classes of compounds with established antibacterial activity. For example, insights into the structural and molecular mechanisms of β-lactamase enzymes have been leveraged in the design of new β-lactam-β-lactamase inhibitor cocktails progressing through clinical development (Bahr et al., 2021; Boyd et al., 2020). In stark contrast, little development of the sulfonamides, an established class of broad-spectrum antimicrobial, has occurred for many years. Reflecting a lack of attention to further diversification of the scaffold, the set of sulfonamides approved for clinical use all share a similar strategy of modification to the core 4-aminobenzenesulfonamide: acylation at the 2-N position with an aromatic moiety (e.g. azole, oxazole, pyridine). While this could be due to several factors, such as the attractiveness of alternative scaffolds, an important reason for the stalling of sulfonamide development is the spread of plasmid-borne sulfa resistance and the lack of molecular understanding into its mechanism. In fact, we underscore that all the approved sulfonamides are compromised by *sul-*based sulfa resistance. In this study we sought to uncover the molecular mechanism of *sul-*based sulfa resistance to provide a foundation for revisiting this antibacterial scaffold and generate avenues for next-generation DHPS/Sul enzyme inhibitors.

The *sul1* gene, in particular, is widespread in pathogenic and environmental species, yet its molecular basis of resistance has remained elusive for nearly 50 years. Its extreme dissemination likely reflects the historical focus on sulfas as antibacterial agents, their indiscriminate use in the clinic and environment in the early 20^th^ century, the resistance advantage or some other growth conferred by acquisition of the *sul1* gene. Using our *in vitro* activity assay, we clearly establish that each of the Sul1, Sul2 and Sul3 enzymes harbors kinetic parameters comparable to *Ec*DHPS as our representative of DHPS enzymes. As well, each *sul* gene fully complemented the growth defect observed upon knockout of *folP*. While we did not monitor production of compounds in the folate biosynthetic pathway downstream of the DHPS product 7,8-dihydropteroate, these data indicate that these enzymes can replace a sulfa-inactivated DHPS enzyme, at least in *E. coli*, without any obvious fitness defects.

To the best of our knowledge, our work is the first demonstration of the *in vitro* kinetic parameters of Sul enzymes and establish an assay for evaluation of potential inhibitory compounds. Using this system, we evaluated the characteristics of the Sul enzymes for utilization of a representative sulfa drug as a substrate or as an inhibitor. Our data clearly demonstrate that SMX does bind the Sul enzymes and it is utilized as a substrate, condensing with the pterin moiety of DHPP to form a covalent adduct. This dead-end product cannot be used in the next step of the folate biosynthesis pathway, which mirrors the mechanism of action of sulfas versus DHPS enzymes. However, our assay showed that the catalytic efficiency for formation of this adduct is more than 150-fold less than for the archetypal *Ec*DHPS. In terms of inhibition, SMX is also a much worse inhibitor of the Sul enzymes than *Ec*DHPS (more than 80-fold worse). While the kinetic properties of Sul1, Sul2 and Sul3 for utilization of the native substrate *p*ABA were very similar, we observed a decreasing series of K_M_ and K_I_ parameters for SMX for Sul1, then Sul2, followed by Sul3. This could reflect the sequence divergence nearby the *p*ABA binding site. Our observations establish that the sulfonamide scaffold does bind Sul enzymes, but very weakly.

Through exhaustive structural analysis of each of the Sul1, Sul2 and Sul3 enzymes, we provide molecular details underlying their intrinsic sulfa resistance. Our observations established that the basis for their intrinsic sulfa resistance is based on modifications of *p*ABA-interaction region of their active sites, which places a Phe residue in position to intercept sulfa drug binding through steric clash. Mutation of the Phe residue to glycine restored *in vitro* binding of SMX to each of the Sul enzymes while not compromising 7,8-dihydropteroate synthesis, indicating that this residue plays a role exclusively in sulfa resistance but not catalysis. Accordingly, *E. coli* Δ*folP trans*-complemented with any of the *sul* genes mutated to encode this Phe-to-Gly substitution abrogated the *pan*-sulfa resistance conferred by these enzymes. The importance of the Phe-Gly insert was underscored in a “gain-of-function” lab-evolved mutant *folP* that resembles the Sul enzymes in possessing this insertion in the equivalent region of the *Ec*DHPS active in the same region as this natural sequence in the Sul enzymes. We also demonstrate that both amino acid insertions are necessary, as a Phe-only insert in a variant *folP* did not confer sulfa resistance. This observation is consistent with our crystal structures showing that the Gly residue plays a role in proper positioning of the inserted Phe residue to occlude sulfa binding.

As further corroboration of the relevance of modifications in the ɑ8 region of the *p*ABA-interaction site of dihydropteroate synthases for resistance, sulfa-resistant clinical isolates of *Neisseria meningitidis* strains MO035 and 418 were shown to harbor a six bp insertion, coding for a Ser-Gly insertion in the *N. meningitidis* DHPS enzyme (*Nm*DHPS) (Fermer et al., 1995). Deletion of this insertion in the *Nm dhps* gene compromised *N. meningitidis* resistance to sulfa drugs (Fermer et al., 1995). Sequence alignment revealed that this Ser-Gly insertion in the *Nm*DHPS enzyme localizes to the same region of ɑ6 as the Phe-Gly insertion identified in the *Ec*DHPS^insFG^ enzyme (**Figure 6b**). Modeling of *Nm*DHPS with the Ser-Gly insertion using AlphaFold2 (Jumper et al., 2021) indicates that this region would resemble the Phe-Gly insertion in *Ec*DHPS^insFG^; the inserted Ser residue would occlude sulfa binding by approaching it by less than 3 Å (**Supporting Information Fig. S14**). A Phe-Gly insertion is also naturally present in three *Ec*DHPS enzymes (NCBI Genbank IDs: HAX1960291.1, EFG0716961.1 and EFI6540711.1). In all three instances, however, the contribution of the *Ec*DHPS^insFG^ mutation to sulfa resistance in *E. coli* was not examined, but we predict these would also be sulfa-resistant and arose due to sulfa exposure.

A Gly-Phe-Phe sequence in ɑ6 of DHPS naturally occurs in hundreds of sequences in NCBI Genbank and has previously been identified as a hallmark of Sul and Sul-like dihydropteroate synthases; some of these were shown to confer sulfa resistance when expressed in *E. coli* (Sánchez-Osuna et al., 2019). Our data are consistent with this study, but our structural data provides the functional rationale behind their role in resistance and refines the motif to insertion of a Phe before a Phe residue that is conserved in Sul and DHPS enzymes (**Figure 3D, Supporting Information Figure S6**), and substitution of a Lys residue in DHPS enzymes to a Leu residue in Sul enzymes. Assuming that Sul-like dihydropteroate synthases with this Phe-Gly insertion are sulfa-resistant, the significance of the natural existence of this sequence in such enzymes is not clear and remains a subject for further research. For example, are bacteria harboring such Sul-like dihydropteroate synthases exposed to sulfa or sulfa-like compounds in their environment and thus require intrinsic resistance, or is it fortuitous? Do such Sul-like dihydropteroate synthases play a role in other biosynthetic pathways through condensation of DHPP with compounds other than *p*ABA, where a more confined binding site would be advantageous? As a corollary, why do some *folP* genes, like those of *Ec*DHPS, *Yp*DHPS and *Ba*DHPS as we analyzed in this study, *not* harbor this Gly-Phe-Phe sequence although provides a selective advantage for growth in the presence of sulfa or sulfa-like compounds? Is there a fitness cost to harboring the Gly-Phe-Phe sequence under conditions not studied here? Furthermore, the ultimate origin and the mechanism by which the sul genes or their predecessors mobilized from the chromosome to plasmids remains elusive.

By tracking conformational changes in the active site of Sul and *Ec*DHPS enzymes via tryptophan fluorescence in the presence of ligands, our analysis also revealed that Sul enzymes possess increased conformational flexibility within the ɑ6 region (harboring the Phe-Gly insertion) and the nearby loop 3 that is responsive to both *p*ABA and DHPP. In contrast, these regions of *Ec*DHPS change conformation only in response to DHPP binding. This is consistent with previous studies that showed the rate-limiting step of DHPS activity is the release of pyrophosphate after DHPP binding, which then forms the *p*ABA binding site (S_N_1 ordered reaction); the lack of additional conformational change in response to *p*ABA by *Ec*DHPS suggests the *p*ABA binding site is rigidified at this stage of the catalytic cycle. With Suls, we observed similar kinetic behavior, where binding of DHPP was first required followed by *p*ABA binding, but even after the first step and conformational changes in response to DHPP, ɑ6 and loop 3 retain additional flexibility. We interpret this observation as an additional capability of Sul enzymes to adopt the conformation necessary to occlude sulfa binding. This has implication for drug design targeting Sul enzymes, which may require interactions that stabilize the conformation of ɑ6 and loop 3 for favorable binding. Notably, loop 3 has not been previously implicated in the catalytic cycle of DHPS enzymes and is not conserved in these enzymes but does show conservation of particular residues (a Lys/Arg, a Gln, and an Ala residue) in Sul enzymes, consistent with a functional role.

The behaviour of the Sul enzymes in the crystal structures, activity assay, ITF were consistent with a sequential ordered binding model reaction mechanism, similar to DHPS enzymes. For crystallography, to resolve the Sul2·7,8-DHP·Mg^2+^·PP_i_ and Sul3·6-MP·Mg^2+^·PP_i_ complexes required a first incubation of the enzymes with DHPP, followed by *p*ABA. In the activity assay, this is reflected in our observation that we could not reach saturation of DHPP in the presence of excess *p*ABA, while the inverse was true. Similarly in the ITF assay, *p*ABA produced ligand-induced fluorescence changes in Sul1 and Sul3 only in presence of excess DHPP. These observations are consistent with DHPP ligand is a necessary first step in the reaction, releasing the pyrophosphate group, followed by the *p*ABA ligand leading to product formation of 7,8-DHP. We interpret the electron density of the Sul3·6-MP·Mg^2+^·PP_i_ complex as showing the carbocation formed after release of pyrophosphate from DHPP, and this structure was extremely similar to that observed for *Yp*DHPS (Yun et al., 2012).

Although many studies have demonstrated that expression of the *sul* genes alone or in combination in *E. coli* confers sulfa resistance, few have employed quantitative assays for MIC determination and of those that have, only a limited number of sulfa drugs have been tested including sulfamethizole (Blahna et al., 2006; Enne et al., 2001; Handford et al., 2009; Kerrn et al., 2002; Perreten and Boerlin, 2003; Rådström et al., 1991; Wu et al., 2010; Zhou et al., 2021). This study thus represents a comprehensive study quantifying the contribution of plasmid-borne *sul* genes to several sulfa antibiotics in a single strain of *E. coli* devoid of additional resistance genes. The results demonstrate that expression of the individual *sul* genes in *E. coli* results in high levels of resistance for all 12 sulfonamide antibiotics tested and in all instances, the MICs surpassed the CLSI resistant MIC breakpoints, indicating that the *sul* genes alone are sufficient to compromise sulfa therapy (“M100Ed32 | Performance Standards for Antimicrobial Susceptibility Testing, 32nd Edition,” n.d.) Moreover, sulfa bacteriostatic potency varied between sulfas for each strain tested and this is consistent with previous studies in which the bacteriostatic activity of sulfonamides correlates with the pKa_2_ values of the sulfa drug (i.e. the lower the pKa_2_ value, the lower the MIC) (Tappe et al., 2008; Zarfl et al., 2008). Since sulfas are used in combination with trimethoprim, which targets a different step in folate biosynthesis, we also demonstrated that expression of the individual *sul* genes in a *folP*-deficient *E. coli* strain conferred increased co-trimoxazole (sulfamethoxazole-trimethoprim) resistance; however, the co-trimoxazole MIC values observed were lower than the CLSI susceptible breakpoint MIC for this drug. These data suggest that the *sul* genes alone are not sufficient to compromise co-trimoxazole therapy and, not surprisingly, that the presence of additional resistance elements (i.e. acquired resistant variants of *dhfr*, the target of trimethoprim) are required to confer clinically relevant co-trimoxazole resistance in *E. coli*.

Analysis of the genetic changes in the *E. coli folP* gene following long-term exposure to sub-inhibitory concentrations of SAA led to the identification of two types of sulfa-resistant isolates. One type harboured mutations in the *folP* gene including codon 62 for threonine or an insertion coding for a Phe-Gly insertion. To the best of our knowledge, both of these *folP* variations and their association with sulfa-resistance have not been reported for *Ec*DHPS. While we did not assess the contribution of the *Ec*DHPS Thr62 substitution to sulfa-resistance in this study, an equivalent substitution in the *Mycobacterium leprae* DHPS enzyme, Thr53, and in the fungal pathogen *Pneumocystis carinii* DHPS enzyme, Thr55, have previously been documented and associated with sulfa resistance (Cambau et al., 2006; Howard, 1975; Kai et al., 1999; Maeda et al., 2001; Mei et al., 1998; Williams et al., 2001). This is not particularly surprising, as Thr62 of *E. coli* is highly conserved in bacterial DHPS enzymes (Achari et al., 1997; Hampele et al., 1997), is located in loop 2 of the *Ec*DHPS enzyme and has been shown to directly interact with the β-phosphate of DHPP via its main-chain NH residue and its hydroxyl group (Achari et al., 1997), or the aniline NH_2_ of the product pteroic acid (Achari et al., 1997; Dennis et al., 2018). The *Ec*DHPS Thr62 mutation was the most common mutation present in the evolved isolates, suggesting that this mutation is preferentially selected for by sulfanilamide; however, future experiments aimed at deep sequencing of the *folP* gene of the evolved bacterial populations may provide further insight into the type and prevalence of sulfa-selected *folP* mutations. Still, it is unclear if *Ec*DHPS Thr62 contributes wholly or partially to the observed sulfa-resistance in these isolates and thus, further studies characterizing the contribution of the Thr62 mutation to sulfa resistance in *E. coli* are warranted. Importantly, *folP* mutations have been shown in this study and in others to explain only partially acquired bacterial resistance to sulfas (Buwembo et al., 2013; Vedantam et al., 1998). For example, *trans*-complementation of a mutated *folP* gene, coding for a Pro64Ser substitution, from sulfathiazole-resistant lab strains of *E. coli* to a wild-type *E. coli* strain resulted in only a low level of sulfathiazole resistance compared to the original isolates (Vedantam et al., 1998). This ultimately led to the identification of additional chromosomal mutations in the sulfathiazole-resistant *E. coli* isolates that were shown to contribute to sulfa resistance (Vedantam et al., 1998). Similarly, in *Streptococcus mutans*, *folP* mutations only partially explained sulfa-resistance, although the identity of these additional chromosomal mutations were not examined (Buwembo et al., 2013). Indeed, we demonstrated that *trans*-complementation of the *Ec*DHPS^ins188FG^ mutant in the *folP* deletion strain only partially restored resistance, consistent with previous observations that additional chromosomal mutations contribute to high-levels of sulfa resistance. Surprisingly, the second type of sulfa-resistant isolates recovered did not harbour any *folP* mutations and demonstrated an increase in resistance to sulfamethoxazole, sulfadiazine, and sulfisoxazole, but not sulfanilamide. These data suggest that non-*folP* sulfa-resistant mutation(s) can be polyspecific (i.e. confer resistance to some sulfa drugs but not all) and may involve other mutations. At present, it is unclear as to why sulfanilimide selected for resistance to other sulfonamides, but not itself. Further experiments including whole genome sequencing of both types of sulfa resistance mutants, *folP* mutants and non-*folP* resistant mutants isolates will undoubtedly enhance our understanding of the development of sulfa resistance in *E. coli* and other bacterial pathogens.

Our structural data allowed modeling of sulfa drugs in the active site of the Sul enzymes. This analysis showed that the ɑ6 Phe residue would clash with the sulfonamide nitrogen and the aromatic substituents at this position of all sulfa drugs approved for use in clinical practice. Thus, we posit that a core pharmacophore is necessary to inhibit Sul enzymes. This compound would need to have restricted size so as to enter the *p*ABA binding pocket of Sul enzymes, evade the ɑ6 Phe residue and/or leverage it for favorable hydrophobic interactions. There has been extensive research into the inhibition of DHPS enzymes through non-sulfonamide scaffolds (Azzam et al., 2020; Babaoglu et al., 2004; Dennis et al., 2018; Hammoudeh et al., 2013; Hevener et al., 2010; Pemble et al., 2010; Zhao et al., 2016, 2012) however to the best of our knowledge, no campaigns have been conducted to discover Sul inhibitors, or, compounds capable of inhibiting both Sul and DHPS enzymes. Given the widespread dissemination the *sul1* gene in particular, dual Sul1+DHPS inhibitors would be advantageous. The molecular and structural data we obtained the Sul enzymes can be utilized to initiate such efforts and provide new avenues for overcoming sulfa resistance.

## Experimental

### Antibiotics

Sulfonamide antibiotic stock solutions for susceptibility testing were prepared as follows: sulfamethoxazole (SMX) (Sigma-Aldrich), 50 mg/mL in DMSO; sulfadiazine (SDZ) (Sigma-Aldrich), 100 mg/ml in 1 M NaOH; sulfanilamide (SAA) (Sigma-Aldrich), 50 mg/ml in acetone; sulfisoxazole (SOZ) (TCI America) 50 mg/ml on acetone; sulfathiazole (STZ)(Sigma-Aldrich) 50 mg/ml in acetone: sulfapyridine (SPY) (Sigma-Aldrich) 40 mg/ml in 0.5M NaOH; sulfamerazine (SMRZ) (AK Scientific), 50 mg/ml in DMSO; sulfamethazine (SMZ) (Alfa Aesar), 50mg/ml in DMSO; sulfameter (SMT) (Sigma-Aldrich) 100 mg/ml in DMSO; sulfacetamide (SAD) (Sigma-Aldrich) 100 mg/ml in DMSO; sulfaquinoxaline (SQX) (Sigma-Aldrich), 50 mg/ml in DMSO; sulfadimethoxine (SDT) (Sigma-Aldrich), 100 mg/ml 1N NaOH.

### Bacterial strains and growth conditions

The bacterial strains and plasmids used in this study are described in **Table S5**. Primers used in the study are listed in Table S6. Bacterial cells were cultured in Luria broth (L-broth) and Luria agar (L-agar), Mueller Hinton II broth (MHII-broth) and Mueller Hinton II –agar with antibiotics, 200 *μ*g/mL thymidine, and 40 *μ*M folinic acid as necessary, at 37 °C. Plasmid pKOV was a gift from George Church (Addgene plasmid # 25769; http://n2t.net/addgene:25769; RRID:Addgene_25769) (Link et al., 1997). In *E. coli* plasmid pKOV and its derivatives were maintained or selected with 25 ug/mL of chloramphenicol at 30^○^C, plasmid pGDP2 and its derivatives were maintained or selected with 25 *μ*g/mL kanamycin. Plasmids pMCSG53 and its derivatives were maintained or selected with ampicillin 100*μ*g/mL Plasmids pNIC-CH and its derivatives were maintained or selected with kanamycin at 50 *μ*g/mL.

### DNA Methods

Standard protocols were used for restriction endonuclease digestion, ligation, transformation, and agarose gel electrophoresis, as described by Sambrook and Russell (Sambrook J, Russell DW (2001) Molecular cloning: a laboratory manual, 3rd ed. Cold Spring Harbor, N.Y.: Cold Spring Harbor Laboratory Press). Plasmid and chromosomal DNA was prepared as before (Fruci and Poole, 2018). DNA fragments used for cloning were extracted from agarose gels using a Wizard^®^ SV gel and PCR clean-up system (Fisher Scientific, Ltd., Nepean, Canada). Dephosphorylation and ligation into the gene replacement vector pKOV was carried out using the Rapid DNA Dephos & Ligation kit (Roche) according to the protocol provided by the manufacturer. CaCl_2_-competent *E. coli* cells were prepared as described (Inoue et al., 1990). Electrocompetent *E. coli* cells were prepared using a method described by New England Biolabs. Oligonucleotide synthesis was carried out by Integrated DNA Technologies (Coralville, IA). Nucleotide sequencing was carried out by ACGT Corp. (Toronto, Canada) using universal primers. In several instances, genes were designed *in silico*, synthesized and/or mutated and cloned into the required plasmid at BioBasic Inc (Markham, Canada). For *in vitro* protein characterization and crystallization, cDNA for *sul1, sul2, sul3* were purchased from IDT DNA Technologies as gblock gene fragments (1-269 amino acids for Sul1, residues 1-271 amino acids for Sul2 and 1-242 amino acids for Sul3) and were codon optimized for expression in *E.coli*. All primers were purchased from IDT DNA technologies. cDNA for *sul2, sul3* were cloned into the vector pMCSG53, coding for a fusion protein with an N-terminal 6x histidine tag cleavable by TEV protease. cDNA for *sul1* was cloned into the vector pNIC-CH, coding for a fusion protein with a non-cleavable 6x histidine tag. *E.coli* K12 DHPS (1-282 amino acids) and *E.coli* K12 HPPPK (*folK*) were cloned into the vector pMCSG53 for enzymatic kinetic studies. The deletion mutants and catalytically inactive mutants of all the Sul enzymes were prepared by using a method modified from the QuickChange site directed mutagenesis protocol (Stratagene, La Jolla, CA).

### Protein purification

*E.coli* BL21(DE3) Gold competent cells were transformed with the pMCSG53 or pNIC-CH expression plasmids harboring *sul1*/*sul2*/*sul3* or *folP*. 20 mL of overnight culture (approx. 16 h growth time) in LB broth was diluted into 1 L of LB media containing selected antibiotics (kanamycin for pNIC-CH plasmids, ampicillin for pMCSG53 plasmids) and grown at 37 °C with shaking. Expression was induced with 0.5 mM IPTG at 17 °C when OD600 reached 0.8 units and allowed to grow overnight for 16-18 h. The large overnight cell cultures were then collected by centrifugation at 7000g. Cells were resuspended in a binding buffer [pH 7.5, 100 mM HEPES, 500 mM NaCl, 5 mM imidazole, and 5% glycerol (v/v)] and lysed. The cell debris was removed by centrifugation at 20000 g. Ni-NTA affinity chromatography was used for protein purification. The soluble cytoplasmic fraction containing the target protein was purified by batch-binding to a Ni-NTA resin, washed with wash buffer [pH 7.5, 100 mM HEPES, 500 mM NaCl, 30 mM imidazole, and 5% glycerol (v/ v)], and the protein was eluted with elution buffer [pH 7.5, 100 mM HEPES, 500 mM NaCl, 250 mM imidazole, and 5% glycerol (v/v)]. The His6-tagged protein was then subjected to overnight (approx. 16 h) TEV protease cleavage using 50 μg of TEV protease per mg of His6-tagged protein and simultaneously dialyzed overnight against a buffer containing no imidazole. The His6-tag and TEV were removed by binding the protein onto a Ni-NTA column again, whereby the flowthrough contained the His-tag cleaved protein of interest. For plasmid constructs with non-cleavable His6-tag, the TEV protease cleavage step was skipped. The final proteins were dialyzed with a minimum of 2 x 2 L dilution cycles in 10 mM HEPES at pH 7.5 with 300 mM NaCl, 0.5 mM TCEP for crystallization or in 50 mM HEPES at pH 7.5 with 300 mM NaCl, 0.5 mM TCEP for kinetics. The purity of the protein was analyzed by SDS-polyacrylamide gel electrophoresis and mass spectrometry. The purified proteins were also subjected to size exclusion chromatography (Superdex 200 16/60) analysis for determination of their oligomeric state. The proteins were concentrated using a Vivaspin concentrator (GE Healthcare) and passed through a 0.2 µm Ultrafree-MC centrifugal filter (Millipore) before storing in aliquots at −80C.

### Crystallization, x-ray data collection and structure solution

All crystallization was performed at room temperature of 21 °C using vapor diffusion method in sitting drops with 0.6 μL protein and/or protein:ligand substrate mix plus 0.6 μL reservoir solution, using a TTP Labtech Mosquito protein crystallization robot. *p*ABA for co-crystallization was used as 20 mM stock solution in water. DHPP was used as 5 mM stock solution in 50 mM HEPES pH 7.5, 10 mM magnesium chloride (MgCl_2_), 3% (v/v) dimethyl sulfoxide (DMSO). For the Sul1·6-HMP complex crystal, 1 mM solution of protein was incubated for 2 h with 2.5 mM DHPP at 4 °C followed by passing through a 0.2 µm Ultrafree-MC centrifugal filter. The reservoir solution for this crystal was 25% (w/v) PEG 5KMME, 0.2 M ammonium sulfate, 0.1 M Tris (pH 8.5), 1% (w/v) tri-isobutyl methylphosphonium tosylate. The crystal was cryoprotected by transferring to well solution with 10% (v/v) 2-Methyl-2, 4-pentanediol followed by paratone oil. The Sul2 apo crystal was grown in reservoir solution 1.6 M ammonium sulphate, 2% (v/v) 1,6-hexanediol, 0.1 M HEPES (pH 7.5) and cryoprotected either by transferring in reservoir solution with 2% (v/v) PEG 200 followed by paratone or using only paratone oil respectively. The Sul2·7,8-DHP·Mg^2+^·PP_i_ complex crystal was grown by incubating a 1 mM solution of protein with 2 mM DHPP for 30 min at 4 °C followed by adding 2 mM *p*ABA and further incubating for 2 h at for 30 min at 4 °C and finally passing through a 0.2 µm Ultrafree-MC centrifugal filter. The reservoir solution for this crystal was 1.6 M ammonium sulfate, 2% (v/v) hexanediol, 0.1 M HEPES pH 7.5. The crystal was cryoprotected by transferring to well solution with paratone oil. Wild type Sul3 failed to form crystals. We utilized the surface entropy reduction approach (Cooper et al., 2007) and the double mutant E142A+E143A crystallized. For the Sul3 apoenzyme complex, Sul3.E142A+E143A crystals were grown in reservoir solution of 2 M ammonium sulphate, 5% (w/v) isopropanol and cryoprotected by transferring in reservoir solution with 2.5% (v/v) glycerol followed by paratone. For the Sul3·6-HMP complex, a 1 mM solution of Sul3 and 2 mM DHPP was incubated for 1h at 4 °C, then grown in reservoir solution 20% (w/v) PEG3350, 0.1 M calcium chloride and 0.05 magnesium chloride. The crystal was cryoprotected by transferring to reservoir solution containing 0.5% trehalose, 0.5% glycerol and paratone oil. For the Sul3·7,8-DHP·*p*ABA·Mg^2+^·PP_i_ complex, a 1 mM solution of Sul3.E142A+E143A was incubated for 30 min at 4 °C with 2.5 mM DHPP followed by 5 mM *p*ABA for a further 2 h at 4 °C and finally passing through a 0.2 µm Ultrafree-MC centrifugal filter. Crystals were grown in reservoir solution 2 M ammonium sulphate and 5% (w/v) isopropanol. The crystal was cryoprotected by transferring to a reservoir solution containing 12% (v/v) glycerol, 5% (v/v) trehalose followed by paratone. Prior to data collection, crystals were flash frozen in liquid nitrogen. X-ray diffraction data at 100 K was collected at the beamline 19-ID, Structural Biology Center, Advanced Photon Source, Argonne National Laboratory. All diffraction data were processed using HKL3000 (Minor et al., 2006). Molecular Replacement was utilized to solve each structure, using the structure of *Ec*DHPS (PDB 1AJ0) (Achari et al., 1997) as the search model, or the apoenzyme version of each enzyme, with Phenix.refine (Liebschner et al., 2019). All model building and refinement were performed using Phenix.autobuild, Phenix.refine and Coot (Emsley et al., 2010). The positions of all ligands was evaluated using simulated annealing omit maps using Phenix.refine. All geometry was evaluated using Phenix.molprobity and the wwPDB validation server. Atomic coordinates have been deposited in the Protein Data Bank with accession codes 7S2I, 7S2J, 7S2K, 7S2L, 7S2M, 7S2O, 7TQ1. Production of figures and analysis was performed using The PyMOL Molecular Graphics System, Version 2.4.0 Schrödinger, LLC. Sequence alignment was performed by the Clustal Omega server with manual adjustments based on structural superpositions, and figure was produced using the ESript server.

### In vitro dihydropteroate synthase activity assay

The substrate, DHPP, was synthesized based on previous methods (Hevener et al., 2010) and its purity was validated by mass spectrometry. *E.coli* K12 HPPK (6-hydromethyl-7,8-dihydropterin pyrophosphokinase) with N-terminal 6x His tag was purified using Ni-NTA chromatography as described in the above section. The substrate 6-HMP (purchased from Schirks Laboratories, Switzerland) was subjected to an enzymatic conversion at 37 C for 1h with shaking using 5 mM ATP, 30 mg/mL HPPK, 10 mM MgCl_2_, 3% DMSO in 50 mM HEPES pH 7.5. The reaction mixture was filtered through a 3 kDa MWCO filter wherein the HPPK was retained in the column. The filtrate containing the product DHPP and traces of ATP and any unconverted substrate 6-HMP was analyzed by mass spectrometry. 99% conversion was achieved as observed by the depletion of the substrate 6-HMP by mass spectrometry. The purified DHPP was droplet frozen in aliquots and stored at −80 C. The substrate, *p*ABA was purchased from Sigma. Dihydropteroate synthase activity by DHPS/Sul enzymes was monitored spectrophotometrically by a coupled Malachite green assay (Baykov et al., 1988). Malachite green solution was prepared in concentrated sulphuric acid (1.1 g in 150 mL conc. H_2_SO_4_ to total 1 L with water). Ammonium molybdate (Fluka) was prepared as a 7.5% (w/v) solution in water. For the assay, fresh Malachite green reaction (MGR) mix was prepared by mixing 4 mL Malachite green solution, 2 mL 7.5% (w/v) ammonium molybdate, 80 uL 11% Tween 20. The free pyrophosphate that was released in the DHPS reaction was cleaved to ortho-phosphate by using inorganic *E.coli* K12 pyrophosphatase in the enzymatic reaction. Inorganic *E.coli* K12 pyrophosphatase (0.1 U/αL) was purchased commercially from Cell Signalling Tech., NEB. The enzyme reaction was carried out in 96-well format containing the substrates DHPP and *p*ABA/SMX, *Ec*DHPS/Sul enzymes, 10 mM MgCl_2_, 2% DMSO, 50 mM HEPES pH 7.5, 0.01 U of inorganic *E.coli* pyrophosphatase (EC 3.6.1.1, NEB), all in a 100 µL reaction volume in the well. The 96-well plate reaction was incubated on a benchtop temperature-controlled plate shaker (ELMI) at 37 C for 20 min at 500 rpm. Ortho-phosphate was colorimetrically detected by adding 25% of total well reaction volume of fresh MGR mix, shaking for 2 min and monitoring absorbance. Absorbance values at 630 nm were instantly monitored on a Infinity MPlex Tecan plate reader with an linear shaking of 60s at 2.5 mm amplitude and 25 pixel readings per well read-out. Data from three independent biological replicates (triplicate data points) were averaged. Kinetic parameters (K_M_) were determined by nonlinear least-squares fitting of the data to the Michaelis-Menten equation, using GraphPad Prism v5.0 software (GraphPad Software, U.S.).

For the bisubstrate enzyme reaction, two different experimental set up was performed. In the first case for DHPP kinetics, DHPP concentrations were varied in the range of 0-500 µM in the presence of 10-fold excess of KD_, pABA_ at 200 µM. For the second experiment, for *p*ABA kinetics, *p*ABA concentrations in the range of 0-200 µM were used in presence of the DHPP concentration set at 20-fold excess of its K_d_ at 200 µM. For the SMX kinetics, SMX concentrations were in the range of 0-20 mM for Suls and 0-80 µM for Sul mutants and *Ec*DHPS, while the DHPP concentration was set at 20-fold excess of its K_d_ at 200 µM. For the Ki determination with SMX, three different SMX concentrations were tested for inhibition, with varying *p*ABA concentrations (0-200 µM) and excess DHPP at 200 µM. Ki for each SMX concentration was calculated using the formula K_i_ = [I] / [(K_Mobs_/K_M_)-1] and averaged. Here, [I] is the SMX concentration used, K_Mobs_ is the observed K_M_ at the respective inhibitor SMX concentration and K_M_ is at zero inhibitor SMX concentration.

### Intrinsic tryptophan fluorescence (ITF)

Site directed mutagenesis (as described earlier) was used to generate the following Trp mutations in Sul1, Sul3 and *Ec*DHPS were introduced into plasmids pMCSG53/pNIC-CH for protein expression and purification or pGDP2 for introduction into the *folP* deletion strain. The experiment was performed in black, opaque 96-well microtiter plate (ThermoScientific) in a 100 µL total reaction volume containing 1 µM of enzyme in 50 mM HEPES pH 7.5, 10 mM MgCl_2_, 2% DMSO. Serial dilutions of substrate DHPP (5 mM stock concentration), pABA (0.5 mM stock concentration), SMX (200 mM stock solution in DMSO), SDZ (200 mM stock solution in DMSO) were used for the experiment. To account for background signal from the ligand, control buffer titrations with the ligand alone (negative control) were performed in triplicate for each titration point. The plate was allowed to incubate for 30 min at 37 °C on a bench plate shaker (ELMI BioSciences). Tryptophan fluorescence spectroscopy experiments were recorded on an Infinite MPlex Tecan plate reader with the following settings: top-read, λex 295 nm, λem 340 nm, photomultiplier gain 40, 60s shaking prior to endpoint read and 25 pixel points/well read-out. Intrinsic tryptophan fluorescence spectra were recorded at 295 nm excitation wavelength and emission scan recorded. Both 285 nm and 295 nm excitation wavelengths produced intensity changes but 295 nm was chosen due to lower background signal and to eliminate any contribution to the fluorescence signal from tyrosine and phenylalanine residues in the protein. The background fluorescence quenching caused by protein dilution with the buffer was monitored by running parallel buffer control titrations. The averaged negative control data was subtracted from the ligand titration data for each titration point to remove the background signal from the ligand (F). The value of F before the start of titration (no ligand added), where the protein is unsaturated is F_o_. The value of (F) at each ligand concentration was subtracted from (F_0_) and this gives the change, ΔF for each ligand concentration. Readings for each experiment were recorded in triplicate and averaged. Data from three independent experiments (biological replicates) were analyzed by plotting fluorescence intensity changes ΔF against corresponding ligand concentrations and fitted to a one-site binding equation (Chergoff Hill’s plot) to generate K_d_ and B_max_. K_d_ is the equilibrium dissociation constant and Bmax is the maximum specific binding. Graphpad Prism v4.0 software (GraphPad Software, U.S.) was used for curve fitting.

### Construction of E. coli BW25113 ΔfolP mutant

To generate an in-frame *folP* gene deletion in WT *E. coli* BW25113, ∼ 1 kb fragments upstream and downstream of *folP* were first PCR amplified and cloned into plasmid pUC19, and then subsequently subcloned into the gene replacement vector pKOV. The *folP* upstream fragment was PCR amplified using: Δ*folP* Up For (HindIII) 5’-GACT **<u>AAGCTT</u>** CAGGTTGTGGTCGGCTTGCC-3’Δ*folP* Up Rev (XbaI) 5’-GACT **<u>TCTAGA</u>** GGCAAAGAGTTTCATGATGTTATCCCTGG-3’ and the *folP* downstream fragment was amplified using Δ*folP* Dn For (XbaI) 5’- GACT **<u>TCTAGA</u>** GTGGTGGAAGCCACTCTGTCTGCA-3’ and Δ*folP* DN Rev BamHI 5’-GACT **GGATCC** AGCCAGTTATCTAACGCTTT-3’. The 1kb upstream and downstream fragments were PCR amplified from the chromosome of *E. coli* BW25113 parent strain in two, spate 50 µl mixtures that contained 1 ug of BW25113 chromosomal DNA, 0.6 µM of the appropriate primer set, 0.2 mM of each dNTP, 1 x Phusion HF buffer, 5% (vol/vol) DMSO, and 1 unit (U) of Phusion DNA polymerase (Finnzymes, New England Biolabs, Pickering, ON, Canada). The mixture was heated for 30 sec at 98°C, followed by 30 cycles of 30 sec at 98°C, 30 sec at 65.0°C, 30 sec at 72°C, concluding with 7 min at 72°C. The PCR products were subsequently gel-purified and digested with HindIII and XbaI or XbaI and BamHI, as appropriate, and separately cloned into appropriately restricted pUC19, yielding plasmids pMJF200 (upstream fragment) and pMJF201 (downstream fragment). Both plasmids were individually introduced into DH5α, and transformants were selected on ampicillin 100 μg/mL. Plasmid DNA was recovered and sequenced to confirm the absence of mutations in the cloned fragments, the upstream fragment was excised from pMJF200 by digestion with HindIII-XbaI and was cloned into HindIII-XbaI restricted pMJF201, yielding pMJF202.

To generate the *folP* deletion construct in the gene replacement vector pKOV, the *folP* upstream and downstream fragments were PCR amplified from the purified plasmid, pMJF202 with the primers Δ*folP* For NotI 5’-GACT **GCGGCCGC** CAGGTTGTGGTCGGCTTGCC-3’ and DN Rev BamHI (see above) in a 50 µl mixture that contained 10 ng of template, 0.6 µM of the appropriate primer, 0.2 mM of each dNTP, 1 x Phusion HF buffer, 5% (vol/vol) DMSO, and 1 U of Phusion polymerase (New England Biolabs). The mixture was heated for 30 sec at 98°C, followed by 30 cycles of 30 sec at 98°C, 30 sec at 65.0°C, 1 min at 72°C, concluding with 7 min at 72°C. The resulting ∼ 2 kb PCR fragment was gel purified, digested with NotI and BamHI, and cloned into dephosphorylated NotI-BamHI restricted, pKOV, to yield pKOV::Δ*folP* (pMJF203). pMJF203 DNA was transformed into DH5α, and transformants selected on 25 ug/mL of chloramphenicol at 30^○^C. Plasmid DNA was recovered and then sequenced to ensure that no mutation(s) had been introduced during PCR. pMJF203 was then electroporated into *E. coli* Keio BW25113 and allowed to recover for 1 h at 30°C. Electroporants were then plated on prewarmed LB agar plates containing 25 μg/mL of chloramphenicol, 200 μg/mL of thymidine, and 40 μM of folinic acid and incubated overnight at 42 °C. From the 42 °C plate, 4-5 colonies were picked and suspended into 1 mL of LB broth with no NaCl, serially diluted, and immediately plated on 5% w/v sucrose, no NaCl, 200 μg/m of thymidine, and 40 μM folinic acid and then incubated at 30 °C overnight. The next day, colonies were replica plated onto LB agar containing 5% w/v sucrose, no NaCl, 200 μg/mL of thymidine, and 40 μM folinic acid, Mueller Hinton II agar ( no thymidine) to screen for loss of DHPS activity, and LB agar containing thymidine, folinic acid and chloramphenicol to confirm loss of the replacement vector. Of the 250 colonies that were replica plated, 2 colonies failed to grow on MHII agar, but not the sucrose containing LB agar plates. Deletion of the *folP* genes was confirmed using colony PCR with primers Δ*folP.* For NotI and DN Rev BamHI. A 10 µl colony PCR reaction mixture contained 2 µl of the chromosomal DNA solution as the template, 0.6 µM of each of primer Δ*folP* For NotI and DN Rev BamHI, 0.2 mM of each dNTP, 1 x Thermopol buffer, 5% (vol/vol) dimethyl sulfoxide (DMSO), and 1 U of Taq DNA Polymerase (New England Biolabs, Whitby, ON, Canada). The mixture was heated for 3 min at 95°C, followed by 35 cycles of 30 sec at 95°C, 45 sec at 61.6°C, 3:30 min at 72°C, concluding with 5 min at 72°C.

### Growth Curve Assay

To monitor the impact of the WT or mutated FolP/Sul enzymes on *E. coli* growth, an *in vitro* growth curve assay was performed. *E. coli* strains were cultured overnight in MHII broth supplemented, as required, with 40 μM folinic acid, 200 μg/mL thymidine and/or 25 μg/mL kanamycin. Overnight cultures were diluted 1/10 in fresh MHII broth, pelleted by centrifugation (13000 rpm, 1 min), washed twice with MHII broth and then standardized to an OD600 nm of 0.1. Using Corning Costar, Clear, 96-well round bottom microplates, bacterial cells were diluted in fresh MHII broth to a final OD600 nm of 0.05. Growth was monitored at an OD 600 nm, every twenty minutes for 24 hrs, with a MultiSkan GO Plate Reader and SKANIT Software CF. In all instances, media only controls were carried out to ensure the absence of contamination.

### Sulfonamide antibiotic susceptibility testing

The susceptibilities of various *E. coli* strains to sulfonamide antibiotics was assessed using the agar dilution method outlined previously (Wiegand et al., 2008) with the exception that MHII agar (Sigma) plates were used, as this agar is explicitly formulated to have low levels of thymine and thymidine content, both of which are known to antagonize the action of sulfonamides. Briefly, freshly streaked *E. coli* colonies were resuspended in sterile 0.9% NaCl solution and the turbidity of the suspension adjusted to that of a McFarland Standard of 0.5 using a Sensititre Nephelometer calibrated to a McFarland 0.5 BaSO_4_ standard. The bacterial suspension was then diluted 1:10 into a well of a sterile 96-well microtitre plate by pipetting 10 ul into a well containing 90 ul of sterile saline. A 48-pin replicator with 1.5 mm pins was used to deliver the final inoculum of 1 μl (∼10^4^ CFU/mL) onto MHII agar plates containing various concentrations of either sulfamethoxazole (SMX), sulfadiazine (SDZ), sulfanilamide (SAA), sulfisoxazole (SOZ) sulfathiazole (STZ), sulfapyridine (SPY), sulfamerazine (SMRZ), sulfamethazine(SMZ), sulfameter (SMT), sulfacetamide (SAD), sulfaquinoxaline (SQX), sulfadimethoxine (SDT). Of note, the MIC for SMT at 8192 μg/mL and for SQX at ≥2048 μg/mL could not be determined as the sulfa drug precipitated out of solution at these concentrations. Inoculated sulfa-agar plates were incubated 37°C for 18 hrs and the minimum inhibitory concentration (MIC) was determined to be the lowest concentration of drug that inhibited bacterial growth. In all instances, a drug-free control plate was included to ensure bacterial growth. Of note, the Δ*folP* deletion strain, which is auxotrophic for thymidine and thus, cannot grow on MHII agar was included as a negative control (no growth control).

MIC values for co-trimoxazole were determined by E-test using the Trimethoprim-sulfamethoxazole (1:19) MIC test strips (Liofilchem, Italy). The MICs were read according to the E-test reading guide for TMP-SMX where the inhibition ellipse intersects the strip at 80% inhibition.

### Whole-cell protein extracts and Western Immunoblotting

Overnight cultures of *E. coli* strains grown in MHII-broth supplemented, when necessary, with 40 μM folinic acid, 200 μg/mL thymidine, and kanamycin 25 μg/mL, were subcultured (1:49) in MHII broth and incubated at 37 °C and shaking at 200 rpm until bacterial densities reached an OD600 of 0.5-0.6. Cells were then standardized to an OD at 600 nm of 0.5 and pelleted by centrifugation at 13,000 rpm. Pellets were resuspended in 200 μl 1x phosphate buffered saline (PBS) and 200 μl 2 x RedMix. Samples were heated at 95 °C for 5 min and sonicated for 25 s at 30% amplitude. Whole-cell protein extracts were then separated on a 12% Mini-PROTEAN TGX Stain-Free protein gels and electroblotted onto a PVDF membrane using a BioRad Trans-BlotR Turbo™ Transfer System according to manufacturer’s instructions. Blotted membranes were subsequently incubated in PBS containing 0.1% (vol/vol) Tween 20 (PBST) and 10% (wt/vol) skim milk for 1 hr. Following two 5-min washes with PBST, the membranes were incubated for 60 min with a primary mouse monoclonal ANTI-FLAG M2 antibody (1:5000 dilution; Sigma-Aldrich), in PBST containing 1% (wt/vol) bovine serum albumin (Sigma-Aldrich) and then washed four times for 10 min each time with PBST. A secondary polyclonal Goat Anti-Mouse IgG H&L (HRP) antibody (1:5000 dilution; Abcam, Massachusetts) in PBST containing 1% bovine serum albumin was then added to the membranes and incubated for 1 hour, and subsequently washed four times for 10 min each time with PBST. All incubations and washings in the immunoblot procedure were carried out at room temperature with gentle agitation. Blots were developed by using the Clarity Western ECL Substrate (Biorad), according to the manufacturer’s instructions and the blots were visualized using DNR Bio-Imaging Systems MicroChemi 4.2 imaging system with GelCapture (DNR Bio-Imaging) software following 15 s exposure.

### Adaptative laboratory evolution

To assess the ability of the prototypical sulfonamide, sulfanilamide, to select for *folP* mutations, a 7-day mutant selection experiment was performed. One hundred microlitres of an overnight culture of *E. coli* Keio BW25113 in MHII broth was transferred into 10 mL of fresh MHII-broth containing either 1024 μg/mL (1/2 MIC) of sulfanilamide or no drug. Acetone, which was used to solubilize sulfanilamide, was included as a vehicle control (at similar concentrations as used in the sulfanilamide-exposed cells). Following a 24-hour incubation period at 37°C, shaking at 200 rpm, 100 μl of culture was transferred to fresh MHII broth containing the same concentration of sulfanilamide or vehicle control or no drug. This process was repeated every 24 hours over a 7-day period. On day 1 and 7, cultures were monitored for resistance development by serially diluting and spread plated in technical duplicate onto MHII-agar containing 256 μg/mL SMX or no drug. Following an 18-hour incubation, presumptive sulfanilamide-selected SMX-resistant colonies were enumerated and used to calculate CFU/mL, resistance frequency [(defined as CFU/mL total *E. coli* (no drug plate) by CFU/mL of SMX-resistant *E. coli*)] and fold-change (defined as resistance frequency of sulfa-treated group divided by resistance frequency of the vehicle control group) for each treatment. Biological triplicates were performed for all treatments. Presumptive SMX-resistant mutants were randomly selected and patched onto MHII agar plates with or without 256 μg/mL or 512 μg/mL of SMX to confirm SMX resistance. A total of 20 SMX-resistant mutants (5 from each biological replicate) were screened for chromosomal *folP* mutations by amplifying the *folP* gene using colony PCR followed by gel-purification and DNA sequencing. A 50 μl colony PCR reaction mixture contained 10 μl of the chromosomal DNA solution as the template, 0.5 μM of each primer (*folP* For: 5’-CGACGCACCGCAGATTGATGACCTG-3’ and *folP*Rev: 5’-CCAGTGCTGACTCCAGCATATAGCC-3’), 0.2 mM dNTPS, 1 x Phusion HF buffer, 5% DMSO, and 1 unit of Phusion DNA polymerase (New England Biolabs). The mixture was heated for 30 sec at 98°C, followed by 35 cycles of 30 sec at 95°C, 30 sec at 65°C, 30 seconds 72°C, and concluding with 7 min at 72°C. The *folP*-containing PCR product was gel-purified and sequenced to confirm the presence of *folP* mutations.

To assess if the identified *folP* chromosomal mutations contributed to sulfa resistance, confirmed *folP* mutants were colony PCR amplified using primers carrying a c-terminal FLAG sequence fusion that DHPS production levels could be monitored, and subcloned into the expression vector, pGDP2. Colony PCR was performed in a similar manner as described above except the following primers were used to amplify the mutated chromosomal *folP* gene: *folP* ins188FG For 5’-GACTTCTAGATTAACTTTAAGAAGGAGATATACATGAAACTCTTTGCCCAGGGTAC-3’; XbaI site underlined) and 200 μM FMR (5’ – GACTAAGCTT**TTA***CTTGTCGTCATCGTCTTTGTAGTC*CTCATAGC GTTTGTTTTCCTTT - 3’; HindIII site underlined, FLAG-tag DNA sequence italicized, and stop codon bolded). PCR amplicons were column purified, restriction-digested with HindIII and XbaI and ligated into HindIII-XbaI restricted, alkaline dephosphorylated pGDP2. Ligated plasmid DNA was transformed into *E. coli* DH5α and transformants were selected on kanamycin 25 μg/mL. Plasmid DNA was recovered and sequenced to confirm the presence of the desired mutation and the absence of PCR-generated mutations. The resultant plasmid, pMJF224 was then introduced into calcium chloride competent *E. coli* Δ*folP* via heat-shock and plasmid-bearing cells were selected on L-agar plates containing kanamycin 25 μg/mL.

## Acknowledgements

We thank Rosa Di Leo for assistance with cloning. We thank the staff of the Structural Biology Center, Advanced Photon Source, Argonne National Laboratory (Youngchang Kim, Kemin Tan, and especially Karolina Michalska) for x-ray diffraction data collection and/or data processing. We thank Robert Flick and the BioZone Mass Spectrometry Facility for analysis of synthesis of DHPP and evaluation of formation of SMX-pterin adduct. We thank Gerry Wright for providing plasmid pGDP2. This work has been funded in whole or in part with U.S. Federal funds from the National Institute of Allergy and Infectious Diseases, National Institutes of Health, Department of Health and Human Services, under Contract No. HHSN272201700060C (Center for Structural Genomics of Infectious Diseases (CSGID, http://csgid.org). This project was funded in part by an AAFC project grant to MF.

