## Supporting Information for "Molecular mechanism of plasmid-borne resistance to sulfonamides"

### SUPPORTING INFORMATION FIGURES

**Supporting Information Figure S1. Chemical structures of sulfonamide compounds.**

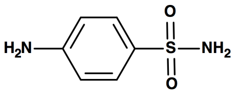

sulfanilamide
(SAA)

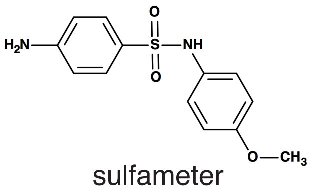

sulfameter
(SMT)

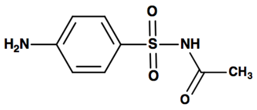

sulfacetamide
(SAD)

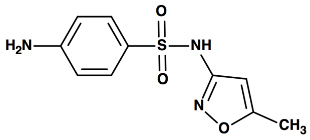

sulfamethoxazole
(SMX)

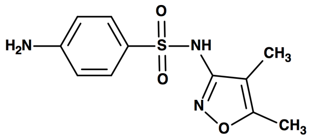

sulfisoxazole
(SOZ)

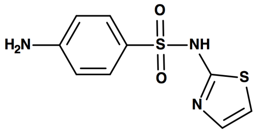

sulfathiazole
(STZ)

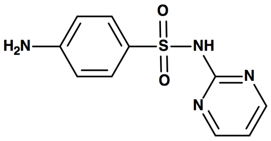

sulfadiazine
(SDZ)

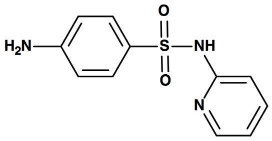

sulfapyridine
(SPY)

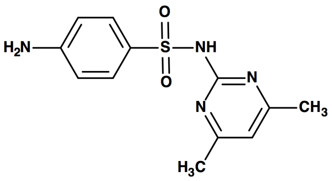

sulfamethazine
(SMZ)

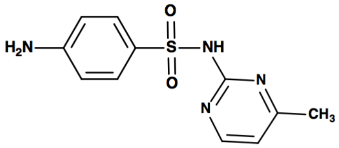

sulfamerazine
(SMRZ)

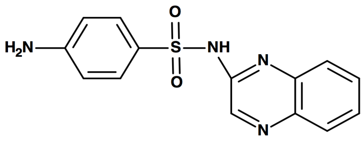

sulfaquinoxaline
(SQX)

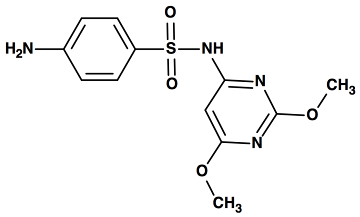

sulfadimethoxine
(SDT)

**Supporting Information Figure S2. Michaelis-Menten kinetics plots for *p*ABA for Sul and *Ec*DHPS enzymes.**

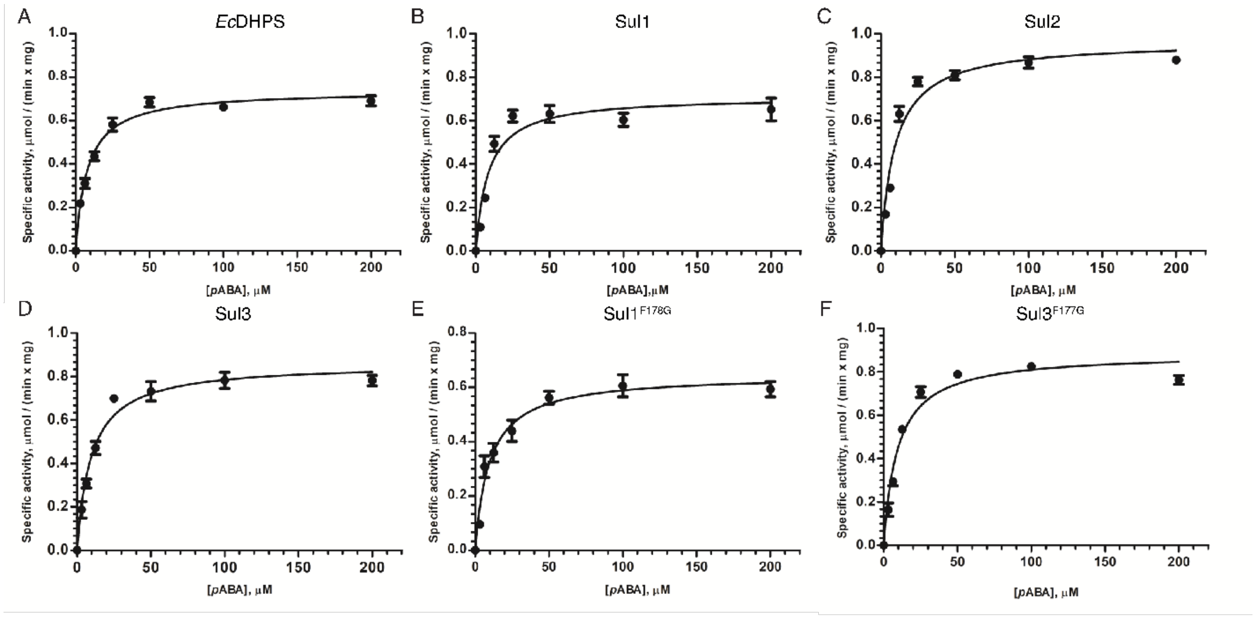


A) *Ec*DHPS, B) Sul1, C) Sul2, D) Sul3, E) Sul1$^{F178G}$, F) Sul3$^{F177G}$ for the substrate *p*ABA in the presence of saturating excess of DHPP at 200 mM. The kinetic parameters, $K_M$ and $V_{max}$ were estimated using the Michaelis-Menten equation by non-linear regression using GraphPad Prism v5.0. Each data point is the mean of three biological replicates for each substrate concentration. Each replicate is plotted as mean ± SD.

**Supporting Information Figure S3. Michaelis-Menten kinetics and inhibition plots for SMX for Sul and *Ec*DHPS enzymes.**

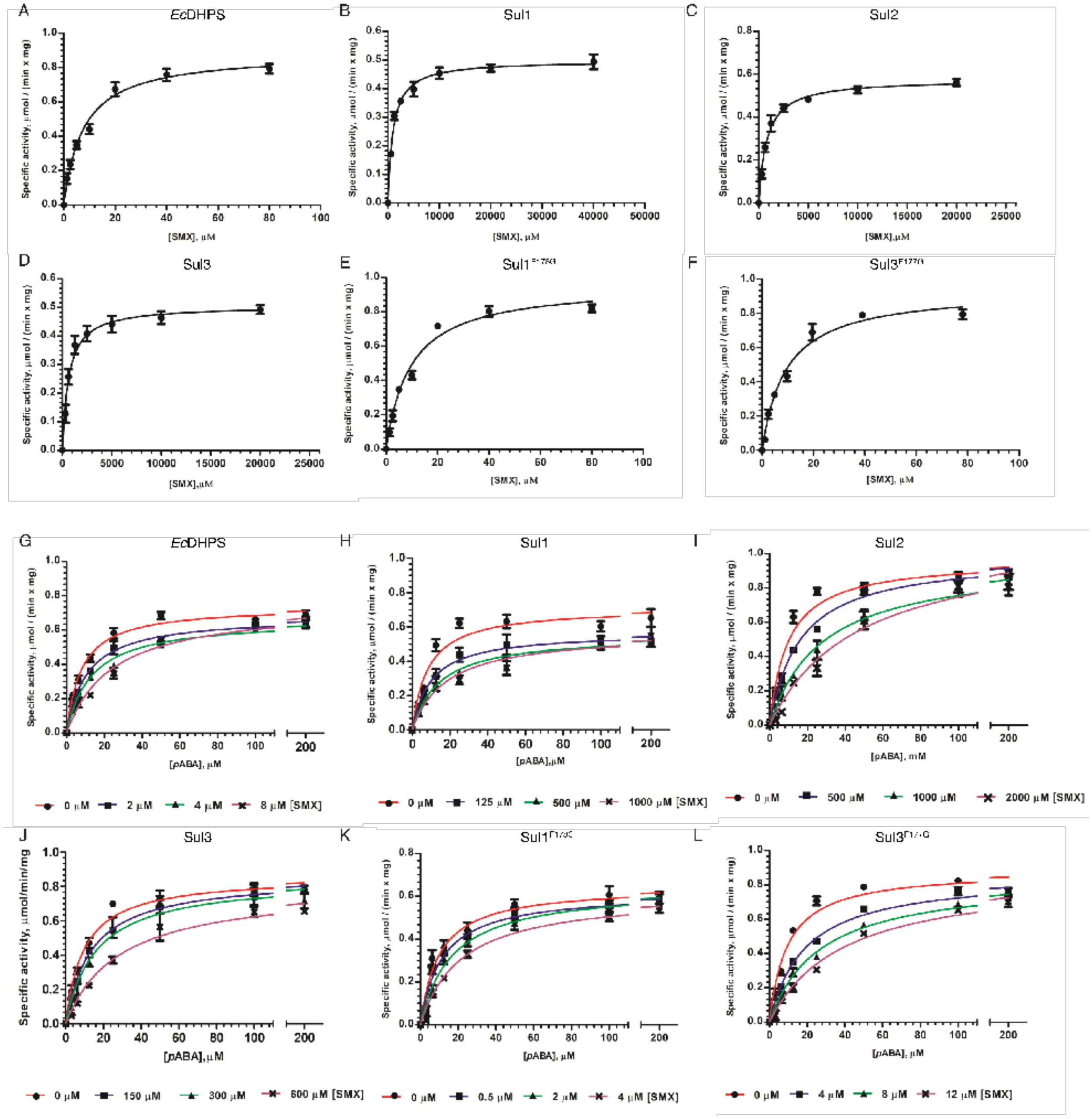


Michaelis Menten plots for A) *Ec*DHPS, B) Sul1, C) Sul2, D) Sul3, E) Sul1$^{F178G}$, F) Sul3$^{F177G}$ for the substrate SMX in the presence of saturating excess of DHPP at 200 mM. The kinetic parameters, $K_M$ and $V_{max}$ were estimated using the Michaelis-Menten equation by non-linear regression using GraphPad Prism v5.0. For calculations of inhibition constant ($K_i$) for SMX for Sul enzymes: Michaelis Menten plots for G) *Ec*DHPS, H) Sul1, I) Sul2, J) Sul3, K) Sul1$^{F178G}$, L) Sul3$^{F177G}$ with varying SMX concentrations, in the presence of saturating excess of DHPP at 200 mM. Inhibition constant $K_i$ for SMX was calculated using the formula as described in the *Experimental*. Each data point is the mean of three biological replicates for each substrate concentration. Each replicate is plotted as mean ± SD.

**Supporting Information Figure S4. Electron density maps of ligands bound to Sul and *Ec*DHPS crystal structures.**

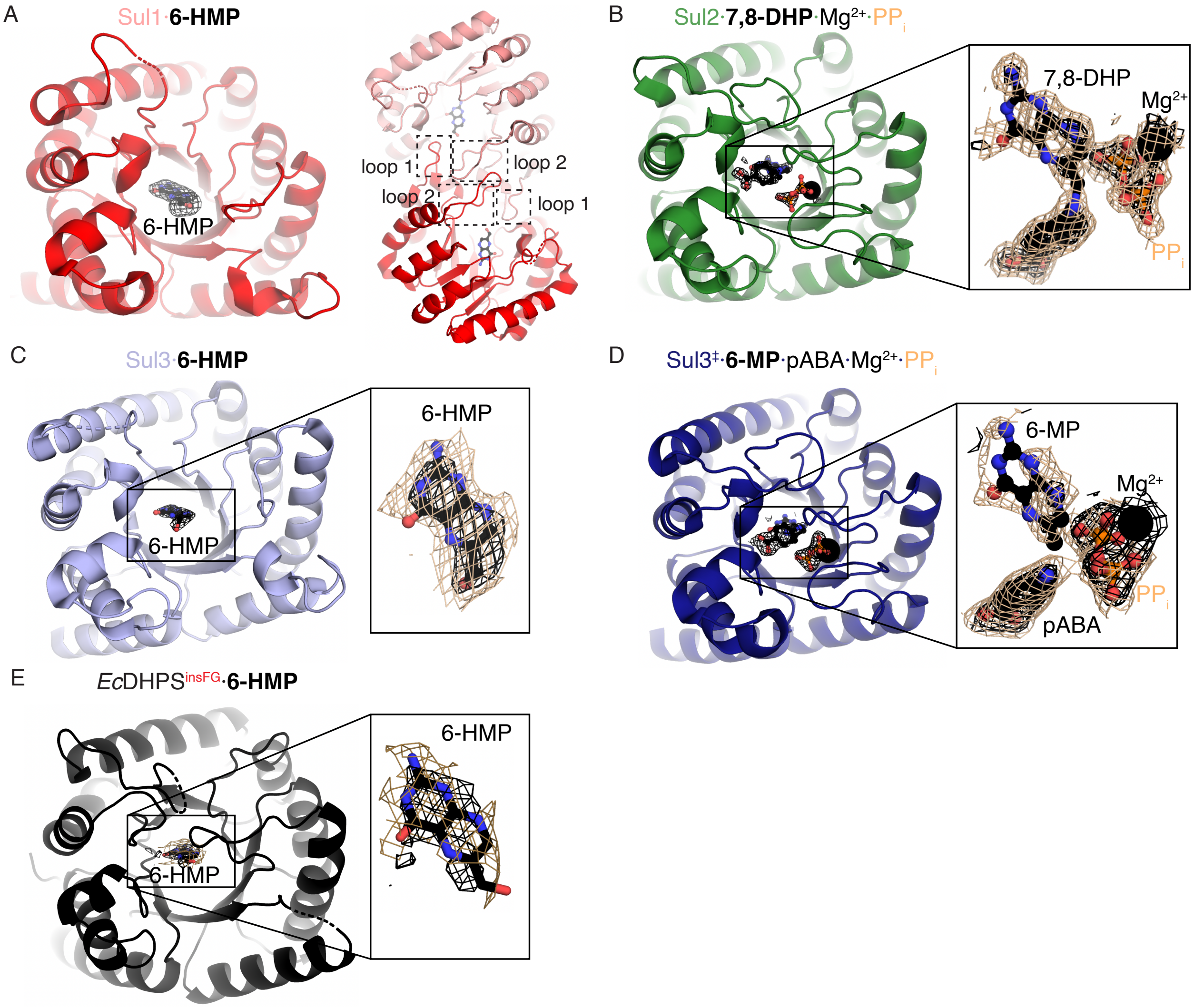


Electron density maps for ligands shown are simulated annealing omit maps (with ligand atoms and all atoms within 5 Å of the ligands deleted) contoured at 3.0σ (black) and 1.0σ (tan). A) Sul1·6-HMP complex. Right = asymmetric unit showing interditigation of loops 1 and 2 into the other subunit's active sites. B) Sul2·7,8-DHP·$Mg^{2+}$·$PP_i$ complex. C) Sul3·6-HMP complex. D) Sul3·6-MP·$Mg^{2+}$·$PP_i$ complex. E) *Ec*DHPS$^{insFG}$·6-HMP complex. Zoom-in show ligands in active sites.

**Supporting Information Figure S5. Comparison of pterin, magnesium and phosphate-interacting residues between Sul2, Sul3 and selected DHPS enzymes, and α6 Phe+loop 3 interactions.**

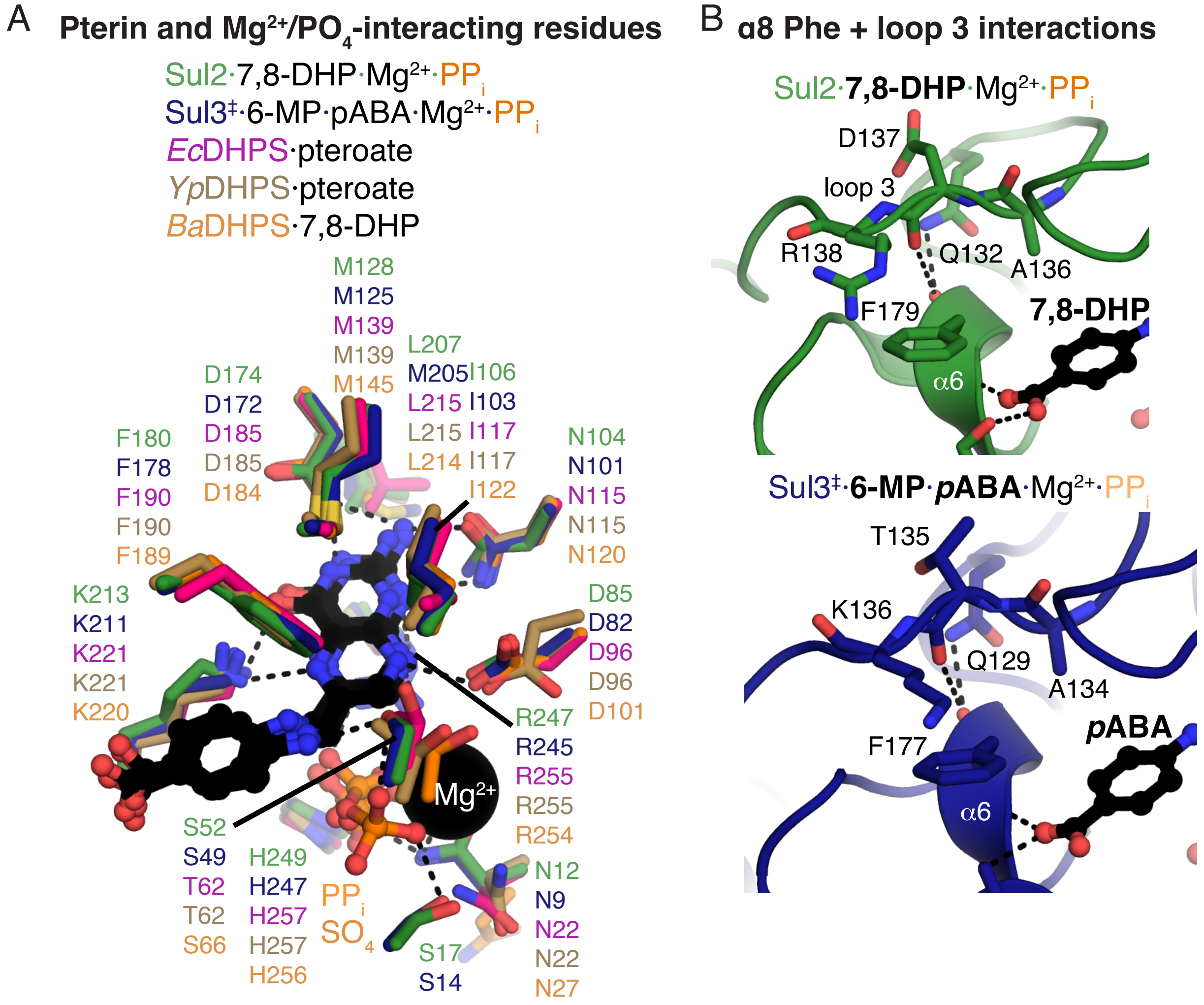


A) Shown are residues forming electrostatic, hydrogen-bonding or hydrophobic interactions with the pterin rings of bound ligands, magnesium ion, or phosphate/sulfate ions bound to the structures of Sul1, Sul3, *Ec*DHPS (PDB 5u10), *Yp*DHPS (PDB 3tyu) or *Ba*DHPS (PDB 3tya). All residues are identical except for Ser/Thr (i.e. Sul2 Ser52) which interacts with a phosphate, and a Leu/Met (i.e. Ser2 Leu207) which interacts with the pterin ring. B) Sul2 and Sul3 residues from loop 3 that interact with the α8 Phe residues. Dashes indicate hydrogen bonds.

**Supporting Information Figure S6. Multiple sequence alignment of Sul and selected DHPS enzymes.**

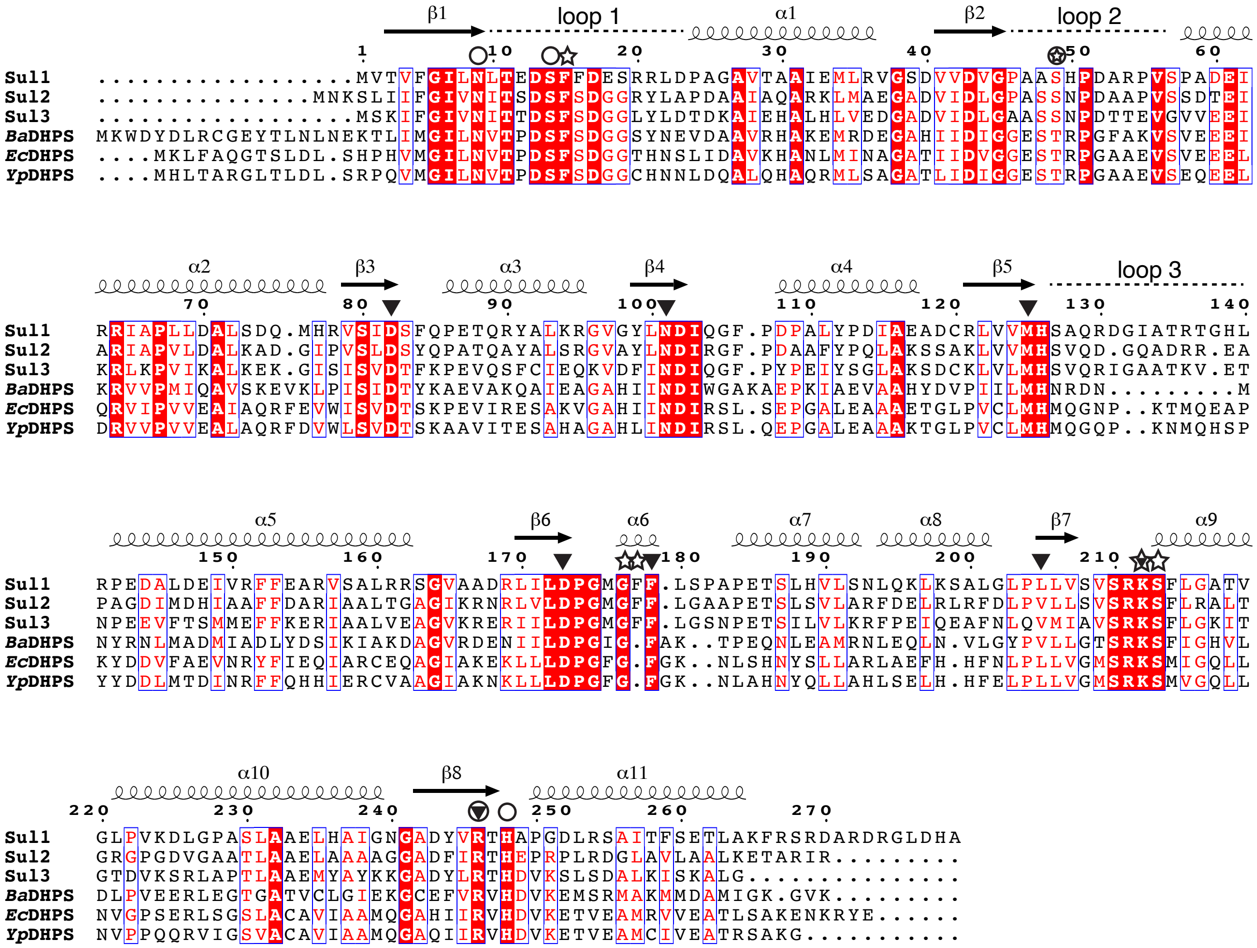


▼ interactions with pterin region

☆ interactions with *p*ABA

○ interactions with $Mg^{2+}$ or $PP_i$

Secondary structure is indicated above the alignment. Positions interacting with ligands are labeled with triangle, star or circle as indicated in the legend. Shading is according to conservation.

**Supporting Information Figure S7. Modeling of sulfa drugs into the active site of Sul2.**

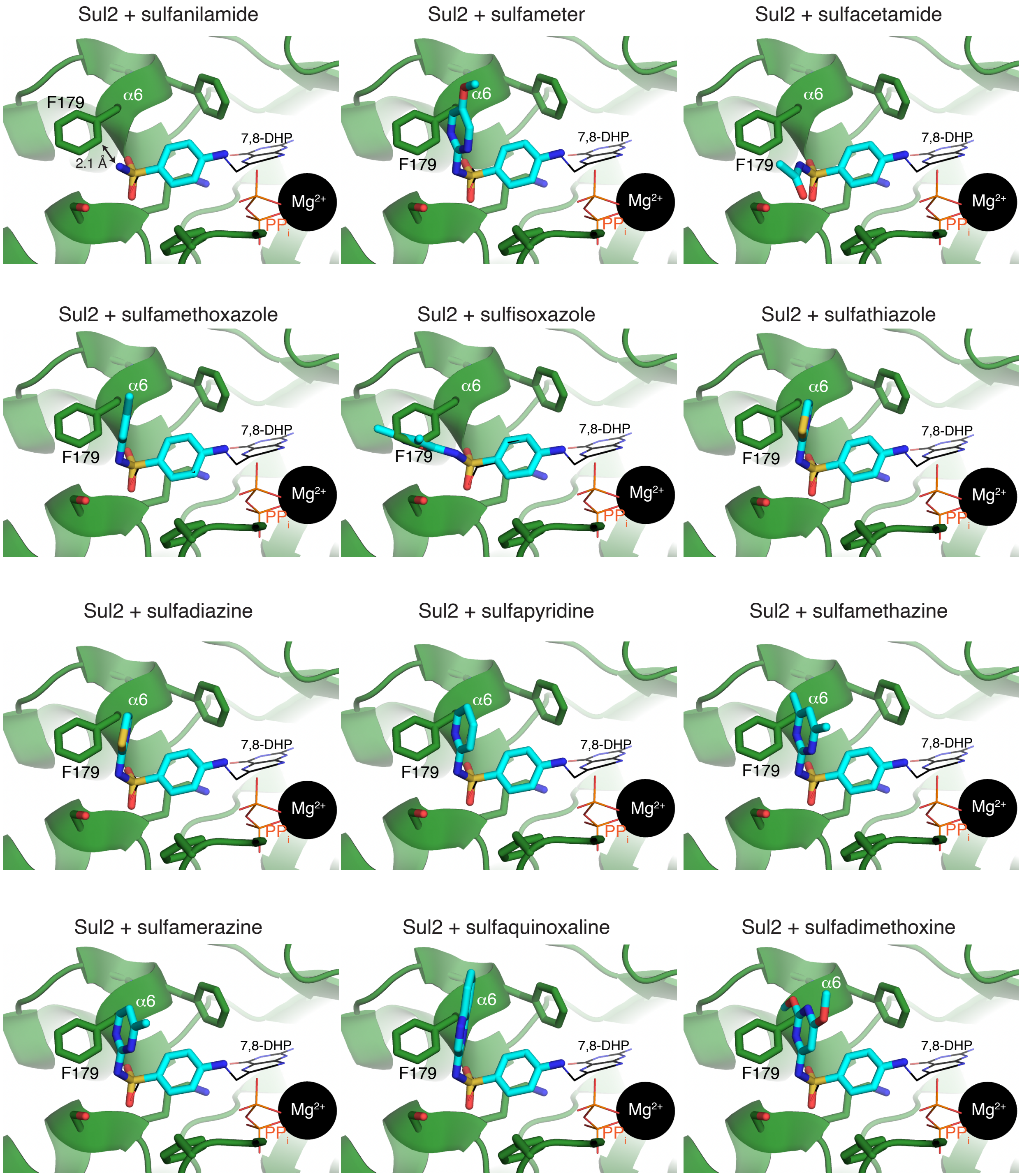


Sulfa compounds (shown in cyan sticks) were superimposed with the *p*ABA moiety observed in the structure of the Sul2·7,8-DHP·$Mg^{2+}$·$PP_i$ complex. 7,8-DHP and $PP_i$ from the Sul2·7,8-DHP·$Mg^{2+}$·$PP_i$ complex are shown in thin lines $Mg^{2+}$ as a black sphere. F179 and α8 are labeled.

**Supporting Information Figure S8. Intrinsic tryptophan fluorescence of Sul variants.**

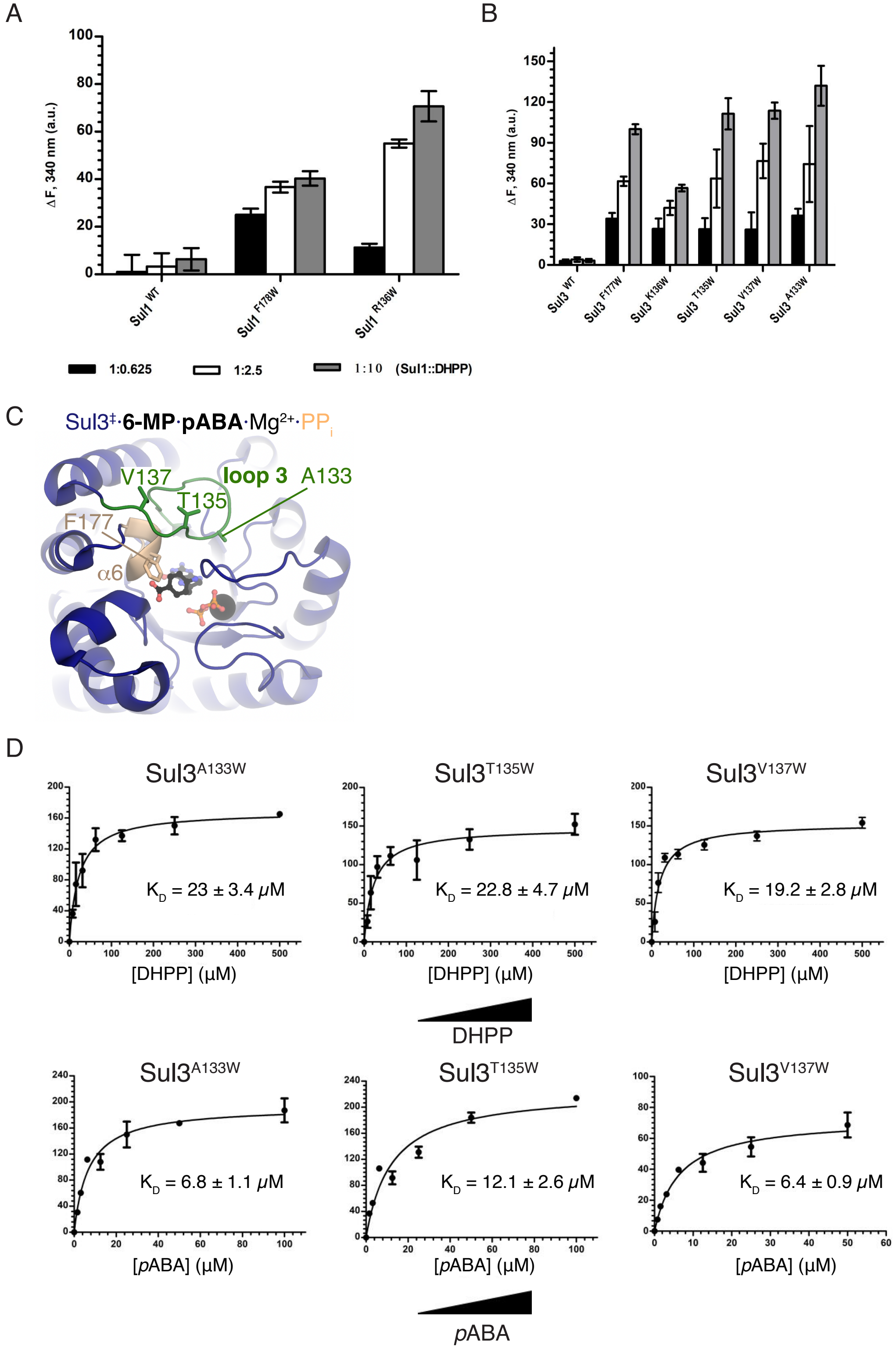


Maximum substrate-induced intrinsic tryptophan fluorescence (ITF) intensity changes monitored at 340 nm emission wavelength for (A) Sul3 and (B) Sul1 variants bound to substrate DHPP. The WT Sul3 and WT Sul1 do not have a Trp residue and therefore, show negligible baseline fluorescence intensity changes. Excitation wavelength was set as 295 nm (where the molar absorptivity of Trp is maximum while those of Phe and Tyr residues are low) and emission wavelength was monitored at 340 nm. Error bars represent ±SD. C) Location of Trp mutation sites in the structure of Sul3. D) Substrate binding affinities ($K_D$) of Sul3 Trp mutants for DHPP and *p*ABA. Substrate-induced fluorescence emission intensity change at 340 nm was plotted against substrate concentrations to derive the $K_D$ by GraphPad Prism v5.0. Triplicate readings were average, and $K_D$ reported as (±SD).

**Supporting Information Figure S9. Impact of expression of the *Ec*DHPS, Sul1, Sul2, or Sul3 enzymes or their tryptophan-substituted derivatives on the growth of the *E. coli* Δ*folP* strain under thymidine-limited conditions (MHII broth).**

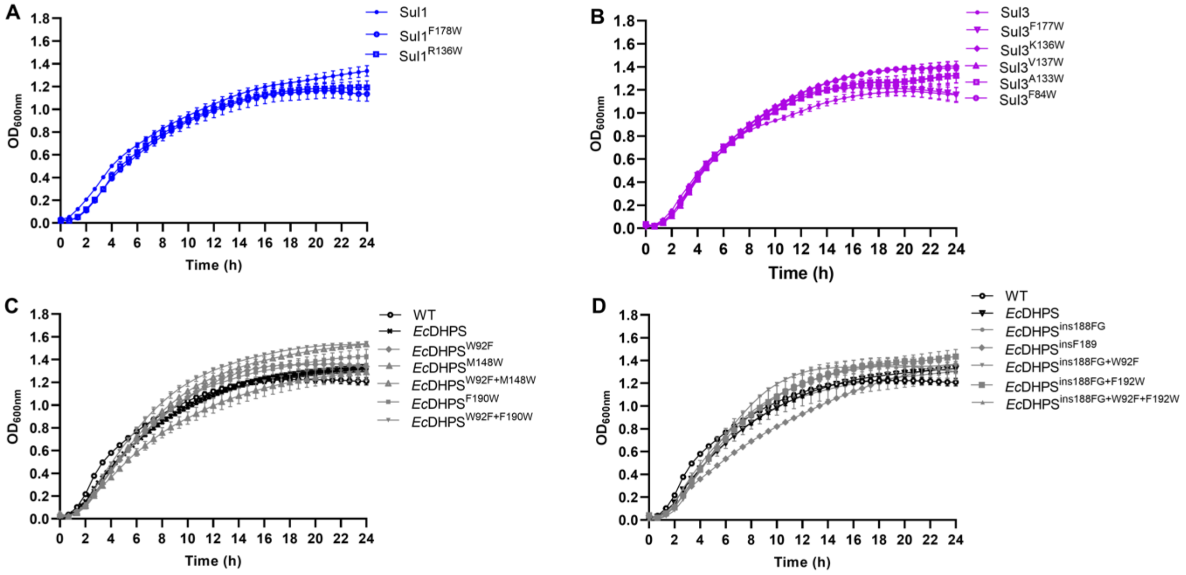


Growth curves for the Δ*folP* strain carrying pGDP2 expressing (A) the WT Sul1, Sul1$^{F178W}$, or Sul1$^{R136W}$ mutated enzymes; (B) the WT Sul3, Sul3$^{F177W}$, Sul3$^{K136W}$, Sul3$^{V137W}$, Sul1$^{A133W}$, or Sul3$^{F84W}$ mutated enzymes (C) the WT *Ec*DHPS, *Ec*DHPS$^{W92F}$, *Ec*DHPS$^{M148W}$, *Ec*DHPS$^{W92F+M148W}$, *Ec*DHPS$^{F190W}$, *Ec*DHPS$^{W92F+F190W}$ and (D) the WT *Ec*DHPS, *Ec*DHPS$^{ins188FG}$, *Ec*DHPS$^{insF189}$, *Ec*DHPS$^{ins188FG+W92F}$, *Ec*DHPS$^{ins188FG+F192W}$, *Ec*DHPS$^{ins188FG+W92F+F192W}$. The wild-type *E. coli* BW25113 (WT) strain (C and D) was included for comparison. Data is taken from at least three biological replicates performed in technical triplicate. ODs at 600 nm were baseline-corrected by subtracting background absorbance at 600 nm of the uninoculated media from the absorbance at 600 nm of the inoculated media.

**Supporting Information Figure S10. Intrinsic tryptophan fluorescence of *Ec*DHPS variants.**

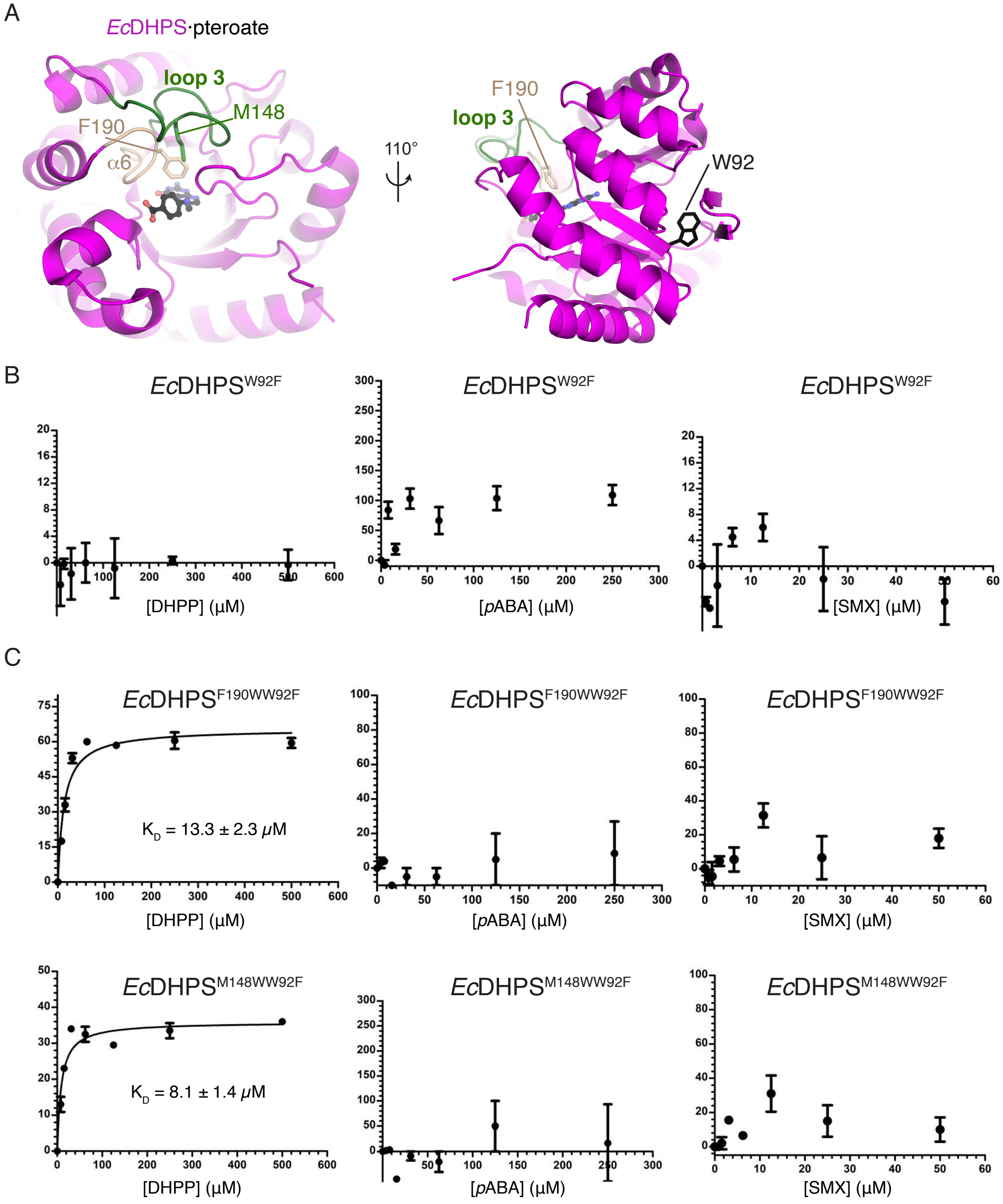


A) Location of Trp mutation sites (F190, M148) and natural W92 residue in the structure of *Ec*DHPS. B) Validation of no change in tryptophan fluorescence of *Ec*DHPS$^{W92F}$ in response to DHPP, *p*ABA or SMX. Excitation wavelength was set as 295 nm (where the molar absorptivity of Trp is maximum while those of Phe and Tyr residues are low) and emission wavelength was monitored at 340 nm. C) Substrate binding affinities ($K_D$) of *Ec*DHPS Trp mutants for DHPP, *p*ABA and SMX. Substrate-induced fluorescence emission intensity change at 340 nm (error bars represent ±SD) was plotted against substrate concentrations to derive the $K_D$ by GraphPad Prism v5.0. Triplicate readings were averaged, and $K_D$ reported as (±SD).

**Supporting Information Figure S11. Thymidine-auxotrophic phenotype of the unmarked, in-frame *folP* deletion *E. coli* mutant strain.**

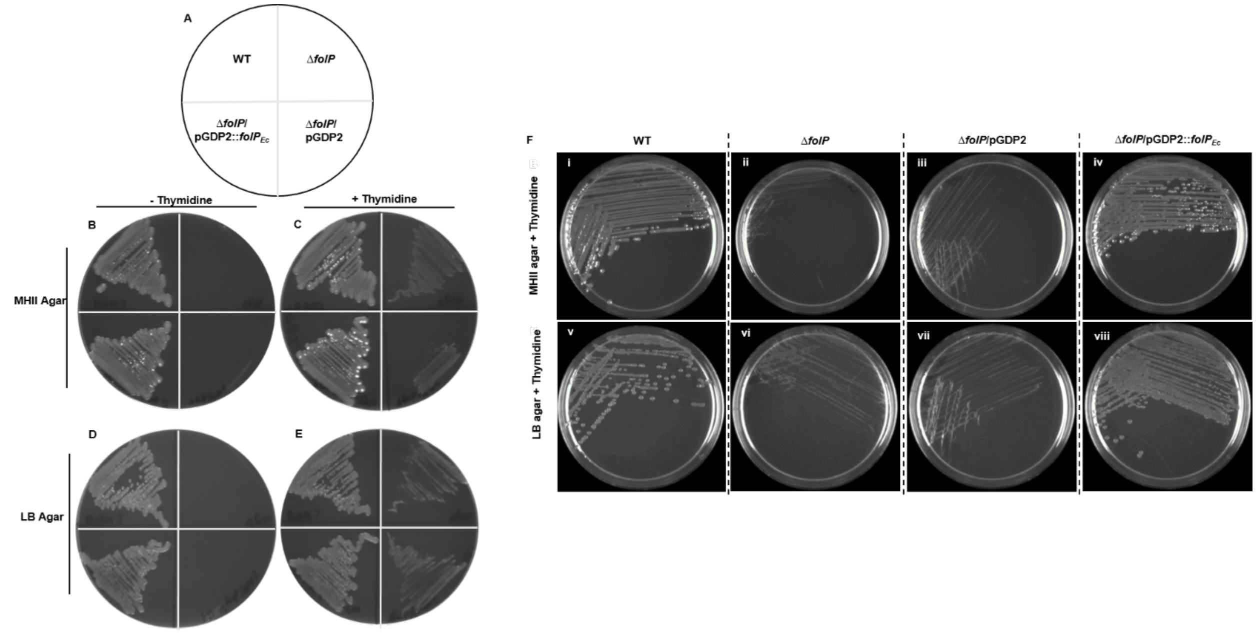


Single colonies of the parent WT *E. coli* BW25113 strain, *E. coli* Δ*folP* strain (Δ*folP* ) and its derivatives carrying the empty plasmid, pGDP2, or pGDP2 expressing the WT *folP*$_{Ec}$ gene were streaked according to the plate map shown in (A) onto either MHII agar without (B; - thymidine) or with 200 μg/mL thymidine (C; +thymidine) or LB agar without (D) or with 200 μg/ml of thymidine (E). (F) MHII agar plates (i-iv) and LB agar plates (v-viii) supplemented with 200 μg/mL thymidine that show the normal phenotype of WT *E. coli* (i and v)) and Δ*folP* strain carrying the plasmid pGDP2 expressing the WT *folP*$_{Ec}$ gene (iv and viii) and the small colony phenotypes of the Δ*folP* strain(ii and vi) and the Δ*folP* strain carrying the empty plasmid, pGDP2 (iii and vii).

**Supporting Information Figure S12. Impact of expressing the WT Sul1, Sul2, or Sul3 enzymes or their derivatives on the growth of the *E. coli* Δ*folP* strain under thymidine-limited conditions (MHII broth).**

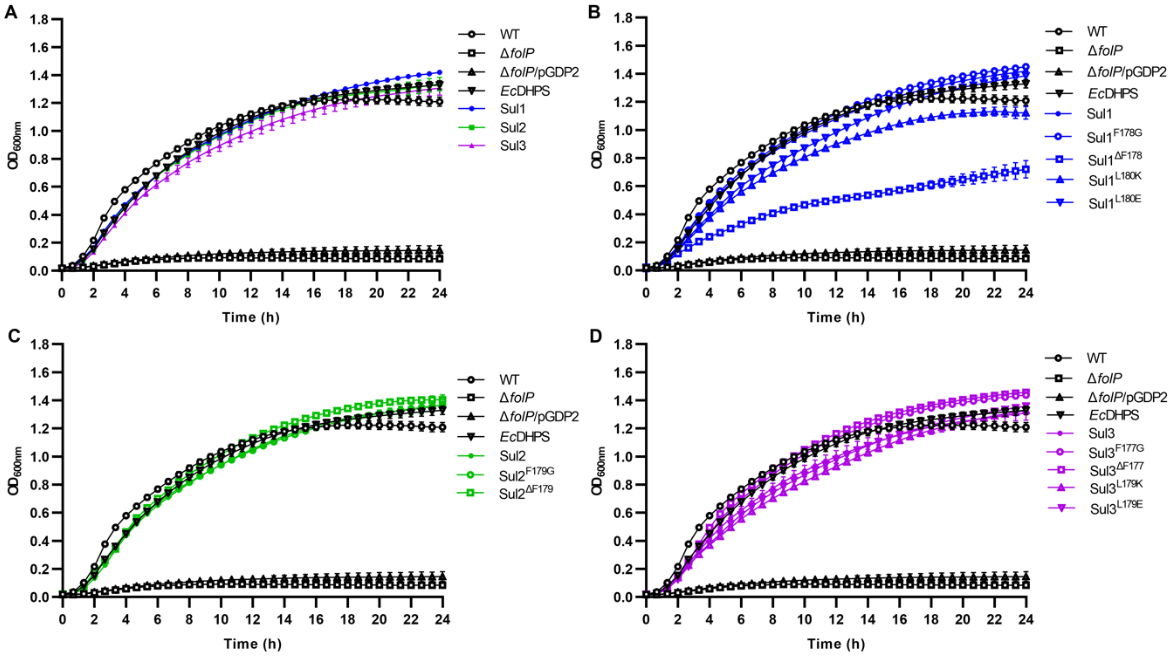


(A-D) Growth curves for wild-type *E. coli* BW25113 (WT), the plasmid-free Δ*folP* strain (Δ*folP*), empty plasmid-carrying Δ*folP* strain (Δ*folP*/pGDP2), or the Δ*folP* strain carrying pGDP2 expressing either *Ec*DHPS, or (A) the WT Sul1, Sul2, and Sul3 enzymes, or (B) the WT Sul1 enzyme and its mutated derivatives, or (C) the WT Sul2 enzyme and its mutated derivatives, or (D) the WT Sul3 enzyme and its mutated derivatives. Data is taken from at least three biological replicates performed in technical triplicate. ODs at 600 nm were baseline-corrected by subtracting background absorbance at 600 nm of the uninoculated media from the absorbance at 600 nm of the inoculated media.

**Supporting Information Figure S13. Expression of WT and mutant DHPS and Sul proteins in the *E. coli* Δ*folP* strain.**

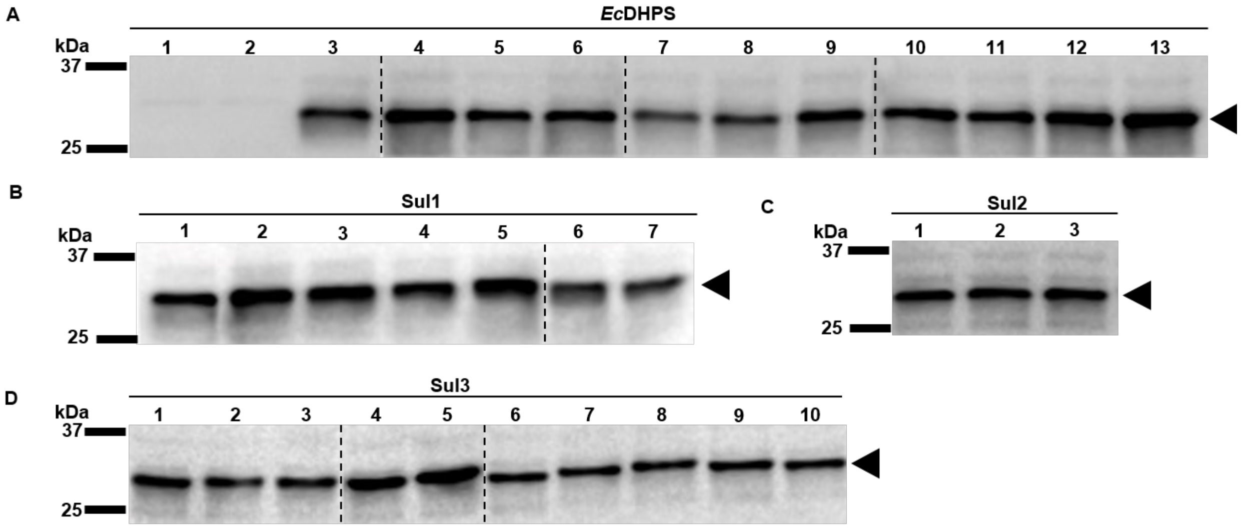


Whole-cell extracts *of* the *E. coli* Δ*folP* strain expressing plasmid pGDP2-borne, C-terminal FLAG-tagged *Ec*DHPS (A), Sul1 (B), Sul2 (C), and Sul3 (D) enzymes and their mutated derivatives were electrophoretically separated by SDS-PAGE, electroblotted, and developed with monoclonal antibodies directed against the c-terminal flag-tag. (A) Whole-cell extracts *of* the *E. coli* Δ*folP* strain with no plasmid (Lane 1), empty plasmid pGDP2 (Lane 2), or carrying plasmid pGDP2 expressing: WT *Ec*DHPS (Lane 3), *Ec*DHPS$^{W92F}$ (Lane 4), *Ec*DHPS$^{F190W}$ (Lane 5), *Ec*DHPS$^{W92F + F190W}$ (Lane 6), *Ec*DHPS$^{M148W}$ (Lane 7), *Ec*DHPS$^{W92F + M148W}$ (Lane 8), *Ec*DHPS$^{insF189}$ (Lane 9), *Ec*DHPS $^{insFG188}$ (Lane 10), *Ec*DHPS$^{ins188FG +W92F}$ (Lane 11), *Ec*DHPS$^{ins188FG + F192W}$ (Lane 12), and *Ec*DHPS$^{ins188FG + W92F + F192W}$ (Lane 13). (B) Whole-cell extracts *of* the *E. coli* Δ*folP* strain carrying plasmid pGDP2 expressing: Sul1 (Lane 1), Sul1 F178G (Lane 2), Sul1 ΔF178 (Lane 3), Sul1 L180K (Lane 4); Sul1 L180E (Lane 5); Sul1 F178W (Lane 6), and Sul1 R136W (Lane 7). (C) Whole-cell extracts of the *E. coli* Δ*folP* strain carrying plasmid pGDP2 expressing: Sul2 (Lane 1), Sul2 F178G (Lane 2), and Sul2 ΔF179G. (D) Whole-cell extracts *of* the *E. coli* Δ*folP* strain carrying plasmid pGDP2 expressing: Sul3 (Lane 1), Sul3 F177G (Lane 2), Sul3 ΔF177 (Lane 3), Sul3 L179K (Lane 4), Sul3 L179E (Lane 5), Sul3 F177W (Lane 6), Sul3 K136W (Lane 7), Sul3 V137W (Lane 8), Sul3 A133W (Lane 9), Sul3 F84W (Lane 10). The migration positions of the relevant molecular mass markers are shown on the left of each blot. Vertical dotted lines indicate non-contiguous lanes where the blots were merged. Shown are representative results from two biological replicates.

**Supporting Information Figure S14. Impact of expressing the *Ec*DHPS$^{Fins189}$ and *Ec*DHPS$^{ins188FG}$ derivatives in E. *coli* Δ*folP* strain under thymidine-limited conditions (MHII broth).**

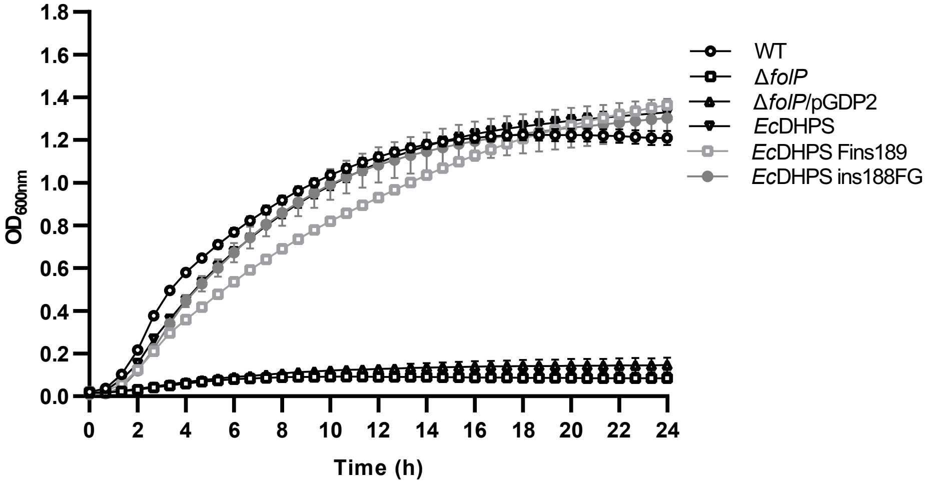

**Supporting Information Figure S15. AlphaFold2 model of *Nm*DHPS$^{insSG}$.**

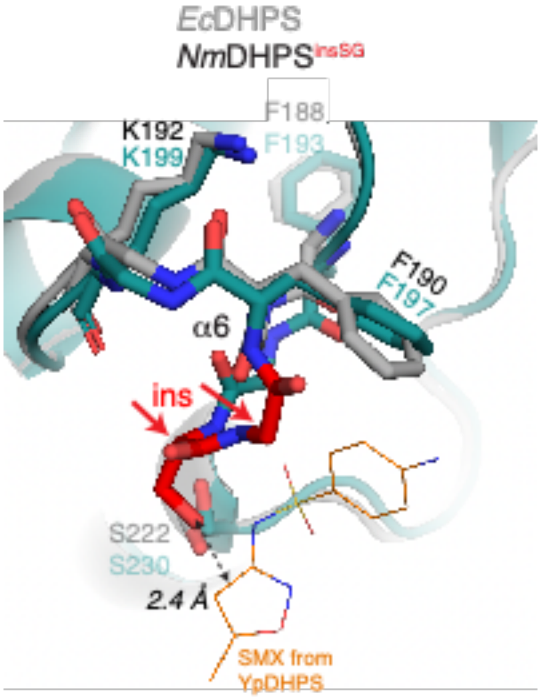


Shown is the α6 region of *Nm*DHPS with insertion SG (*Nm*DHPS$^{insSG}$) that confers sulfonamide resistance (from Fermer *et al*, *J Bacteriol.* 1995 Aug;177(16)-4669-75) overlaid with the structure of *Ec*DHPS (PDB 5U10, Dennis ML *et al*, *Chemistry* 2018 24:1922-1930) and SMX from the structure of *Yp*DHPS (PDB 3TZF, Yun MK *et al*, *Science* 2012 335:1110-1114). The *Nm*DHPS insertion Ser-Gly amino acids are coloured red and labeled.

### SUPPORTING INFORMATION TABLES

### Table S1. X-ray crystallographic statistics.

| Structure | Sul1•6-HMP | Sul2 apoenzyme | Sul2•7,8-DHP•$Mg^{2+}$•$PP_i$ |
|---|---|---|---|
| PDB code | 7S2I | 7S2J | 7S2K |
| *Data collection* | | | |
| Space group | $P4_132$ | $P2_1$ | $P2_1$ |
| Cell dimensions | | | |
| *a*, *b*, *c* (Å) | 186.6, 186.6, 186.6 | 39.8, 143.3, 85.9 | 45.7, 74.2, 67.9 |
| α, β, γ, (°) | 90, 90, 90 | 90, 90.5, 90 | 90, 96.1, 90 |
| Resolution, Å | 30.00 – 2.32 | 50.00 – 1.85 | 30.0 – 1.74 |
| $R_{merge}$[a] | 0.119 (2.135)* | 0.105 (0.764) | 0.071 (0.396) |
| $R_{pim}$[b] | 0.034 (0.520) | 0.043 (0.328) | 0.034 (0.200) |
| $CC_{1/2}$* | 0.520 | 0.912 | 0.994 |
| $I / \sigma (I)$ | 24.5 (1.04) | 19.3 (2.07) | 20.9 (2.2) |
| Completeness, % | 100 (100) | 99.4 (99.3) | 98.0 (97.4) |
| Redundancy | 13.4 (12.6) | 6.6 (6.2) | 4.6 (4.3) |
| *Refinement* | | | |
| Resolution, Å | 29.14 – 2.32 | 42.94 – 1.85 | 28.76 – 1.74 |
| No. unique reflections: working, test | 48269, 2000 | 81545, 3896 | 45534, 1999 |
| $R_{work}/R_{free}$[c] | 17.7/19.5 (28.1/29.7) | 16.2/20.7 (25.7/34.4) | 18.0/21.3 (24.0/31.4) |
| No. atoms | | | |
| Protein | 4008 | 7963 | 3911 |
| Ligands | 28 | N/A | 66 |
| Solvent | 61 | 89 | 7 |
| Water | 289 | 1168 | 394 |
| *B*-factors | | | |
| Protein | 72.3 | 35.9 | 30.5 |
| Ligands | 53.8 | N/A | 28.3 |
| Solvent | 116.5 | 52.6 | 39.7 |
| Water | 75.7 | 41.8 | 36.9 |
| R.m.s. deviations | | | |
| Bond lengths, Å | 0.005 | 0.006 | 0.006 |
| Bond angles, ° | 0.763 | 0.895 | 1.012 |

| | | | |
|---|---|---|---|
| Ramachandran plot | | | |
| Favored, % | 97.1 | 98.1 | 98.1 |
| Allowed, % | 2.9 | 1.9 | 1.9 |
| Outliers, % | 0 | 0 | 0 |

| **Structure** | **Sul3 apoenzyme** | **Sul3•6-HMP** | **Sul3•6-MP•*p*ABA•$Mg^{2+}$•$PP_i$** | ***Ec*DHPS$^{insFG}$·6-HMP** |
|---|---|---|---|---|
| PDB code | 7S2L | 7S2M | 7S2O | 7TQ1 |
| *Data collection* | | | | |
| Space group | $P6_222$ | $P3_112$ | $P3_22_1$ | I222 |
| Unit cell | | | | |
| *a*, *b*, *c* (Å) | 123.8, 123.8, 434.4 | 50.1, 50.1, 583.0 | 123.4, 123.4, 72.6 | 77.9, 84.6, 175.5 |
| α, β, γ, (°) | 90, 90, 120 | 90, 90, 120 | 90, 90, 120 | 90, 90, 90 |
| Resolution, Å | 30.00 – 2.80 | 30.00 – 2.36 | 50.0 – 2.00 | 50.0 – 2.75 |
| $R_{merge}$[a] | 0.153 (2.124)[*] | 0.093 (0.738) | 0.083 (2.091) | |
| $R_{pim}$[b] | 0.060 (0.815) | 0.050 (0.457) | 0.019 (0.527) | 0.189 (0.945) |
| | | | | 0.081 (0.435) |
| $CC_{1/2}$[*] | 0.994 (0.587 | 0.749 | 0.604 | 0.764 |
| $I$ / $\sigma$ $(I)$ | 16.38 (1.06) | 25.39 (1.0) | 41.20 (1.32) | 10.05 ( (1.04 |
| Completeness, % | 100 (100) | 98.8 (86.7) | 100 (99.9) | 95.4 (82.2) |
| Redundancy | 27.2 (19.7) | 6.4 (3.4) | 19.4 (16.1) | 6.6 (5.4) |
| *Refinement* | | | | |
| Resolution, Å | 30.00 – 2.79 | 30.04 – 2.42 | 47.0– 2.00 | 46.77 – 2.75 |
| No. unique reflections: working, test | 49650, 3687 | 26784, 1335 | 43246, 3867 | 2305, 119 |
| $R_{work}$/ $R$-$_{free}$[c] | 22.8/27.0 (40.4/40.4) | 25.0/30.0 (33.7/34.4) | 17.0/19.1 (34.5/31.3) | 23.6/28.8 (33.1/42.4) |
| No. atoms, | | | | |
| Protein | 7965 | 5900 | 2042 | 4124 |
| Ligands | N/A | 42 | 33 | 28 |
| Solvent | 91 | N/A | 61 | N/A |
| Water | 128 | 352 | 328 | 52 |

| | | | | |
|---|---|---|---|---|
| *B*-factors | | | | |
| Protein | 76.0 | 43.1 | 59.7 | 77.1 |
| Ligand | N/A | 23.3 | 51.4 | 75.3 |
| Solvent | 97.3 | N/A | 93.3 | N/A |
| Water | 63.3 | 36.1 | 73.7 | 61.5 |
| R.m.s. deviations | | | | |
| Bond lengths, Å | 0.005 | 0.002 | 0.012 | 0.004 |
| Bond angles, ° | 0.771 | 0.546 | 1.218 | 0.646 |
| Ramachandran plot | | | | |
| Favored, % | 95.0 | 94.2 | 96.2 | 95.6 |
| Allowed, % | 5.0 | 5.8 | 38 | 4.4 |
| Outliers, % | 0 | 0 | 0 | 0 |

[*]All values in brackets and $CC_{1/2}$ values refer to highest resolution shells.

[a]$R_{merge} = \Sigma_{hkl}\Sigma_j|I_{hkl.j} - \langle I_{hkl}\rangle|/\Sigma_{hkl}\Sigma_j I_{hk,j}$, where $I_{hkl,j}$ and $\langle I_{hk,j}\rangle$ are the *j*th and mean measurement of the intensity of reflection *j*.

[b]$R_{pim} = \Sigma_{hkl}\sqrt{(n/n-1)}\ \Sigma^{n}_{j=1}|I_{hkl.j} - \langle I_{hkl}\rangle|/\Sigma_{hkl}\Sigma_j I_{hk,j}$

[c]$R = \Sigma|F_p^{obs} - F_p^{calc}|/\Sigma F_p^{obs}$, where $F_p^{obs}$ and $F_p^{calc}$ are the observed and calculated structure factor amplitudes, respectively.

N/D = not applicable.

**Table S2. Sulfonamide susceptibility of the *E. coli* Δ*folP* deletion expressing the WT or mutated Sul enzymes.** [a]

| Strain | Plasmid | DHPS[c] | Growth on MHII agar[d] | MIC (µg/mL)[b] SMX | SDZ | SOZ | SPY | STZ | SMRZ | SMZ | SAA | SMT[e] | SAD | SQX[f] | SDT | MIC (µg/mL)[g] SXT (SMX-TMP) |
|---|---|---|---|---|---|---|---|---|---|---|---|---|---|---|---|---|
| WT | None | *Ec*DHPS | G | 16 | 32 | 32 | 64 | 16 | 32 | 256 | 2048 | 128 | 512 | 512 | 256 | 0.094 |
| Δ*folP* | None | None | NG | NG | NG | NG | NG | NG | NG | NG | NG | NG | NG | NG | NG | NG |
| Δ*folP* | pGDP2 | None | NG | NG | NG | NG | NG | NG | NG | NG | NG | NG | NG | NG | NG | NG |
| Δ*folP* | pGDP2 | *Ec*DHPS | G | 16 | 32 | 32 | 64 | 16 | 32 | 256 | 2048 | 128 | 512 | 512 | 256 | 0.064 |
| Δ*folP* | pGDP2 | Sul1 | G | 2048 | 4096 | 2048 | 4096 | 4096 | 4096 | 4096 | 8192 | >4096 | 2048 | >1024 | 8192 | 0.19 |
| Δ*folP* | pGDP2 | Sul1$^{F178G}$ | G | 16 | 32 | 64 | 64 | 64 | 32 | 128 | 4096 | 64 | 256 | 64 | 256 | 0.064 |
| Δ*folP* | pGDP2 | Sul1$^{F\Delta178}$ | NG | NG | NG | NG | NG | NG | NG | NG | NG | NG | NG | NG | NG | NG |
| Δ*folP* | pGDP2 | Sul1$^{L180K}$ | G | 2048 | 1024 | 2048 | 256 | 256 | 1024 | 2048 | 4096 | >4096 | 2048 | >1024 | 2048 | 0.19 |
| Δ*folP* | pGDP2 | Sul1$^{L180E}$ | G | 64 | 64 | 2048 | 64 | 32 | 64 | 256 | 1024 | 256 | 1024 | 256 | 512 | 0.023 |
| Δ*folP* | pGDP2 | Sul2 | G | 2048 | 4096 | 2048 | 4096 | 4096 | 4096 | 4096 | 8192 | >4096 | 2048 | >1024 | 8192 | 0.19 |
| Δ*folP* | pGDP2 | Sul2$^{F179G}$ | G | 32 | 64 | 128 | 128 | 128 | 128 | 512 | 4096 | 512 | 256 | 512 | 512 | 0.064 |
| Δ*folP* | pGDP2 | Sul2$^{\Delta F179}$ | G | 4 | 8 | 4 | 4 | 2 | 32 | 32 | 128 | 16 | 32 | 32 | 32 | 0.016 |
| Δ*folP* | pGDP2 | Sul3 | G | 2048 | 4096 | 2048 | 4096 | 4096 | 4096 | 4096 | 8192 | >4096 | 2048 | >1024 | 8192 | 0.19 |
| Δ*folP* | pGDP2 | Sul3$^{F177G}$ | G | 16 | 32 | 128 | 128 | 16 | 64 | 512 | 4096 | 512 | 512 | 512 | 256 | 0.064 |
| Δ*folP* | pGDP2 | Sul3$^{\Delta F177}$ | G | 2 | 8 | 16 | 16 | 2 | 16 | 32 | 256 | 32 | 32 | 16 | 8 | 0.023 |
| Δ*folP* | pGDP2 | Sul3$^{L179K}$ | G | 2048 | 4096 | 2048 | 4096 | 4096 | 4096 | 4096 | 8192 | >4096 | 2048 | >1024 | 8192 | 0.25 |
| Δ*folP* | pGDP2 | Sul3$^{L179E}$ | G | 64 | 128 | 256 | 64 | 16 | 128 | 1024 | 1024 | >4096 | 2048 | >1024 | 128 | 0.094 |

Sensitive

Resistant

[a] The sulfa susceptibility of the WT *E. coli* BW25113 (WT) strain and the *E. coli ΔfolP* strain carrying the indicated plasmids expressing wild-type *Ec*DHPS *or WT Sul1, Sul2, Sul3 or Sul1, Sul2, and Sul3* derivatives with the indicated amino acid substitutions or deletions is reported. Results for the WT *E. coli* BW25113 strain is provided for comparison purposes. Results for the plasmid-free and empty plasmid-carrying (pGDP2) *E. coli folP* deletion are provided to confirm the absence of thymidine in the MHII agar media. For all strains and drugs tested, a minimum of 3 biological replicates were performed.

[b] SMX, sulfamethoxazole; SDZ, Sulfadiazine; SOZ, sulfisoxazole; SPY, sulfapyridine; STZ, sulfathiazole; SMRZ, sulfamerazine; SMZ, sulfamethazine; SAA, sulfanilamide; SMT, sulfameter; SAD, sulfacetamide; SQX, sulfaquinoxaline; and SDT, sulfadimethoxine. Susceptibility testing for these agents was conducted using the agar dilution method and MHII agar.

[c] DHPS status of the indicated strains; *Ec*DHPS, *E. coli* WT DHPS enzyme.

[d] NG, No growth; or G, growth on MHII agar.

[e] For SMT MIC values of >4096 µg/mL indicate that the MIC is greater than 4096 µg/mL. SMT concentrations greater than 4096 µg/mL were not tested because of the drug's limited solubility in MHII agar.

[f] For SQX, MIC values of >1024 µg/mL indicate that the MIC is greater than 1024 µg/mL. SQX concentrations greater than 1024 µg/mL were not tested because of the drug's limited solubility in MHII agar.

[g] SXT, sulfamethoxazole (SMX)-trimethoprim (TMP) (1:19), values indicate TMP concentration. MICs for SXT were determined using the E-test method and MHII agar.

**Table S3. Sulfa susceptibility of the *E. coli* Δ*folP* deletion expressing the *Ec*DHPS, Sul1 or Sul3 tryptophan substitution mutant derivatives.** [a]

| Strain | Plasmid | DHPS[c] | Growth on MHII agar[d] | MIC (µg/mL)[b] SMX | SDZ | SOZ |
|---|---|---|---|---|---|---|
| WT | None | *Ec*DHPS | G | 16 | 32 | 32 |
| Δ*folP* | pGDP2 | *Ec*DHPS | G | 16 | 32 | 32 |
| Δ*folP* | pGDP2 | *Ec*DHPS$^{W92F}$ | G | 32 | 32 | 64 |
| Δ*folP* | pGDP2 | *Ec*DHPS$^{F190W}$ | G | 8 | 16 | 16 |
| Δ*folP* | pGDP2 | *Ec*DHPS$^{W92F\ F190W}$ | G | 16 | 16 | 16 |
| Δ*folP* | pGDP2 | *Ec*DHPS$^{M148W}$ | G | 8 | 16 | 16 |
| Δ*folP* | pGDP2 | *Ec*DHPS$^{W92F\ M148W}$ | G | 4 | 16 | 8 |
| Δ*folP* | pGDP2 | *Ec*DHPS$^{ins188FG}$ | G | 32 | 128 | 64 |
| Δ*folP* | pGDP2 | *Ec*DHPS$^{ins188FG\ W92F}$ | G | 64 | 256 | 128 |
| Δ*folP* | pGDP2 | *Ec*DHPS$^{ins188FG\ F192W}$ | G | 16 | 64 | 64 |
| Δ*folP* | pGDP2 | *Ec*DHPS$^{ins188FG\ W92F\ F192W}$ | G | 32 | 64 | 64 |
| Δ*folP* | pGDP2 | Sul1 | G | 2048 | 4096 | 2048 |
| Δ*folP* | pGDP2 | Sul1$^{F178W}$ | G | 2048 | 4096 | 4096 |
| Δ*folP* | pGDP2 | Sul1$^{R136W}$ | G | 2048 | 4096 | 4096 |
| Δ*folP* | pGDP2 | Sul3 | G | 2048 | 4096 | 2048 |
| Δ*folP* | pGDP2 | Sul3$^{F177W}$ | G | 2048 | 4096 | 4096 |
| Δ*folP* | pGDP2 | Sul3$^{K136W}$ | G | 2048 | 4096 | 4096 |
| Δ*folP* | pGDP2 | Sul3$^{V137W}$ | G | 2048 | 4096 | 4096 |
| Δ*folP* | pGDP2 | Sul3$^{A33W}$ | G | 2048 | 4096 | 4096 |
| Δ*folP* | pGDP2 | Sul3$^{F84W}$ | G | 2048 | 4096 | 4096 |

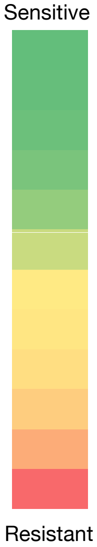


[a] The sulfa susceptibility of the WT *E. coli* BW25113 (WT) strain and the *E. coli ΔfolP* strain carrying the indicated plasmids expressing wild-type *Ec*DHPS, *Ec*DHPS insFG188, Sul1, Sul3 or their tryptophan substitution mutant derivatives with the indicated amino acid is reported. Results for the *E. coli* BW25113 strain (WT) is provided for comparison purposes. For all strains and drugs tested, a minimum of 3 biological replicates were performed using the agar dilution method and MHII agar.

[b] SMX, sulfamethoxazole; SDZ. Sulfadiazine; SOZ, sulfisoxazole.

[c] DHPS status of the indicated strains ; *Ec*DHPS, *E. coli* WT DHPS enzyme

[d] NG, No growth; or G, growth on MHII agar.

**Table S4. Sulfa susceptibility[a] of the sulfamethoxazole-resistant DHPS mutants, *Ec*DHPS$^{ins188FG}$ of *E. coli* BW25113 selected following a 7-day sulfanilamide exposure and susceptibility of the Δ*folP* strain expressing plasmid-borne *Ec*DHPS$^{ins188FG}$.**

| | | | MIC (µg/mL)[b] | | | | | | | | | | | | MIC (µg/mL)[e] |
|---|---|---|---|---|---|---|---|---|---|---|---|---|---|---|---|
| Strain | Plasmid | DHPS | SMX | SDZ | SOZ | SPY | STZ | SMRZ | SMZ | SAA | SMT[c] | SAD | SQX[d] | SDT | SXT (SMX-TMP)[e] |
| WT | None | *Ec*DHPS | 16 | 32 | 32 | 64 | 16 | 32 | 256 | 2048 | 128 | 512 | 512 | 256 | 0.094 |
| Δ*folP* | pGDP2 | *Ec*DHPS | 16 | 32 | 32 | 64 | 16 | 32 | 256 | 2048 | 128 | 512 | 512 | 256 | 0.064 |
| SMX$^{R}$-1.4 | None | *Ec*DHPS$^{ins188FG}$ | 256 | 512 | 256 | ND[f] | ND | ND | ND | 4096 | ND | ND | ND | ND | ND |
| SMX$^{R}$-3.4 | None | *Ec*DHPS$^{ins188FG}$ | 256 | 512 | 512 | ND | ND | ND | ND | 4096 | ND | ND | ND | ND | ND |
| SMX$^{R}$-4.4 | None | *Ec*DHPS$^{ins188FG}$ | 256 | 512 | 512 | ND | ND | ND | ND | 4096 | ND | ND | ND | ND | ND |
| Δ*folP* | pGDP2 | *Ec*DHPS$^{ins188FG}$ | 32 | 128 | 64 | 256 | 64 | 256 | 512 | 4096 | 1024 | 1024 | >1024 | 512 | 0.125 |
| Δ*folP* | pGDP2 | *Ec*DHPS$^{Fins189}$ | 8 | 16 | 16 | 64 | 8 | 8 | 64 | 256 | 64 | 128 | 256 | 64 | 0.047 |

Sensitive

Resistant

[a] Wild-type *E. coli* BW25113 was exposed to sulfanilamide (half the MIC; 1024 µg/mL) over 7 days and mutants resistant to 256 µg/mL of sulfamethoxazole (SMX) were selected. Results for the *E. coli* BW25113 strain (WT) and for the three SMX-resistant mutants (SMX$^{R}$) harboring an FG insertion at position 188 of the WT *Ec*DHPS coding region are reported. The antimicrobial susceptibility of *E. coli* Δ*folP* strain carrying the plasmid pGDP2 expressing (WT) *Ec*DHPS or EcDHPS derivatives with the indicated amino acid insertions are also reported. For all strains and drugs tested, a minimum of 3 biological replicates were performed using the agar dilution method and MHII agar.

[b] SMX, sulfamethoxazole; SDZ, Sulfadiazine; SOZ, sulfisoxazole; SPY, sulfapyridine; STZ, sulfathiazole; SMRZ, sulfamerazine; SMZ, sulfamethazine; SAA, sulfanilamide; SMT, sulfameter; SAD, sulfacetamide; SQX, sulfaquinoxaline; and SDT, sulfadimethoxine.

[c] For SMT MIC values of >4096 µg/mL indicate that the MIC is greater than 4096 µg/mL. SMT concentrations greater than 4096 µg/mL were not tested because of the drug's limited solubility in MHII agar.

[d] For SQX, MIC values of >1024 µg/mL indicate that the MIC is greater than 1024 µg/mL. SQX concentrations greater than 1024 µg/mL were not tested because of the drug's limited solubility in MHII agar.

[e] SXT, sulfamethoxazole (SMX)-trimethoprim (TMP) (1:19), values indicate TMP concentration. MICs for SXT were determined using the E-test method and MHII agar.

[f] ND, not determined.

### Table S5. Bacterial strains and plasmids used in this study.

| Strain | Relevant Genotype | Source |
|---|---|---|
| ***E. coli*** | | |
| DH5α | *φ80ΔlacZΔM15 endA1 recA1 hsdR17(rK -mK+) supE44 thi-1 gyrA96 relA1 F- Δ(lacZYA-argF) U169* | Ausubel FM, Brent R, Kingston RE, Moore DD, Seidman JG, et al. (1992) Short protocols in molecular biology, 2nd ed. New York: John Wiley & Sons, Inc. |
| BW25113 | *E. coli* K-12 BW25113 wild type: Δ(*araD-araB*)*567* Δ*lacZ4787*(::*rrnB-3*) *rph-1* Δ(*rhaD-rhaB*)*568 hsdR514* | Keio Collection |
| Δ*folP* | BW25113 carrying an unmarked, in-frame *folP* gene deletion | This study |
| BL21(DE3) Gold | *E. coli B F- dcm+ The ompT hsdS*($r_B$-$m_B$-) *gal λ (DE3) endA Tet*$^r$ | Agilent |
| SMX$^{R}$-14.1 | Sulfa-resistant BW25113 derivative carrying a *folP*$_{T62K}$ mutation that was selected following a 7-day exposure to 1024 µg/mL of sulfanilamide | This study |
| SMX$^{R}$-20.1 | Sulfa-resistant BW25113 derivative carrying a *folP*$_{T62A}$ mutation that was selected following a 7-day exposure to 1024 µg/mL of sulfanilamide | This study |
| SMX$^{R}$-22.1 | Sulfa-resistant BW25113 derivative carrying a *folP*$_{T62K}$ mutation that was selected following a 7-day exposure to 1024 µg/mL of sulfanilamide | This study |
| SMX$^{R}$-24.1 | Sulfa-resistant BW25113 derivative carrying a *folP*$_{T62A}$ mutation that was selected following a 7-day exposure to 1024 µg/mL of sulfanilamide | This study |
| SMX$^{R}$-26.1 | Sulfa-resistant BW25113 derivative carrying a *folP*$_{T62A}$ mutation that was selected following a 7-day exposure to 1024 µg/mL of sulfanilamide | This study |
| SMX$^{R}$-12.2 | Sulfa-resistant BW25113 derivative carrying a *folP*$_{T62A}$ mutation that was selected following a 7-day exposure to 1024 µg/mL of sulfanilamide | This study |
| SMX$^{R}$-42.2 | Sulfa-resistant BW25113 derivative carrying a *folP*$_{T62A}$ mutation that was selected following a 7-day exposure to 1024 µg/mL of sulfanilamide | This study |
| SMX$^{R}$-46.2 | Sulfa-resistant BW25113 derivative carrying a *folP*$_{T62A}$ mutation that was selected following a 7-day exposure to 1024 µg/mL of sulfanilamide | This study |
| SMX$^{R}$-48.2 | Sulfa-resistant BW25113 derivative carrying a *folP*$_{T62A}$ mutation that was selected following a 7-day exposure to 1024 µg/mL of sulfanilamide | This study |
| SMX$^{R}$-60.2 | Sulfa-resistant BW25113 derivative carrying a *folP*$_{T62A}$ mutation that was selected following a 7-day exposure to 1024 µg/mL of sulfanilamide | This study |
| SMX$^{R}$-12.3 | Sulfa-resistant BW25113 derivative carrying a *folP*$_{T62A}$ mutation that was selected following a 7-day exposure to 1024 µg/mL of sulfanilamide | This study |
| SMX$^{R}$-16.3 | Sulfa-resistant BW25113 derivative carrying a *folP*$_{T62A}$ mutation that was selected following a 7-day exposure to 1024 µg/mL of sulfanilamide | This study |
| SMX$^{R}$-18.3 | Sulfa-resistant BW25113 derivative carrying a *folP*$_{T62A}$ mutation that was selected following a 7-day exposure to 1024 µg/mL of sulfanilamide | This study |
| SMX$^{R}$-20.3 | Sulfa-resistant BW25113 derivative carrying a *folP*$_{T62A}$ mutation that was selected following a 7-day exposure to 1024 µg/mL of sulfanilamide | This study |

| | | |
|---|---|---|
| $SMX^R$-22.3 | Sulfa-resistant BW25113 derivative carrying a *folP*$_{T62A}$ mutation that was selected following a 7-day exposure to 1024 µg/mL of sulfanilamide | This study |
| $SMX^R$-22.3 | Sulfa-resistant BW25113 derivative carrying a *folP*$_{T62A}$ mutation that was selected following a 7-day exposure to 1024 µg/mL of sulfanilamide | This study |
| $SMX^R$-22.3 | Sulfa-resistant BW25113 derivative carrying a *folP*$_{T62A}$ mutation that was selected following a 7-day exposure to 1024 µg/mL of sulfanilamide | This study |
| $SMX^R$-1.4 | Sulfa-resistant BW25113 derivative carrying a *folP* $_{Gly187\_Phe188insPheGly}$ mutation that was selected following a 7-day exposure to 1024 µg/mL of sulfanilamide | This study |
| $SMX^R$-2.4 | Sulfa-resistant BW25113 derivative with no mutation in *folP* that was selected following a 7-day exposure to 1024 µg/mL of sulfanilamide | This study |
| $SMX^R$-3.4 | Sulfa-resistant BW25113 derivative carrying a *folP* $_{Gly187\_Phe188insPheGly}$ mutation that was selected following a 7-day exposure to 1024 µg/mL of sulfanilamide | This study |
| $SMX^R$-4.4 | Sulfa-resistant BW25113 derivative carrying a *folP* $_{Gly187\_Phe188insPheGly}$ mutation that was selected following a 7-day exposure to 1024 µg/mL of sulfanilamide | This study |
| $SMX^R$-5.4 | Sulfa-resistant BW25113 derivative with no mutation in *folP* that was selected following a 7-day exposure to 1024 µg/mL of sulfanilamide | This study |
| **Plasmids** | | |
| pUC19 | *E. coli* gene expression vector, $Ap^R$ | PMID:6323249 |
| pMJF200 | pUC19:: Δ*folP* upstream fragment | This study |
| pMJF201 | pUC19:: Δ*folP* downstream fragment | This study |
| pMJF202 | pUC19:: Δ*folP* | This study |
| pKOV | *repA101*(Ts) *sacB* $Cm^r$ ; Gene replacement vector | PMID:9335267 |
| pMJF203 | pKOV::Δ*folP* | This study |
| pMCSG53 | N-terminal $His_6$-tag expression vector; $Ap^R$ | PMID:24057978 |
| pNIC-CH | C-terminal $His_6$-tag expression vector; $Km^R$ | |
| pGDP2 | *E. coli* gene expression vector; $Km^R$ | PMID:28017602 |
| pMJF204 | pGDP2::*sul1*$_{WT}$-FLAG | This study |
| pMJF205 | pGDP2::*sul1*$_{F178G}$-FLAG | This study |
| pMJF206 | pGDP2::*sul1*$_{\Delta178}$-FLAG | This study |
| pMJF207 | pGDP2::*sul1*$_{L180K}$-FLAG | This study |
| pMJF208 | pGDP2::*sul1*$_{L180E}$-FLAG | This study |
| pMJF209 | pGDP2::*sul1*$_{F178W}$-FLAG | This study |

| | | |
|---|---|---|
| pMJF210 | pGDP2::*sul1*$_{R136W}$-FLAG | This study |
| pMJF211 | pGDP2::*sul2*$_{WT}$-FLAG | This study |
| pMJF212 | pGDP2::*sul2*$_{F179G}$-FLAG | This study |
| pMJF213 | pGDP2::*sul2*$_{\Delta F179}$-FLAG | This study |
| pMJF214 | pGDP2::*sul3*$_{WT}$-FLAG | This study |
| pMJF215 | pGDP2::*sul3*$_{F177G}$-FLAG | This study |
| pMJF216 | pGDP2::*sul3*$_{\Delta F177}$-FLAG | This study |
| pMJF217 | pGDP2::*sul3*$_{L179K}$-FLAG | This study |
| pMJF218 | pGDP2::*sul3*$_{L179E}$-FLAG | This study |
| pMJF219 | pGDP2::*sul3*$_{F177W}$-FLAG | This study |
| pMJF220 | pGDP2::*sul3*$_{K136W}$-FLAG | This study |
| pMJF213 | pGDP2::*sul3*$_{V137W}$-FLAG | This study |
| pMJF221 | pGDP2::*sul3*$_{A33W}$-FLAG | This study |
| pMJF222 | pGDP2::*sul3*$_{F84W}$-FLAG | This study |
| pMJF223 | pGDP2::*folP*$_{Ec}$-FLAG | This study |
| pMJF224 | pGDP2::*folP* $_{Gly187\_Phe188insPheGly}$ –FLAG | This study |
| pMJF225 | pGDP2::*folP* $_{Gly189\_Phe190insPhe}$ –FLAG | This study |
| pMJF226 | pGDP2::*folP*$_{W92F}$ –FLAG | This study |
| pMJF227 | pGDP2::*folP*$_{F190W}$ –FLAG | This study |
| pMJF228 | pGDP2::*folP*$_{W92F+F190W}$ –FLAG | This study |
| pMJF229 | pGDP2::*folP*$_{F148W}$ –FLAG | This study |
| pMJF230 | pGDP2::*folP*$_{W92F+M148W}$ –FLAG | This study |
| pMJF231 | pGDP2::*folP*$_{W92F+M148W}$ –FLAG | This study |
| pMJF232 | pGDP2::*folP* $_{W92F+Gly189\_Phe190insPhe}$ –FLAG | This study |
| pMJF233 | pGDP2::*folP* $_{W92F+Gly189\_Phe190insPhe\ +F192W}$ –FLAG | This study |
| pMJF234 | pGDP2::*folP* $_{W92F+Gly189\_Phe190insPhe+F192W}$ –FLAG | This study |

Note: $Ap^R$ ampicillin resistant; $Cm^R$ chloramphenicol resistant; $Km^R$, kanamycin resistant; $SMX^R$, sulfamethoxazole resistant; WT, wild type; FLAG, FLAG-tag.

### Table S6. Primers used in this study.

| Primer Name | Sequence (5'-3') |
|---|---|
| Δ*folP* Up For | GACTAAGCTTCAGGTTGTGGTCGGCTTGCC<br>HindIII |
| Δ*folP* Up Rev | GACTTCTAGAGGCAAAGAGTTTCATGATGTTATCCCTGG<br>XbaI |
| Δ*folP* Dn For | GACTTCTAGAGTGGTGGAAGCCACTCTGTCTGCA<br>XbaI |
| Δ*folP* Dn Rev | GACTGGATCCAGCCAGTTATCTAACGCTTT<br>BamHI |
| Δ*folP* For | GACTGCGGCCGCCAGGTTGTGGTCGGCTTGCC<br>NotI |
| *folP* For | CGACGCACCGCAGATTGATGACCTG |
| *folP* Rev | CCAGTGCTGACTCCAGCATATAGCC |
| *folP*$_{ins188FG}$ For | GACTTCTAGATTAACTTTAAGAAGGAGATATACATGAAACTCTTTGCC<br>XbaI<br>CAGGGTAC |
| *folP*$_{ins188FG}$ Rev | GACTAAGCTT**TTA***CTTGTCGTCATCGTCTTTGTAGTC*CTCATAGCGTT<br>HindIII<br>TGTTTTCCTTT |

Note: Underlined sequence demarcate restriction enzyme cut sites; bolded sequence codes for a stop codon; italicized sequence codes for the FLAG-tag.
